# Monitoring the Coating of Single DNA Origami Nanostructures with a Molecular Fluorescence Lifetime Sensor

**DOI:** 10.1101/2024.10.28.620667

**Authors:** Michael Scheckenbach, Gereon Andreas Brüggenthies, Tim Schröder, Karina Betuker, Lea Wassermann, Philip Tinnefeld, Amelie Heuer-Jungemann, Viktorija Glembockyte

## Abstract

The high functionality of DNA nanostructures makes them a promising tool for biomedical applications, their intrinsic instability under application-relevant conditions, still remains challenging. Protective coating of DNA nanostructures with materials like silica or cationic polymers has evolved as a simple, yet powerful strategy to improve their stability even under extreme conditions. While over time, various materials and protocols have been developed, the characterization and quality assessment of the coating is either time consuming, highly invasive or lacks detailed insights on single nanostructures. Here, we introduce a cyanine dye based molecular sensor designed to non-invasively probe the coating of DNA origami by either a cationic polymer or by silica, in real-time and on a single nanostructure level. The cyanine dye reports changes in its local environment upon coating via increased fluorescence lifetime induced by steric restriction and water exclusion. Exploiting the addressability of DNA origami, the molecular sensor can be placed at selected positions to probe the coating layer with nanometer precision. We demonstrate the reversibility of the sensor and use it to study the stability of the different coatings in degrading conditions. To showcase the potential for correlative studies, we combine the molecular fluorescence lifetime sensor with DNA PAINT super-resolution imaging to investigate coating and structural integrity as well as preserved addressability of DNA nanostructures. The reported sensor presents a valuable tool to probe the coating of DNA nanodevices in complex biochemical environments in real-time and at the single nanosensor level and aids the development of novel stabilization strategies.

## INTRODUCTION

DNA nanotechnology and, in particular, the DNA origami technique have advanced rapidly in the last decades and have reached an unprecedent level of complexity and functionality at the nanoscale.^1^ While the easy design of DNA origami and the spatial control of chemical modifications with base pair precision have opened up a plethora of potential applications in different fields, such as, plasmonics, biosensing, drug delivery or nanorobotics, the intrinsic instability towards external factors often remains a bottleneck.^2–11^

While DNA itself is susceptible to degradation by nucleases, employing it as building material in closely-packed self-assemblies necessitates the presence of specific cations like Mg^2+^ to compensate the anionic charge of the phosphate backbone, limiting the application window of DNA nanostructures to mild conditions (buffer conditions, specific ion concentrations, mild pH values, low temperatures and irradiation) and to generally short device lifecycles.^12–16^ Consequently, multiple strategies have been developed to increase the stability of DNA self-assemblies, for example, by optimizing the design,^17^ by modifying the ends of staple strands with polymers such as polyethylene glycol (PEG),^18, 19^ by covalently connecting neighboring thymine bases via UV light induced cross-linking ^20–22^ or by replacing defective staple strands via dynamic self-repair.^23^ A simple, yet extremely effective strategy to increase the stability of functional DNA nanodevices is protective coating via electrostatic interactions between the negatively charged phosphate backbone of the DNA and a positively charged coating agent (Figure 1a). Using the cationic silica precursor *N*-[3-(trimethoxysilyl)propyl]-*N,N,N*-trimethylammonium chloride (TMAPS) in a mixture with the classic precursor tetraethyl orthosilicate (TEOS) enables the growth of nanometers thick silica layers on DNA origami, either in solution or immobilized on various substrate surfaces.^24–26^ In a similar approach, DNA nanostructures can be coated with cationic polymers, such as poly-L-lysine polyethylene glycol block copolymer (PLL-PEG), leading to a sub-nanometer thick shell.^27–29^ While the silicification of DNA nanostructures leads to highly increased mechanical, chemical, biological and thermal stability ^25, 30, 31^, the coating with PLL-PEG increases the chemical stability in low-salt and serum conditions and it can be reversed by the addition of an anionic polymer such as dextran sulfate.^27, 28^ Despite coating with a thick silica shell or a PLL-PEG layer, DNA docking sites remain accessible and addressable enabling DNA binding assays even in degrading conditions.^32, 33^ While the highly improved stability and preserved functionality of coated DNA nanostructures broadens the scope of applications, the verification of the coating process and its quality is still time-consuming, highly invasive or rather indirect. So far, the silicification of DNA origami has been investigated either by transmission electron microscopy (TEM), atomic force microscopy (AFM) or x-ray spectroscopy techniques such as energy dispersive X-ray spectroscopy (EDX) or Small-angle X-ray scattering (SAXS).^24–26, 30, 31, 33^ While the sub-nanometer thick PLL-PEG coating is hardly visible in AFM (Figure 1b), it can be characterized by TEM or probed by gel electrophoresis, as DNA origami completely covered with cationic polymer lose their charge and electrophoretic mobility.^27–29, 32^ While gel electrophoresis is the most commonly used quality check for the coating with PLL-PEG, this technique remains blind to aggregates, that can occur during coating in solution, also resulting in a suppressed electrophoretic mobility.^27, 29^ Imaging techniques such as TEM, AFM or X-ray spectroscopy, on the other hand, are highly invasive or time-consuming preventing a quick and easy characterization of coated nanostructures under application conditions. Leveraging the non-invasive nature and sensitivity of single molecule fluorescence imaging, here, we report a novel strategy to study the coating of DNA nanodevices on a single structure level.

**Figure 1.**
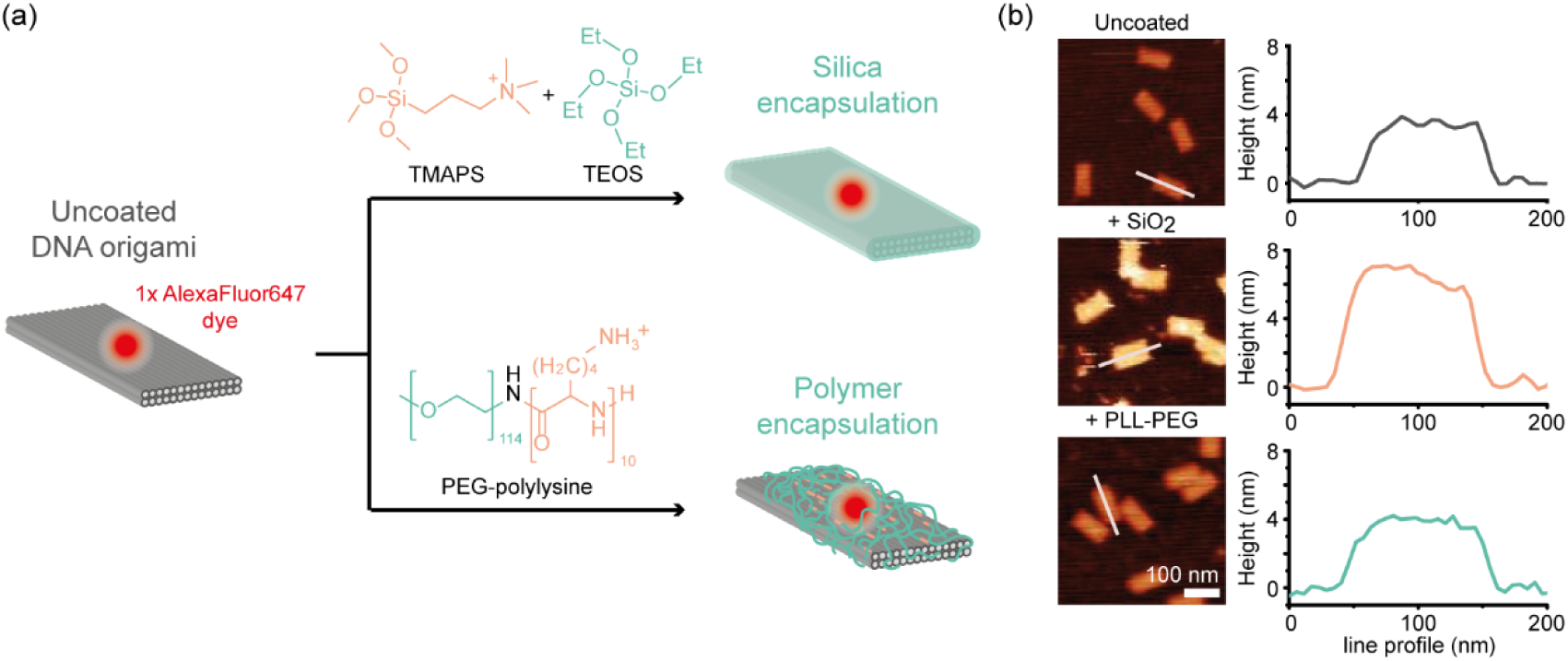
Principle of coating a DNA origami and its characterisation using AFM. (a) Scheme of Two-Layer Origami (TLO) with a single AlexaFluor647 label, coated by silica or by PLL-PEG. (b) Exemplary AFM scans and height-profiles of uncoated and coated TLOs exhibit a measurable height increase only for coating by silica. Scalebar is 100 nm.

Since a coating layer increases the local viscosity and at the same time decreases the steric freedom of chemical modifications on the DNA origami, we reasoned that labeling a DNA nanostructure with an environment-sensitive fluorophore at selected positions could enable probing of the coating process with nanometer precision on a single nanostructure level. Cyanine dyes are highly sensitive to their microenvironment since steric hindrance or a high viscosity slows down the photoisomerization from the emissive *trans* to the non-emissive *cis* state resulting in an increased fluorescence lifetime and photon count rate.^34, 35^ This effect has been exploited to study the binding of proteins or nucleic acids in the closed vicinity of a cyanine dye and has been termed protein-induced (or photoisomerization-related) fluorescence enhancement (PIFE).^36–38^ Additionally, a fluorophore embedded in the coating layer is also potentially less accessible for solvent molecules. For silicification, for example, the displacement of more than 40 % of the internal hydration water has been reported, indicating a strong hydrophobic condensation effect within silicified DNA nanostructures.^31^ Water has been shown to quench the fluorescence of red-emitting dyes (absorption and emission > 600 nm) via a resonant energy transfer from the excited S_1_ state of the fluorophore to harmonics and combination bands of OH vibrational modes in the H_2_O molecule.^39^ Quenching can be suppressed by replacing water with its heavy analogue D_2_O leading to an increased fluorescence lifetime and photon count rate of the dye.^40, 41^ This effect has been exploited to sense the number of water molecules in the hydration sphere of red-emitting dyes encapsulated in reverse micelles.^42^ Expecting that a coating layer on a DNA origami changes the local environment and decreases the concentration of water molecules in the hydration sphere of an embedded fluorophore, we reasoned that the red-emitting cyanine dye AlexaFluor647 (AF647) could be a suitable candidate as a molecular sensor for the coating process. To remain independent of laser and setup fluctuations, we chose the fluorescence lifetime as a non-invasive readout to investigate the coating of single, AF647 labeled DNA origami nanostructures with PLL-PEG and silica. We demonstrate the feasibility of the molecular sensor to probe the coating at different positions on the nanostructure and its stability in degrading conditions (low ionic strength, degrading enzymes), even without photostabilization or without time-consuming post processing. Finally, we combine the fluorescence lifetime sensor with DNA points accumulation for imaging in nanoscale topography (DNA-PAINT) super-resolution imaging to simultaneously probe the coating layer, the structural integrity of coated DNA origami and the retained addressability of DNA docking sites.

## RESULTS

To test, whether the designed molecular fluorescence lifetime sensor can probe the coating on DNA nanostructures, we designed a two-layer DNA origami nanostructure (TLO, Figure 1a). The TLOs were equipped with multiple biotin-labelled staple strands to enable immobilization on neutravidin or streptavidin functionalized glass slides and were labeled with environment-sensitive AF647. Before investigating the fluorescence lifetime of the cyanine dye, we first aimed to verify the successful coating of immobilized TLO nanostructures by PLL-PEG or silica via AFM as previously reported for silicified DNA origami (see Figure S2a).^25, 26, 30, 33^ Uncoated TLO exhibited a height of around 4 nm, while silicification led to a height increase of around 2 nm. In the case of the PLL-PEG coating, no significant height increase was visible in the AFM scans, either because the encapsulation shell is too thin (sub-nanometer thickness as measured in cryo-EM studies) or not rigid enough to be measured by the AFM cantilever. Nevertheless, the successful coating by PLL-PEG could be confirmed by subsequent incubation in degrading low salt conditions (Mg^2+^ free and EDTA-containing buffer). While uncoated TLO degraded and collapsed into rod-shaped debris, the PLL-PEG and silica-coated nanostructures both remained intact, indicating successful protection for PLL-PEG coating otherwise undetectable in AFM imaging (see Figure S2b).

Next, we investigated the fluorescence lifetime of the cyanine label at different positions on uncoated and coated DNA nanostructures. To this end, we immobilized the nanostructures on biotinylated BSA and NeutrAvidin functionalized microscope glass slides and coated them with PLL-PEG or silica. Single-molecule fluorescence lifetime imaging microscopy (FLIM) scans were acquired on a custom-built confocal microscope with a time-correlated single photon counting unit (TCSPC).^43^ Acquired FLIM scans were analyzed by picking individual nanostructures and extracting the spot-integrated fluorescence lifetime information. Distributions of the obtained fluorescence lifetime values were then fitted by Gaussian distribution functions, to determine the mean of each individual distribution. First, we labelled the TLO nanostructure internally with a single AF647, *i.e.,* directly at the 3’-end of a selected staple strand, to ensure that the sensor dye is embedded in the coating layer. To probe the coating process at the surface and inside of the DNA nanostructure, we once placed the sensor dye internally at the upper DNA surface and once internally at the interface of the two DNA layers (Figure 2). FLIM scans of uncoated DNA origami (Figure 2a) revealed that the two labeling positions result in different microenvironments of the AF647 label and consequently in different fluorescence lifetime distributions. The fluorescence lifetime distribution of the internal sensor label at the upper surface of the TLO origami revealed a sharp peak around 1.08 ± 0.05 ns close to the reported fluorescence lifetime of a free AF647 dye (Figure 2a, first row). The fluorescence lifetime distribution of the internal label at the DNA interface within the TLO design though revealed two populations of higher lifetimes (for fitted values see Table S5) indicating a more complex environment of the dye within the DNA origami with more steric restriction and possible interactions of the cyanine dye with the DNA origami backbone. As proposed in our sensor design, coating with PLL-PEG (Figure 2b) or silica (Figure 2c) resulted in similar, significant increase in the fluorescence lifetime of AF647 for both labeling positions of at least 0.2 ns (Table S5). For the AF647 label at the surface of the TLO, the initially single Gaussian distribution was shifted by both coating agents to two populations of higher fluorescence lifetimes indicating two different microenvironments of the dye after the formation of the protective layers. For the TLO labeled with AF647 internally at the DNA interface, both coating agents induced a shift of the initial two fluorescence lifetime populations to higher fluorescence lifetimes. The similar fluorescence lifetime shifts for both coating agents indicate that our molecular sensor design works independently of the coating material and can be employed to probe coating-induced effects, such as steric restriction or water repulsion, not only at the surface of a coated DNA nanostructure but also at a position inside of it.

**Figure 2.**
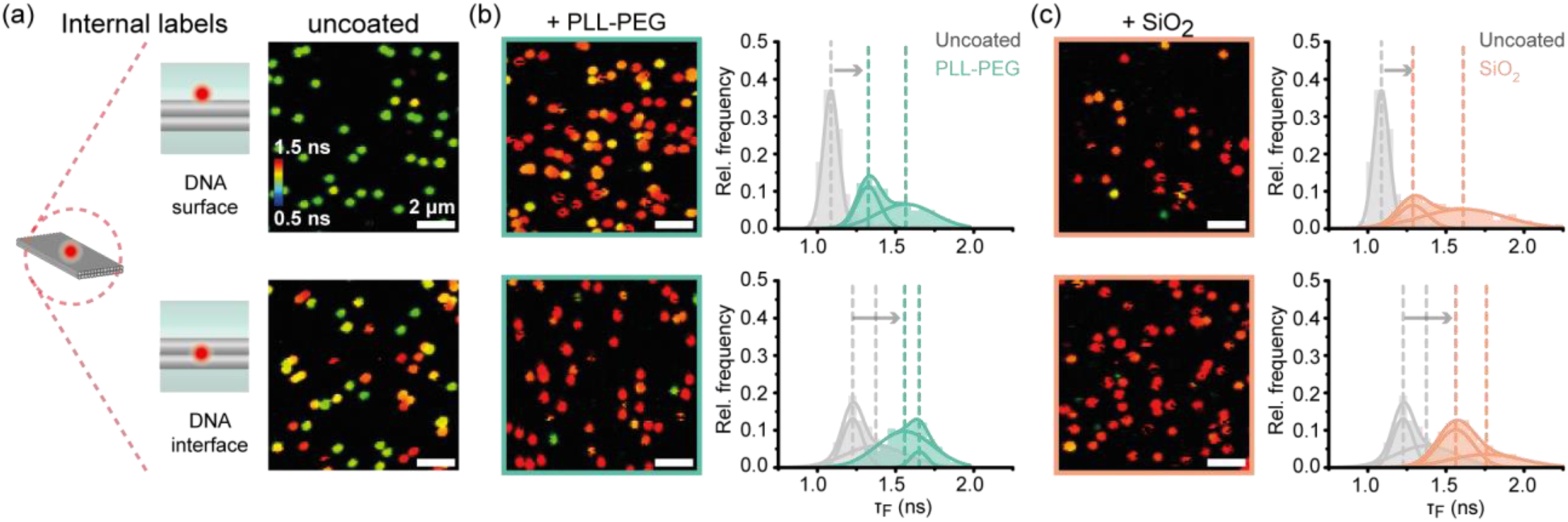
Characterization of the fluorescence lifetime-based sensor to probe the coating of different positions of DNA origami nanostructures by directly modifying selected staple strands (internal label). a) Different internal labeling positions of the AF647 fluorophore on the TLO (left). FLIM scans of Alexa647-labelled TLO origami after immobilization (right). b) FLIM scans after coating with PLL-PEG and spot-integrated lifetime distributions. c) FLIM scans after silicification and spot-integrated lifetime distributions. Dotted lines indicate mean values of Gaussian fits.

To exploit the *in situ* sensing ability and to better understand the coating process we studied the different DNA origami coating strategies over time (Figures S3 and S4). Already after 30 minutes, we observed a complete fluorescence lifetime shift when the TLO nanostructures were incubated with PLL-PEG indicating the rapid formation of the protective polymer layer. A complete fluorescence lifetime shift for silicification, on the other hand, required an incubation of the precursor solution for at least 24 h, which is in agreement with previously reported silicification kinetics.^31^ To obtain absolute lifetime values, the acquired lifetime decay curve of every picked fluorescent spot in the FLIM scans representing a single DNA origami nanostructure was re-convoluted with the measured instrument response function (IRF) of the confocal microscope (Figure S5). As the absolute lifetime shift induced by the coating layers was unaffected by the re-convolution step, spot-integrated lifetime populations were used throughout this study to highlight the fast and straight-forward readout of the coating process (for more details see SI Section 1.10). While FLIM imaging was performed in a photostabilization buffer (see Materials and Methods section for details) to obtain high photon numbers, the fluorescence lifetime shift after coating with PLL-PEG or silica could also be obtained from FLIM scans performed without photostabilization (Figure S6). This allowed for probing the coating process of DNA origami *in situ* and in different application conditions without the need of specialized imaging buffer.

We further aimed to probe the coating process at different labeling positions using a more modular and less costly labeling strategy. To this end, we used a 3’-AF647-labelled, 21-nt oligonucleotide which can hybridize externally to a complementary ss-DNA extension positioned on DNA origami. In this manner, a single fluorescently labeled oligonucleotide is sufficient to screen the presence of the coating at different positions on various nanostructures making the approach more cost-effective (Figure 3). We then probed the homogeneity of the coating layers at different positions on the TLO by positioning the DNA docking sites for external labelling either in the center or in a corner of the upper DNA origami surface. To underline that our molecular sensor can be applied to any DNA origami nanostructure, we also studied AF647 labeled twelve helix bundle (12HB) DNA origami nanostructures, which is based on honeycomb lattice compared to the square lattice of TLO.

**Figure 3.**
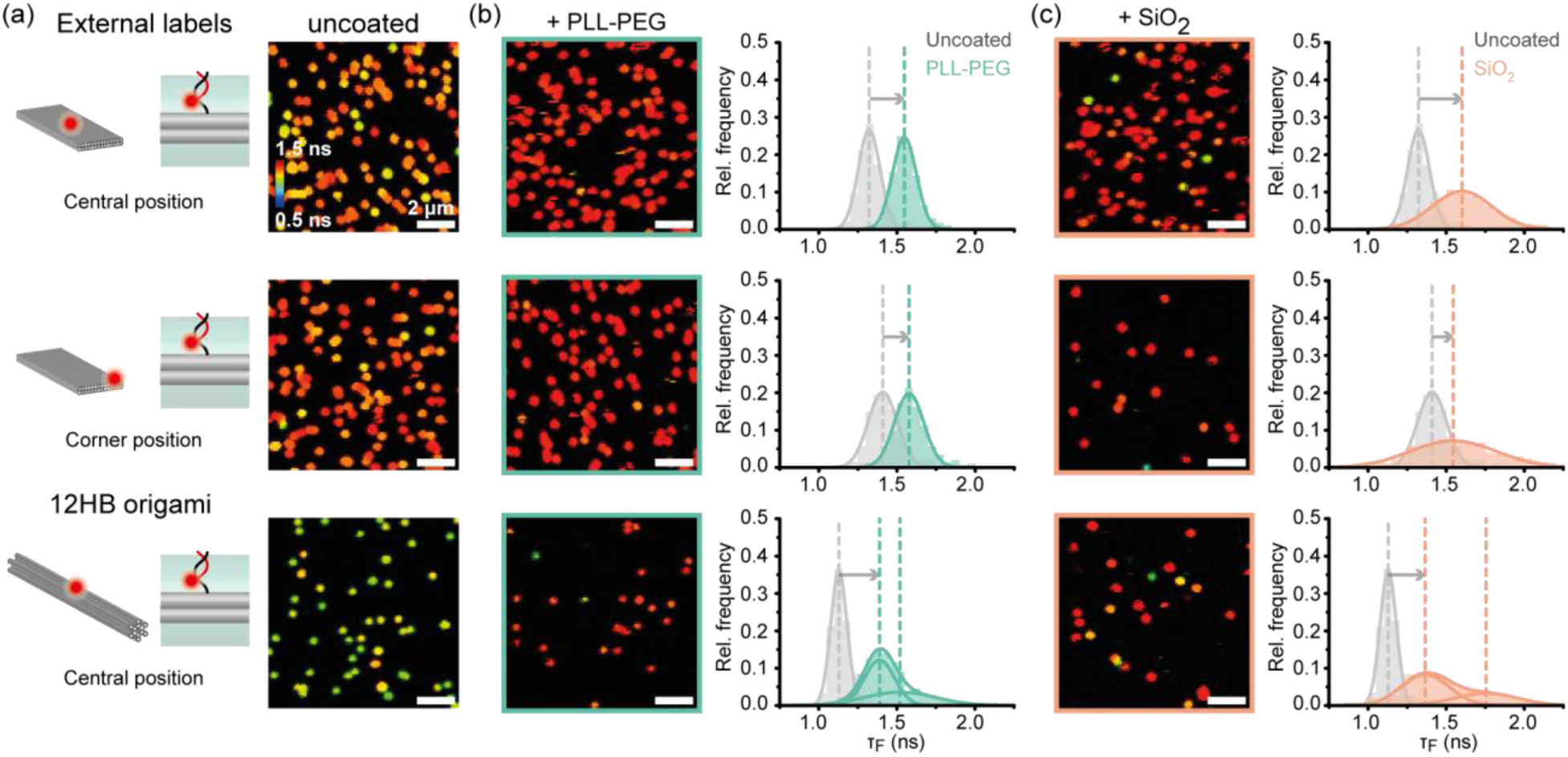
Characterization of fluorescence lifetime-based sensor to probe the coating of different positions of DNA origami nanostructures by externally binding a fluorescently modified imager strand via hybridization (external label). a) Different labeling positions of a single AF647 fluorophore on the TLO (left). FLIM scans Alexa647-labelled TLO origami after immobilization (right). b) FLIM scans after coating with PLL-PEG and spot-integrated lifetime distributions. c) FLIM scans after silicification and spot-integrated lifetime distributions. Dotted lines indicate mean values of Gaussian distribution fits.

Using the external labeling approach at different positions on the DNA origami led to different microenvironments of the cyanine dye and, in turn, different fluorescence lifetime distributions for uncoated DNA origami (Figure 3a). While AF647 labels bound externally to uncoated TLO exhibited higher lifetimes (Table S5) than the internal dye at the DNA surface, indicating an already higher restricted microenvironment on the external DNA docking sites, the external cyanine label on the 12HB origami exhibited a lower fluorescence lifetime distribution indicating a relatively free microenvironment. Again, coating with both PLL-PEG and silica induced similar shifts in the fluorescence lifetime distributions independently of the labeling position or DNA origami design (Table S5). While revealing different fluorescence lifetimes for uncoated TLO origami, both external cyanine labels showed similar fluorescence lifetime populations after coating with both silica or PLL-PEG. The external label on the 12HB DNA origami, on the other hand, revealed two fluorescence lifetime subpopulations after coating with either PLL-PEG or silica indicating a more restricted but more heterogenous microenvironment of the dye than before the coating. Except for the case of coated 12HB nanostructures, the external labels generally revealed unimodal distributions indicating simpler binding situations than for the internal labels, as shown in Figure 3.

After successfully employing internal and external dye labeling strategies to probe the coating of DNA nanostructures at different positions, we went on to test the reversibility of the FLIM sensor by probing the fluorescence lifetime after the coating with PLL-PEG and after subsequent removal of the polymer coating by the addition of an anionic polymer. To this end, PLL-PEG coated TLO nanostructures were incubated with anionic dextran sulfate, which has been reported to decomplex and thus remove the cationic polymer coating.^28^ Indeed, the fluorescence lifetime distribution of PLL-PEG coated TLO with an external cyanine label at the center of the upper surface revealed a quantitative shift back to the initial lifetime distribution of the uncoated nanostructure (Figure 4). Accordingly, we also observed a reversible shift of the fluorescence lifetime for the molecular sensors at internal label positions on the upper surface and at the interface inside the TLO design (Figure S7) highlighting the reversibility of the molecular sensor and the polymer coating independently on the labeling position. Furthermore, we studied the decomplexation process in real-time by scanning the same field of view over time after addition of dextran sulfate and saw a complete shift back to uncoated nanostructures within the first 5 minutes highlighting the fast kinetics of this process (Figure S8).

**Figure 4.**
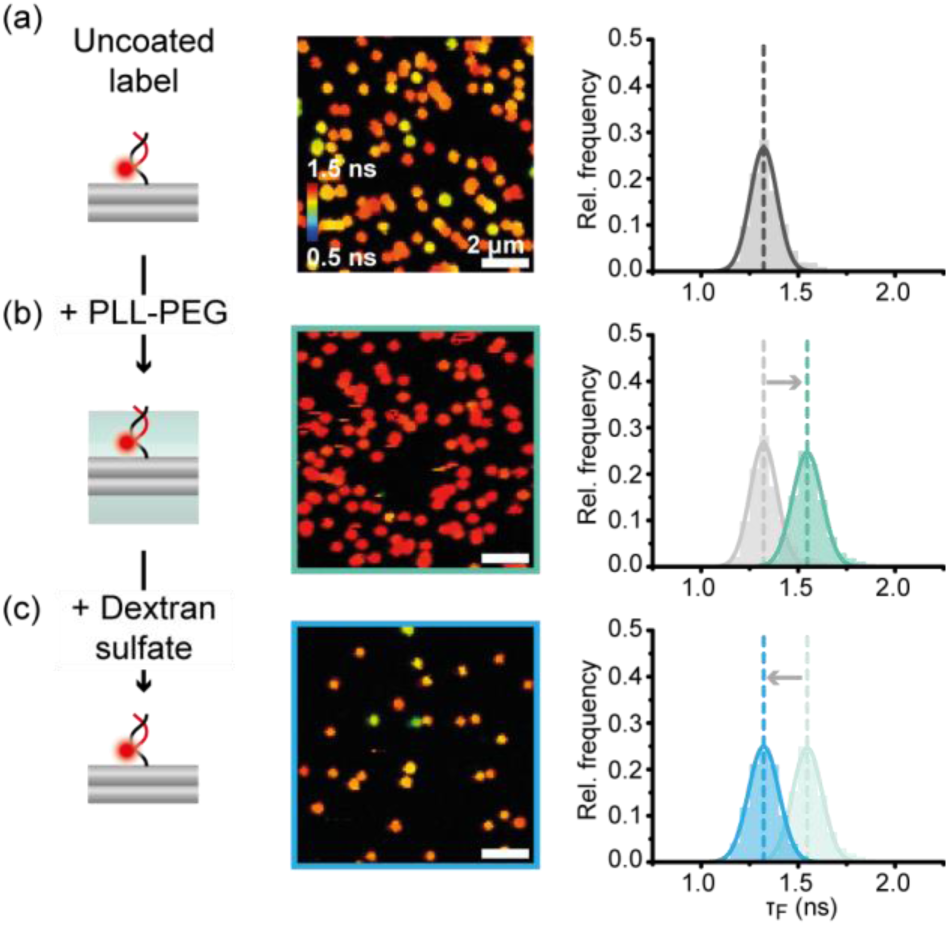
Probing the reversible coating with PLL-PEG and dextran sulfate on a TLO nanostructure with an external label. a) FLIM scan and fluorescence lifetime distribution for uncoated TLO with the externally labeled sensor dye. b) FLIM scan and spot-integrated fluorescence lifetime distribution for PLL-PEG coated TLO with the externally labeled sensor dye reveals a lifetime shift to higher values. c) FLIM scan and spot-integrated fluorescence lifetime distribution for initially PLL-PEG coated TLO from b) reveals a quantitative shift back to a lifetime distribution corresponding to uncoated TLO nanostructures.

The quantitative reversibility of the molecular sensor enables not only the investigation of the integrity of the DNA nanostructure but also of the coating layers at a single nanostructure level and in real-time. We thus next aimed to monitor the stability of the DNA origami and the coating layers in harsh and degrading conditions by first confirming that the observed fluorescence lifetime shifts correlate with the expected improved stability. To this end, we coated TLO labelled with AF647 at different positions with PLL-PEG or silica and subsequent exposure the nanostructures to degrading conditions, *i.e.,* Mg^2+^ free buffer containing EDTA (Figure 5a, c) or a solution containing degrading enzyme (DNAse I, Figure 5b, d). First, the qualitative degradation of uncoated TLO was probed in real-time by applying the degrading solutions and subsequently acquiring FLIM images of the same field of view every 5 min (Figure S9). Incubation of nanostructures in Mg^2+^ free buffer led to complete degradation of uncoated TLO within the first 5 min, while addition of DNAse I degraded all nanostructures within the first 15 min, highlighting the low stability of uncoated DNA origami nanostructures in low-salt conditions and in the presence of nucleases (Figures 5a, 5b, and S9a). PLL-PEG and silica-coated structures, on the other hand, survived the incubation by either low salt buffers or DNAse I over the full 30 min tested, indicating the successful stabilization by the protective coatings (Figure S9). Since the coated nanostructures revealed higher fluorescence lifetimes even after 30 min incubation in degrading conditions, we concluded that a higher fluorescence lifetime of the sensor dye indeed goes hand in hand with successful coating and the effective stabilization of DNA origami nanostructures in degrading conditions.

**Figure 5.**
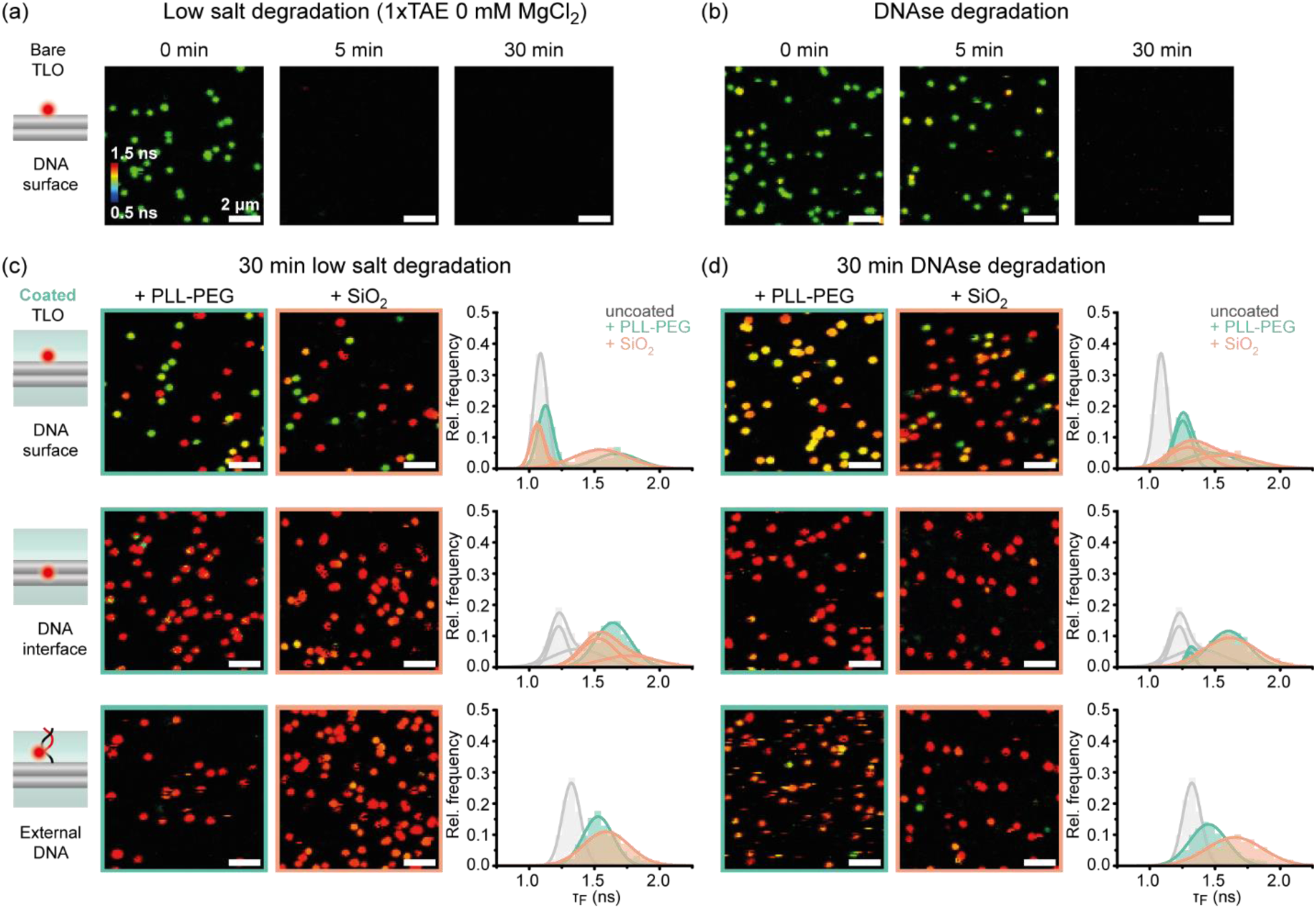
Using the fluorescence lifetime sensor to probe the stability of DNA origami structures and the coating layers in degrading conditions. a) Real-time degradation study of uncoated TLO in Mg^2+^ free 1×TAE buffer. b) Real-time degradation study of uncoated TLO in DNAse I solution. c) FLIM scans and spot-integrated fluorescence lifetime distributions for PLL-PEG and silica coated TLOs after 30 min incubation in Mg^2+^ 1×TAE buffer. d) FLIM scans and spot-integrated fluorescence lifetime distributions for PLL-PEG (green) and silica (orange) coated TLOs after 30 min incubation in DNAse I solution. Grey graphs represent fluorescence lifetime distributions of intact, uncoated TLO as reference.

To investigate the integrity of the coating layers further, we quantified the shift in fluorescence lifetime distributions of PLL-PEG and silica coated TLOs after 30 min incubation in either Mg^2+^ free buffer containing EDTA or in the presence of DNAse I (Figure 5c and 5d). In general, all coated TLO nanostructures withstood the degrading conditions and showed preserved high fluorescence lifetime populations corresponding to an intact coating layer. Only the TLO with an internal AF647 label at the DNA surface exhibited a partial degradation of both the polymer and silica coating when incubated in a Mg^2+^-free buffer (as indicated by the arising peak around 1.05 ns in fluorescence lifetime distribution previously assigned to uncoated TLO (Figure 3c, first row, Table S10). To test if the nanostructures contributing to this fluorescence lifetime population are still stabilized by a partially degraded coating or whether they are indeed uncoated and, thus, prone to degradation, we incubated the resulting sample additionally in DNAse I solution. The second degradation step resulted in loss of nanostructures giving rise to fluorescence lifetimes around 1.05 ns, indicating that the observed lower fluorescence lifetime populations can be attributed to DNA origami structures that had compromised coating upon the first degradation (Figure S10). These results highlight the sensitivity of the FLIM sensor and showcase its ability to probe the stability of different DNA origami coating agents in degrading conditions ultimately aiding the optimization of existing and development of new protective strategies.

Last, we combined our molecular sensor with DNA PAINT super-resolution imaging to highlight its applicability for correlative single-molecule imaging techniques.^44^ To this end, we incorporated DNA PAINT docking sites in a rectangular pattern (lengths of ca. 30 and 65 nm) into a TLO nanostructure internally labeled with an AF647 dye at the surface (Figure 6a). This design allowed us to obtain complementary information about the coated nanostructures: while the FLIM sensor reports on the successful coating of the nanostructure, DNA PAINT imaging provides insights into the structural integrity of the DNA origami design and into the addressability of designed DNA docking sites. In this matter, we first acquired FLIM scans of uncoated and coated TLO nanostructures and subsequently performed DNA PAINT imaging of the same samples on a wide-field microscope. A shift of the fluorescence lifetime distributions of PLL-PEG and silica-coated nanostructures indicated again the successful coating for both coating strategies (Figure 6b, c; fitted values in Table S11). After acquiring FLIM scans, the imaging buffer was exchanged with a Cy3B labeled DNA PAINT imager solution and the same slides of uncoated or coated DNA origami were imaged on a total internal reflection fluorescence microscopy (TIRFM) widefield setup with 532 nm excitation to obtain super-resolved DNA PAINT images (Figure 6d). Distance analysis of picked nanostructures revealed the same rectangular labeling pattern with comparable distances for both coating agents as for uncoated TLO, highlighting the structural integrity after the coating process (Figure S11).

**Figure 6.**
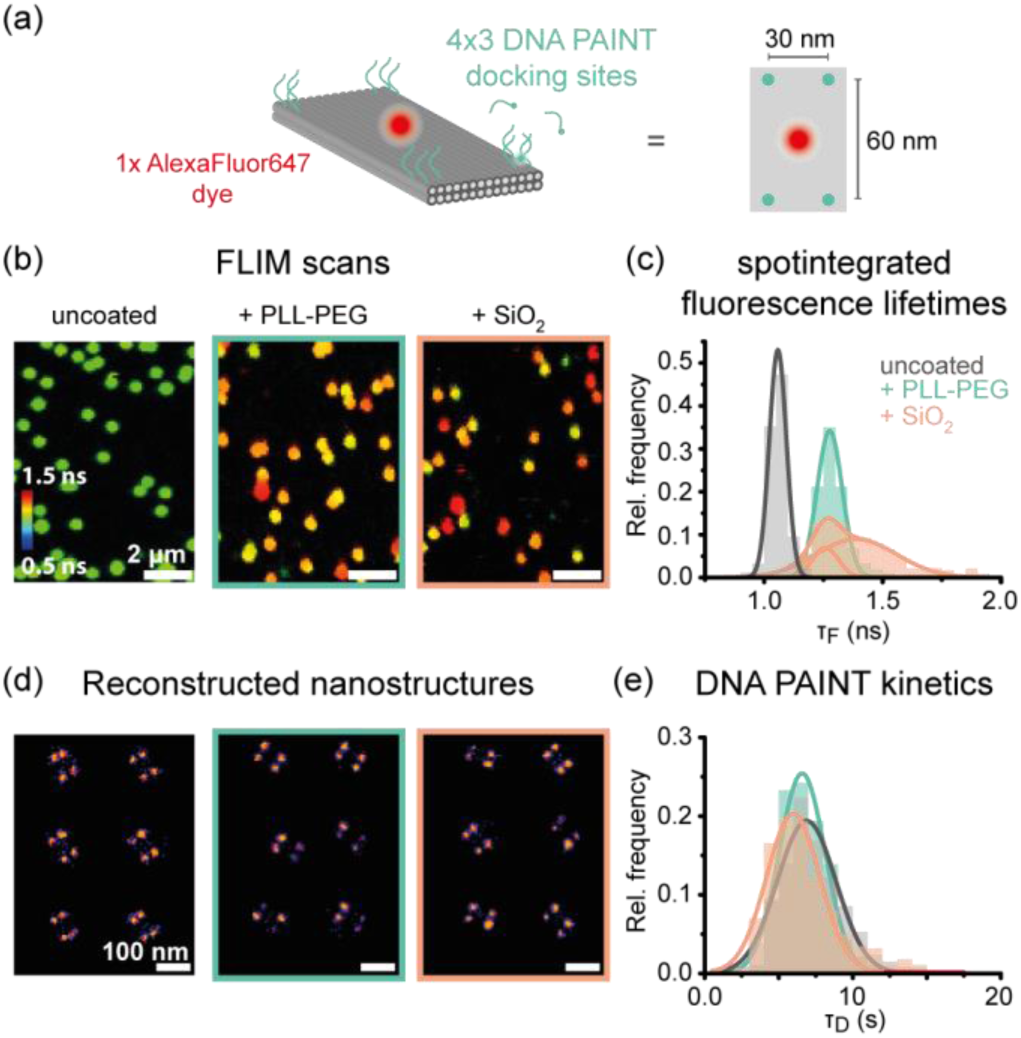
Combining the molecular sensor with DNA PAINT imaging for simultaneous investigation of the coating, structural integrity of the nanostructure, and addressability of DNA docking sites. a) Labeling scheme of a TLO internally labeled with AF647 at the upper surface and 4× three DNA PAINT docking sites in the corner regions of the TLO design. b) FLIM scans of uncoated, PLL-PEG and silica coated TLO indicate successful coating. c) Spot-integrated fluorescence lifetime distributions of uncoated, PLL-PEG and silica coated TLO reveal a shift upon coating. d) Picked and aligned DNA PAINT images of uncoated, PLL-PEG and silica coated TLO revealing the designed rectangular geometry. e) Extracted DNA PAINT dark time (τ_D_) distributions for uncoated, PLL-PEG and silica coated TLO indicate similar accessibility for imager strands in solution.

Still, for both coating strategies a small subpopulation of a dual spot pattern was observed, which could be interpreted as folding of the TLO origami along its longer axis induced by the coating agent. From time to time, we observed this effect also in AFM imaging where a subpopulation of TLO nanostructures revealed a rod-shaped geometry with an increased height (Figure S12). Similar folding defects have been previously observed for the coating of monolayer DNA origami with PLL-PEG but with higher efficiencies (>50%) due to the higher flexibility of the investigated nanostructure.^28^ By extracting the DNA PAINT dark times, *i.e.,* the time between two binding events, the accessibility of the DNA PAINT docking sites on uncoated and coated nanostructures was compared. Both PLL-PEG and silica-coated TLO nanostructures revealed similar dark times as the uncoated TLO (Figure 6E, fitted values in Table S12), highlighting the preserved accessibility and addressability of DNA docking sites even after the formation of a protective layer.^32, 33^

## DISCUSSION

Our results demonstrate that the red cyanine dye AF647 is a suitable molecular FLIM sensor for probing the coating of DNA origami nanostructures in real time. However, the high sensitivity of the molecular sensor to its microenvironment makes it also susceptible to variations in fluorescence lifetime when labelled at different positions on the DNA origami. Consequently, for every labeling position, the sensor dye revealed a distinct fluorescence lifetime distribution before and after the coating which in turn can lead to varying fluorescence lifetime shifts. Nevertheless, we observed a pronounced absolute increase in the fluorescence lifetime in the range of 0.2 to 0.5 ns for all labeling positions, independent of the coating material.

To better understand the mechanistic origin of the observed fluorescence lifetime shift upon coating the DNA nanostructure, we carried out further mechanistic studies with the TLO origami labelled with an internal AF647 at the DNA surface. To quantify the sensitivity of the AF647 label on the DNA origami towards quenching by water, we measured the fluorescence lifetime in a D_2_O buffer (Figure S13) and observed a shift of around 0.4 ns, similar to the observed fluorescence lifetime shifts upon coating with PLL-PEG or silica. To further investigate the role of a restricted photoisomerization of the cyanine dye in the coating layer, we performed fluorescence intensity correlation analysis of single-molecule trajectories of AF647 immobilized on DNA origami before and after coating (Figure S14). While quenching by water only affects the S_1_ excited state of the fluorophore, a shift in the photoisomerization rate affects the photophysics of the dye (*i.e.,* occurring dark states), which can be probed via intensity autocorrelation. To shed more light on two different fluorescence lifetime populations that were observed for coated DNA origami nanostructures, we grouped the intensity autocorrelation curves obtained from single-molecule trajectories to two populations based on their fluorescence lifetimes as observed in Figure 2. Intensity autocorrelation curves obtained from PLL-PEG coated TLOs with shorter fluorescence lifetime were comparable to those of uncoated TLOs indicating that the observed fluorescence lifetime shift is not related to the restricted photoisomerization or other effects on dye photo-physics, but perhaps is predominantly induced by reduced water quenching. In the intensity autocorrelation curves of PLL-PEG coated TLOs with higher fluorescence lifetime, we observed a slowed down photoisomerization with a reduced amplitude, indicating that the larger fluorescence lifetime shift could be additionally induced by restricted photoisomerization (for more detailed discussion, see Supplementary Note 1). Even though we found similar trends for silica coating, no significant difference between the lower and the higher fluorescence lifetime subpopulations was observed which we attributed to a dynamic interchange of the dye between two environments with different fluorescence lifetimes. This was also visible in continuous FLIM scans of the same field of view and far more pronounced for the silica-coated nanostructures (Figure S15). Altogether, the mechanistic studies suggest, that both effects, a restricted photoisomerization and reduced water quenching, contribute to the fluorescence lifetime shifts upon coating whereas the exact microenvironment around the dye within the coating layer determine which effect has a higher impact.

Correlative FLIM and DNA PAINT imaging revealed a deformed subpopulation of the TLO nanostructures induced by PLL-PEG and silica coating, as it has been reported for PLL-PEG-coated DNA monolayer origamis previously. This highlights that coating-induced effects can lead to severe deformation even for rigid nanostructures and showcases that non-invasive fluorescence-based methods as the one reported here could be suitable tools to assess these deformations or artifacts induced by the coating agents, especially on a single nanostructure level.

While we performed FLIM measurements on an advanced single-molecule microscope to study the coating layer on a single nanostructure level, one could also envision that ensemble lifetime readout could be used for a quick assessment of the coating of the nanostructures in solution. To further improve the fluorescence contrast of the molecular sensor, in the future one could explore a palette of alternative environment-sensitive dyes, *e.g.* red-emitting dyes such as Atto647N or Cy7 which have been reported to be quenched by water even more efficiently.^39^ Self-blinking dyes like silica rhodamines, on the other hand, could potentially be applied to our sensor design to realize a fluorescence blinking or intensity-based readout, enabling the probing of the coating layer also on widefield microscopes.^45^

## CONCLUSION

In this work we exploited the environment-sensitive cyanine dye AF647 to design a simple single-molecule sensor to probe the protective coating of DNA origami nanostructures by either the block copolymer PLL-PEG or by silica. By acquiring FLIM scans, the lifetime shift towards longer lifetimes can be utilized to screen the coating process qualitatively on a single nanostructure level, non-invasively and in real-time. By placing the fluorophore at different positions on the DNA origami, the coating process and its effects on the nanostructure were investigated at the position of interest with nanometer precision. Further mechanistic studies suggested that both reduced quenching by water and restricted photoisomerization in the coating layer could be attributed to the observed fluorescence lifetime increase of the employed sensor dye AF647. The reversibility of the molecular FLIM sensor could be exploited to follow the quantitative decomplexation of PLL-PEG coating by dextran sulfate and to investigate the integrity of the coating in degrading conditions in real time. By combining the molecular sensor design with DNA-PAINT we could probe for the first time the successful coating, structural integrity and addressability of DNA docking sites on the same sample, which was previously not possible due to the invasive nature of structural characterization methods, such as TEM or AFM. The observed independence of the fluorescence lifetime shift on the coating material makes it a potential tool to study other coating strategies, such as the encapsulation of DNA nanostructures with proteins.^46^ The non-invasive character and the possibility to combine the FLIM sensor design with other single-molecule techniques such as super-resolution microscopy or fluorescence resonance energy transfer (FRET) enables the probing of coatings in a multitude of applications, such as drug delivery and release, biosensing, or biocomputing.

## METHODS

### Materials

The p8064 scaffold strand used for the folding of the DNA origami nanostructures was extracted from M13mp18 bacteriophages (produced in-house). Unmodified staple strands were purchased from Eurofins Genomics GmbH (Germany) and Integrated DNA Technologies (USA). Dye labeled oligonucleotides for DNA PAINT imaging or permanent labeling were purchased from Eurofins Genomics GmbH (Germany).

### DNA origami folding and purification

All investigated TLO DNA origami nanostructures (also shown in Figure S1) were folded in a 1× TE buffer containing 12 mM MgCl_2_ with a linear thermal annealing ramp from 60°C to 44°C with 1h/°C after an initial 65°C denaturation step. The 12HB DNA origami nanostructures were folded in a 1× TAE buffer containing 16 mM MgCl_2_ using the same scaffold strand as for the TLO. The structures were folded with a non-linear thermal annealing ramp starting at 65 °C and then cooling down to 4 °C over a period of 25 hours.^47^ Modifications of the DNA origami were realized using caDNAno (version 2.2.0). A full list of unmodified and modified staple strands and sequences of are given in Table S2-4, S13 and S14 in the Supporting Information. Folded DNA origami nanostructures were purified by filtration using Amicon Ultra filters (100 K, Merck, Germany). Concentrations of purified sample solution were measured via UV/vis spectroscopy (NanoDrop, Fischer Scientific, USA). More details on DNA origami design, folding and purification procedures are given in SI, Section 1.1.

### Sample preparation

Cleaned high precision µm microscope cover glass (170 µm, 22 × 22 mm, No. 1.5H glass slides, Carl Roth GmbH, Germany) were assembled into inverted flow chambers as described previously.^48^ The assembled chambers were passivated with BSA-biotin (Sigma Aldrich, USA) and functionalized with either neutravidin or streptavidin (Sigma Aldrich, USA). For immobilization, purified DNA origami was diluted to approximately 50 pM in 1× PBS buffer containing 500 mM NaCl and incubated in the chambers for ca. 5 minutes and stored in a 1× TAE containing 10 mM MgCl_2_. Sufficient surface density was probed with a TIRF microscope. For more details on sample preparation, see SI Section 1.5.

### Coating with PLL-PEG or silica

The PLL-PEG block copolymer K10PEG (1K) (Alamanda polymers, USA) was dissolved and stored in ultra-pure water at a concentration of 2 mM.^27, 28^ Aliquots were stored at −20 °C and thawed and ultrasonicated for 10 min before use. To coat immobilized DNA origami nanostructures, the 2 mM PLL-PEG solution was diluted in a 1× TAE buffer containing 10 mM MgCl_2_ to a final concentration of 20 µM and incubated for 30 min. To decomplex the cationic PLL-PEG coating from the DNA origami, a 20 µM solution of anionic dextran sulfate (Sigma Aldrich, M = 20 000 g/mol)^28^ in 1× TAE 10 mM MgCl_2_ was incubated in the coated sample chambers for 30 min. After washing, samples were then stored in 1× TAE containing 10 mM MgCl_2_. For the silicification of immobilized DNA origami an adapted version of the protocol of Fan and co-workers was applied.^26, 33^ After hydrolysis of 100 µL TMAPS (50% (wt/wt) in methanol, TCI America) in 5 mL 1× TAE (40 mM Tris, 2 mM EDTA, 12.5 mM MgAc_2_, pH=8.0) for 20 min under vigorous stirring, 100 µL TEOS (98%, Sigma Aldrich, USA) was added and stirred for another 20 min. Freshly prepared precursor solution was incubated for 24 h. The coated sample chambers were washed with 80% ethanol and with ultra-pure water. The samples were then stored in 1× TAE containing 10 mM MgCl_2_. For AFM imaging, mica slides with immobilized DNA origami were analogously incubated with either 20 µM PLL-PEG solution or with freshly prepared silica precursor solution. For more details on coating of immobilized DNA nanostructures, see SI Section 1.6.

### Degradation studies

To probe the stability of coated and bare DNA origami in degrading conditions, either a low-salt buffer or a DNAse I solution were incubated on immobilized nanostructures. Magnesium ion free conditions were realized by incubation of a 1× TAE buffer on surface immobilized nanostructures for 30 min. Enzymatic degradation of immobilized DNA origami nanostructures was tested by incubation of a DNAse I solution (1:10 dilution of DNase I (1 U/μl) in 1× TAE containing 10 mM MgCl_2_, Thermo Fisher Scientific, USA) for 30 min.

### AFM imaging

AFM scans in aqueous solution (AFM buffer = 40 mM Tris, 2 mM EDTA, 12.5 mM Mg(OAc)_2_·4 H_2_O) were performed on a NanoWizard® 3 ultra AFM (JPK Instruments AG). Measurements were performed in AC mode on a scan area of 3 x 3 µm with a micro cantilever (νres = 110 kHZ, kspring = 9 N/m, Olympus Corp.). Leveling, background correction and extraction of height histograms of obtained AFM images were realized with the software Gwyddion (version 2.60).^49^ For more details on sample preparation and AFM imaging, see SI Section 1.4.

### Confocal microscopy

Fluorescence lifetime imaging microscopy (FLIM) and intensity autocorrelation studies were performed on a home-built confocal microscope based on an Olympus IX-71 inverted microscope as described previously.^50^ AF647 modifications labeled to surface-immobilized DNA origami were excited by a pulsed 640 nm excitation at a repetition rate of 40 MHz. The setup was controlled by a commercial software package (SymPhoTime64, PicoQuant GmbH, Germany). For more details on FLIM imaging and intensity autocorrelation studies, see SI Section 1.8.

### Wide-field microscopy

DNA-PAINT measurements were carried out on a commercial Nanoimager S (ONI Ltd., UK). Red excitation at 640 nm was realized with a 1100 mW laser, green excitation at 532 nm with a 1000 mW laser, respectively. For imaging, a 1× PBS buffer containing 500 mM NaCl and an imager concentration of 5 nM was used. The 8 nt imager oligonucleotide with a Cy3B label on the 3’-end was purchased from Eurofins Genomics GmbH (Germany) and consisted of the sequence 5’-GGAATGTT-3’. Acquired DNA-PAINT raw data were analyzed using the Picasso software package.^48^ After drift correction, individual DNA nanostructures were picked, aligned and corresponding blinking kinetics extracted for further analysis. Distance analysis of obtained DNA PAINT images was performed with a custom written Python code. For more details on DNA PAINT imaging and data analysis, see SI Section 1.11.

## ASSOCIATED CONTENT

### Supporting Information

Supporting information includes additional information on the methods (Sections S1.1-S1.11), supplementary note on mechanistic studies on the fluorescence lifetime shift (Supplementary Note S1), Tables S1-14, and Figures S1-S18.

The following files are available free of charge. Supporting Information (PDF)

## AUTHOR INFORMATION

### Present Addresses

^†^If an author’s address is different than the one given in the affiliation line, this information may be included here.

### Author Contributions

V.G., M.S., and A.H.-J. conceived the idea. G.A.B. designed the TLO origami. M.S.‡ and G.A.B.‡ fabricated all samples and carried out AFM and FLIM imaging experiments, single-molecule intensity autocorrelation studies, and data analysis. T.S. contributed to the mechanistic intensity autocorrelation studies. K.B. and L.M.W. contributed to the silica coating experiments. M.S. and G.A.B. prepared figures. V.G., A.H.-J. and P.T. supervised the study. M.S., V.G. and G.A.B. wrote the manuscript with contributions from all authors. All authors have given approval to the final version of the manuscript.

## ACKNOWLEDGMENT

The authors thank M. Pinner and H. Dietz for providing an initial PLL-PEG sample for first tests with PLL-PEG coating. We also thank A. Szalai and G. Ferrari for providing the FLIM analysis Python code and E. Plötz and J. Bauer for fruitful discussions.

The authors have declared no competing interest.

## ABBREVIATIONS

AF647: AlexaFluor647
FLIM: Fluorescence lifetime imaging microscopy
TLO: two-layer DNA origami nanostructure
PLL-PEG: poly-*L*-lysine polyethylene glycol block copolymer
TEM: transmission electron microscopy
SAXS: Small-angle X-ray scattering
EDX: energy dispersive X-ray spectroscopy
FRET: fluorescence resonance energy transfer
AFM: atomic force microscopy
DNA-PAINT: DNA points accumulation for imaging in nanoscale topography
TCSPC: time-correlated single photon counting
TIRF: total internal reflection fluorescence microscopy.

## REFERENCES

(1) Rothemund, P. W. K. Folding DNA to create nanoscale shapes and patterns. Nature 2006, 440 (7082), 297–302. DOI: 10.1038/nature04586.

(2) Dey, S.; Fan, C.; Gothelf, K. V.; Li, J.; Lin, C.; Liu, L.; Liu, N.; Nijenhuis, M. A. D.; Saccà, B.; Simmel, F. C.;, et al. DNA origami. Nature Reviews Methods Primers 2021, 1 (1), 13. DOI: 10.1038/s43586-020-00009-8.

(3) Wang, S.; Zhou, Z.; Ma, N.; Yang, S.; Li, K.; Teng, C.; Ke, Y.; Tian, Y. DNA Origami-Enabled Biosensors. In Sensors, 2020; Vol. 20.

(4) Dass, M.; Gür, F. N.; Kołątaj, K.; Urban, M. J.; Liedl, T. DNA Origami-Enabled Plasmonic Sensing. The Journal of Physical Chemistry C 2021, 125 (11), 5969–5981. DOI: 10.1021/acs.jpcc.0c11238.

(5) Zhang, Q.; Jiang, Q.; Li, N.; Dai, L.; Liu, Q.; Song, L.; Wang, J.; Li, Y.; Tian, J.; Ding, B.;, et al. DNA Origami as an In Vivo Drug Delivery Vehicle for Cancer Therapy. ACS Nano 2014, 8 (7), 6633–6643. DOI: 10.1021/nn502058j.

(6) Kopperger, E.; List, J.; Madhira, S.; Rothfischer, F.; Lamb, D. C.; Simmel, F. C. A self-assembled nanoscale robotic arm controlled by electric fields. Science 2018, 359 (6373), 296–301. DOI: 10.1126/science.aao4284 (acccessed 2024/10/24).

(7) Mills, A.; Aissaoui, N.; Maurel, D.; Elezgaray, J.; Morvan, F.; Vasseur, J. J.; Margeat, E.; Quast, R. B.; Lai Kee-Him, J.; Saint, N.;, et al. A modular spring-loaded actuator for mechanical activation of membrane proteins. Nature Communications 2022, 13 (1), 3182. DOI: 10.1038/s41467-022-30745-2.

(8) Glembockyte, V.; Grabenhorst, L.; Trofymchuk, K.; Tinnefeld, P. DNA Origami Nanoantennas for Fluorescence Enhancement. Accounts of Chemical Research 2021, 54 (17), 3338–3348. DOI: 10.1021/acs.accounts.1c00307.

(9) Grabenhorst, L.; Pfeiffer, M.; Schinkel, T.; Kümmerlin, M.; Maglic, J. B.; Brüggenthies, G. A.; Selbach, F.; Murr, A. T.; Tinnefeld, P.; Glembockyte, V. Engineering Modular and Tunable Single Molecule Sensors by Decoupling Sensing from Signal Output. bioRxiv 2023, 2023.2011.2006.565795. DOI: 10.1101/2023.11.06.565795.

(10) He, Z.; Shi, K.; Li, J.; Chao, J. Self-assembly of DNA origami for nanofabrication, biosensing, drug delivery, and computational storage. iScience 2023, 26 (5), 106638. DOI: 10.1016/j.isci.2023.106638 From NLM.

(11) Douglas, S. M.; Bachelet, I.; Church, G. M. A Logic-Gated Nanorobot for Targeted Transport of Molecular Payloads. Science 2012, 335 (6070), 831–834. DOI: doi:10.1126/science.1214081.

(12) Amendola, V.; Meneghetti, M. Self-healing at the nanoscale. Nanoscale 2009, 1 (1), 74–88. DOI: 10.1039/b9nr00146h.

(13) Fang, W.; Xie, M.; Hou, X.; Liu, X.; Zuo, X.; Chao, J.; Wang, L.; Fan, C.; Liu, H.; Wang, L. DNA Origami Radiometers for Measuring Ultraviolet Exposure. Journal of the American Chemical Society 2020, 142 (19), 8782–8789. DOI: 10.1021/jacs.0c01254.

(14) Kielar, C.; Xin, Y.; Shen, B.; Kostiainen, M. A.; Grundmeier, G.; Linko, V.; Keller, A. On the Stability of DNA Origami Nanostructures in Low-Magnesium Buffers. Angewandte Chemie International Edition 2018, 57 (30), 9470–9474. DOI: 10.1002/anie.201802890.

(15) Ramakrishnan, S.; Shen, B.; Kostiainen, M. A.; Grundmeier, G.; Keller, A.; Linko, V. Real-Time Observation of Superstructure-Dependent DNA Origami Digestion by DNase I Using High-Speed Atomic Force Microscopy. ChemBioChem 2019, 20 (22), 2818–2823. DOI: 10.1002/cbic.201900369.

(16) Sobczak, J.-P. J.; Martin, T. G.; Gerling, T.; Dietz, H. Rapid Folding of DNA into Nanoscale Shapes at Constant Temperature. Science 2012, 338 (6113), 1458–1461. DOI: doi:10.1126/science.1229919.

(17) Xin, Y.; Piskunen, P.; Suma, A.; Li, C.; Ijäs, H.; Ojasalo, S.; Seitz, I.; Kostiainen, M. A.; Grundmeier, G.; Linko, V.;, et al. Environment-Dependent Stability and Mechanical Properties of DNA Origami Six-Helix Bundles with Different Crossover Spacings. Small 2022, 18 (18), 2107393. DOI: 10.1002/smll.202107393.

(18) Conway, J. W.; McLaughlin, C. K.; Castor, K. J.; Sleiman, H. DNA nanostructure serum stability: greater than the sum of its parts. Chemical Communications 2013, 49 (12), 1172–1174, 10.1039/C2CC37556G. DOI: 10.1039/C2CC37556G.

(19) Wang, Y.; Baars, I.; Berzina, I.; Rocamonde-Lago, I.; Shen, B.; Yang, Y.; Lolaico, M.; Waldvogel, J.; Smyrlaki, I.; Zhu, K.;, et al. A DNA robotic switch with regulated autonomous display of cytotoxic ligand nanopatterns. Nat Nanotechnol 2024. DOI: 10.1038/s41565-024-01676-4 From NLM Publisher.

(20) Gerling, T.; Kube, M.; Kick, B.; Dietz, H. Sequence-programmable covalent bonding of designed DNA assemblies. Science Advances 2018, 4 (8), eaau1157. DOI: doi:10.1126/sciadv.aau1157.

(21) Engelhardt, F. A. S.; Praetorius, F.; Wachauf, C. H.; Brüggenthies, G.; Kohler, F.; Kick, B.; Kadletz, K. L.; Pham, P. N.; Behler, K. L.; Gerling, T.;, et al. Custom-Size, Functional, and Durable DNA Origami with Design-Specific Scaffolds. ACS Nano 2019, 13 (5), 5015–5027. DOI: 10.1021/acsnano.9b01025.

(22) Brown, T. M.; Fakih, H. H.; Saliba, D.; Asohan, J.; Sleiman, H. F. Stabilization of Functional DNA Structures with Mild Photochemical Methods. Journal of the American Chemical Society 2023, 145 (4), 2142–2151. DOI: 10.1021/jacs.2c08808.

(23) Scheckenbach, M.; Schubert, T.; Forthmann, C.; Glembockyte, V.; Tinnefeld, P. Self-Regeneration and Self-Healing in DNA Origami Nanostructures. Angewandte Chemie International Edition 2021, 60 (9), 4931–4938. DOI: 10.1002/anie.202012986.

(24) Nguyen, L.; Döblinger, M.; Liedl, T.; Heuer-Jungemann, A. DNA-Origami-Templated Silica Growth by Sol–Gel Chemistry. Angewandte Chemie International Edition 2019, 58 (3), 912–916. DOI: 10.1002/anie.201811323.

(25) Liu, X.; Zhang, F.; Jing, X.; Pan, M.; Liu, P.; Li, W.; Zhu, B.; Li, J.; Chen, H.; Wang, L.;, et al. Complex silica composite nanomaterials templated with DNA origami. Nature 2018, 559 (7715), 593–598. DOI: 10.1038/s41586-018-0332-7.

(26) Jing, X.; Zhang, F.; Pan, M.; Dai, X.; Li, J.; Wang, L.; Liu, X.; Yan, H.; Fan, C. Solidifying framework nucleic acids with silica. Nature Protocols 2019, 14 (8), 2416–2436. DOI: 10.1038/s41596-019-0184-0.

(27) Ponnuswamy, N.; Bastings, M. M. C.; Nathwani, B.; Ryu, J. H.; Chou, L. Y. T.; Vinther, M.; Li, W. A.; Anastassacos, F. M.; Mooney, D. J.; Shih, W. M. Oligolysine-based coating protects DNA nanostructures from low-salt denaturation and nuclease degradation. Nature Communications 2017, 8 (1), 15654. DOI: 10.1038/ncomms15654.

(28) Agarwal, N. P.; Matthies, M.; Gür, F. N.; Osada, K.; Schmidt, T. L. Block Copolymer Micellization as a Protection Strategy for DNA Origami. Angewandte Chemie International Edition 2017, 56 (20), 5460–5464. DOI: 10.1002/anie.201608873.

(29) Bertosin, E.; Stömmer, P.; Feigl, E.; Wenig, M.; Honemann, M. N.; Dietz, H. Cryo-Electron Microscopy and Mass Analysis of Oligolysine-Coated DNA Nanostructures. ACS Nano 2021, 15 (6), 9391–9403. DOI: 10.1021/acsnano.0c10137.

(30) Nguyen, M.-K.; Nguyen, V. H.; Natarajan, A. K.; Huang, Y.; Ryssy, J.; Shen, B.; Kuzyk, A. Ultrathin Silica Coating of DNA Origami Nanostructures. Chemistry of Materials 2020, 32 (15), 6657–6665. DOI: 10.1021/acs.chemmater.0c02111.

(31) Ober, M. F.; Baptist, A.; Wassermann, L.; Heuer-Jungemann, A.; Nickel, B. In situ small-angle X-ray scattering reveals strong condensation of DNA origami during silicification. Nature Communications 2022, 13 (1), 5668. DOI: 10.1038/s41467-022-33083-5.

(32) Eklund, A. S.; Comberlato, A.; Parish, I. A.; Jungmann, R.; Bastings, M. M. C. Quantification of Strand Accessibility in Biostable DNA Origami with Single-Staple Resolution. ACS Nano 2021, 15 (11), 17668–17677. DOI: 10.1021/acsnano.1c05540.

(33) Wassermann, L. M.; Scheckenbach, M.; Baptist, A. V.; Glembockyte, V.; Heuer-Jungemann, A. Full Site-Specific Addressability in DNA Origami-Templated Silica Nanostructures. Advanced Materials 2023, 35 (23), 2212024. DOI: 10.1002/adma.202212024.

(34) Lerner, E.; Ploetz, E.; Hohlbein, J.; Cordes, T.; Weiss, S. A Quantitative Theoretical Framework For Protein-Induced Fluorescence Enhancement–Förster-Type Resonance Energy Transfer (PIFE-FRET). The Journal of Physical Chemistry B 2016, 120 (26), 6401–6410. DOI: 10.1021/acs.jpcb.6b03692.

(35) Widengren, J.; Schwille, P. Characterization of Photoinduced Isomerization and Back-Isomerization of the Cyanine Dye Cy5 by Fluorescence Correlation Spectroscopy. The Journal of Physical Chemistry A 2000, 104 (27), 6416–6428. DOI: 10.1021/jp000059s.

(36) Kozlov, A. G.; Lohman, T. M. Stopped-Flow Studies of the Kinetics of Single-Stranded DNA Binding and Wrapping around the Escherichia coli SSB Tetramer. Biochemistry 2002, 41 (19), 6032–6044. DOI: 10.1021/bi020122z.

(37) Steffen, F. D.; Sigel, R. K. O.; Börner, R. An atomistic view on carbocyanine photophysics in the realm of RNA. Physical Chemistry Chemical Physics 2016, 18 (42), 29045–29055, 10.1039/C6CP04277E. DOI: 10.1039/C6CP04277E.

(38) Ploetz, E.; Ambrose, B.; Barth, A.; Borner, R.; Erichson, F.; Kapanidis, A. N.; Kim, H. D.; Levitus, M.; Lohman, T. M.; Mazumder, A.;, et al. A new twist on PIFE: photoisomerisation-related fluorescence enhancement. Methods Appl Fluoresc 2023, 12 (1). DOI: 10.1088/2050-6120/acfb58 From NLM Medline.

(39) Maillard, J.; Klehs, K.; Rumble, C.; Vauthey, E.; Heilemann, M.; Fürstenberg, A. Universal quenching of common fluorescent probes by water and alcohols. Chemical Science 2021, 12 (4), 1352–1362, 10.1039/D0SC05431C. DOI: 10.1039/D0SC05431C.

(40) Lee, S. F.; Vérolet, Q.; Fürstenberg, A. Improved Super-Resolution Microscopy with Oxazine Fluorophores in Heavy Water. Angewandte Chemie International Edition 2013, 52 (34), 8948–8951. DOI: 10.1002/anie.201302341.

(41) Klehs, K.; Spahn, C.; Endesfelder, U.; Lee, S. F.; Fürstenberg, A.; Heilemann, M. Increasing the Brightness of Cyanine Fluorophores for Single-Molecule and Superresolution Imaging. ChemPhysChem 2014, 15 (4), 637–641. DOI: 10.1002/cphc.201300874.

(42) Maillard, J.; Rumble, C. A.; Fürstenberg, A. Red-Emitting Fluorophores as Local Water-Sensing Probes. The Journal of Physical Chemistry B 2021, 125 (34), 9727–9737. DOI: 10.1021/acs.jpcb.1c05773.

(43) Tinnefeld, P.; Buschmann, V.; Herten, D.-P.; Han, K.-T.; Sauer, M. Confocal Fluorescence Lifetime Imaging Microscopy (FLIM) at the Single Molecule Level. Single Molecules 2000, 1 (3), 215–223. DOI: 10.1002/1438-5171(200009)1:3<215::AID-SIMO215>3.0.CO;2-S.

(44) Jungmann, R.; Steinhauer, C.; Scheible, M.; Kuzyk, A.; Tinnefeld, P.; Simmel, F. C. Single-Molecule Kinetics and Super-Resolution Microscopy by Fluorescence Imaging of Transient Binding on DNA Origami. Nano Letters 2010, 10 (11), 4756–4761. DOI: 10.1021/nl103427w.

(45) Püntener, S.; Rivera-Fuentes, P. Single-Molecule Peptide Identification Using Fluorescence Blinking Fingerprints. Journal of the American Chemical Society 2023, 145 (2), 1441–1447. DOI: 10.1021/jacs.2c12561.

(46) Auvinen, H.; Zhang, H.; Nonappa; Kopilow, A.; Niemelä, E. H.; Nummelin, S.; Correia, A.; Santos, H. A.; Linko, V.; Kostiainen, M. A. Protein Coating of DNA Nanostructures for Enhanced Stability and Immunocompatibility. Advanced Healthcare Materials 2017, 6 (18), 1700692. DOI: 10.1002/adhm.201700692.

(47) Nickels, P. C.; Wünsch, B.; Holzmeister, P.; Bae, W.; Kneer, L. M.; Grohmann, D.; Tinnefeld, P.; Liedl, T. Molecular force spectroscopy with a DNA origami–based nanoscopic force clamp. Science 2016, 354 (6310), 305–307. DOI: 10.1126/science.aah5974.

(48) Schnitzbauer, J.; Strauss, M. T.; Schlichthaerle, T.; Schueder, F.; Jungmann, R. Super-resolution microscopy with DNA-PAINT. Nature Protocols 2017, 12 (6), 1198–1228. DOI: 10.1038/nprot.2017.024.

(49) Nečas, D.; Klapetek, P. Gwyddion: an open-source software for SPM data analysis. Open Physics 2012, 10 (1), 181–188. DOI: doi:10.2478/s11534-011-0096-2 (acccessed 2024-07-23).

(50) Schröder, T.; Scheible, M. B.; Steiner, F.; Vogelsang, J.; Tinnefeld, P. Interchromophoric Interactions Determine the Maximum Brightness Density in DNA Origami Structures. Nano Letters 2019, 19 (2), 1275–1281. DOI: 10.1021/acs.nanolett.8b04845.

