## Supporting Information for "Monitoring the Coating of Single DNA Origami Nanostructures with a Molecular Fluorescence Lifetime Sensor"

### 1. Methods and materials

#### 1.1. General materials

The two layer origami (TLO) was folded, purified and stored in a 1× TE buffer consisting of 10 mM Tris, 1 mM EDTA and 12 mM $MgCl_2$. The 12 helix bundle origami (12HB) was folded, purified and stored in a 1× TAE buffer consisting of 40 mM Tris, 20 mM acetic acid, 1 mM EDTA, and 16 mM $MgCl_2$.

The p8064 scaffold strand for the folding of the DNA Origami nanostructures were extracted from M13mp18 bacteriophages (produced in- house). Unmodified staple strands were purchased from Eurofins Genomics GmbH and Integrated Device Technology Inc. Dye labeled oligonucleotides for DNA PAINT imaging or permanent labeling were purchased from Eurofins Genomics GmbH (Germany).

Specific materials used for individual experiments are described in the sections below.

#### 1.1. DNA Origami folding

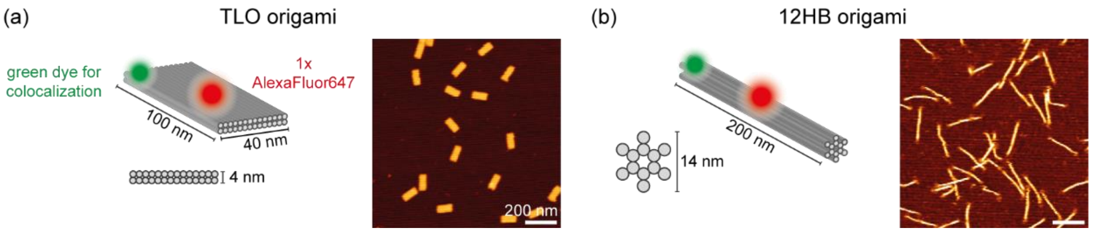

**Figure S1.** Schemes of the DNA origami nanorulers used in this study. Exemplary schemes and AFM scans of TLO (a) and 12HB (b) as rigid DNA platforms.[1] Scale bars represent 200 nm.

All investigated TLO DNA origami nanostructures (depicted in Figure S1) were folded in a 1× TE buffer containing 12 mM $MgCl_2$ using the corresponding p8064 scaffold strand extracted from M13mp18 bacteriophages realized with a linear thermal annealing ramp from 60°C to 44°C with 1h/°C after an initial 65°C denaturation step. Modifications of the DNA origami were realized using caDNAno (version 2.2.0).[2] A full list of the unmodified staple strands and sequences of the TLO DNA origami is given in Table S13, a list of modified staple strands is given in Table S2 (sensor labels and colocalization label) and Table S4 (DNA PAINT docking sites). A caDNAno design file is given in Figure S17.

The 12HB DNA origami nanostructures (depicted in Figure S1) were folded in a 1× TAE buffer containing 16 mM $MgCl_2$ using the same scaffold strand as for the TLO. The structures were folded with a non-linear thermal annealing ramp starting at 65 °C and then cooling down to 4 °C over a period of 25 hours.[3] Modifications of the DNA origami were realized using caDNAno (version 2.2.0). A full list of the unmodified staple strands and sequences of the 12HB DNA origami is given in Table S14, a list of modified staple strands is given in Table S3. A caDNAno design file is given in Figure S18.

For the folding of the DNA origami nanostructures, the scaffold strand and the staple strands were mixed as given in Table S1 in the corresponding 1× folding buffer. Unmodified core staple strands, which are completely incorporated in the origami structure, were used in 10-fold excess with respect to the scaffold strand. Fluorescently labeled staple strands with dye modifications at the 3'-ends were used in a 30-fold excess with respect to the scaffold strand. Staple strands with protruding 3'-ends, which act as docking sites for DNA PAINT or labeling experiments, were used in 30-fold excess with respect to the scaffold strand. Biotinylated staple strands, which were incorporated for surface immobilization of the DNA origami structures, were used in a 30-fold excess with respect to the scaffold strand. Folding mixes had a total volume of 100 µl with final concentrations of scaffold strand, core staple strands and modified staple strands (biotinylated, DNA PAINT docking sites) as given in Table S1.

**Table S1.** Final concentrations and relative equivalents of scaffold strand, unmodified staple strands (core staple strands) and modified staple strands (e.g., biotinylated staple strands for immobilization, cyanine dye modification for fluorescence lifetime imaging and DNA PAINT docking site staple strands for super-resolution imaging) used within this study.

| Reagent | Final concentration [nM] | Equivalents |
|---|---|---|
| Scaffold strand | 20 | 1 |
| Core staple strands | 200 | 10 |
| Dye modified staple strands | 600 | 30 |
| Docking site staple strands | 600 | 30 |
| Biotinylated staple strands | 600 | 30 |

Folded DNA origami nanostructures were purified by filtration using Amicon Ultra filters (100 K, Merck, Germany). The filter was first centrifuged with folding buffer for 7 minutes at 6000 g. The sample solution was then loaded into the filter and centrifuged for 15 minutes at 6000 g. 300 µL of folding buffer was loaded into the filter and centrifuged for 15 minutes at 6000 g, which was repeated. After three washing steps, the filter was inverted and placed into a new collection tube. The purified sample could then be collected by centrifugation for 2 minutes at 1000 g.

Concentrations of purified sample solution were measured via UV/vis spectroscopy (NanoDrop, Fischer Scientific, USA).

#### 1.2. Labeling of DNA origami with fluorescence lifetime sensor

To probe the encapsulation process of DNA origami by either PLL-PEG or silica, the environment dependent cyanine fluorophore AlexaFluor647 was labeled at different positions on the TLO and the 12HB DNA origami (see Tables S2 and S3). To screen the encapsulation process at the surface of the TLO and at the interface of the two DNA layers, the AlexaFluor647 modification was placed in the central area of the DNA origami at the 3' end of a staple pointing out of the upper DNA layer (surface) or into the interface between the two DNA layers (interface). For a modular and cheaper labeling strategy, a 21-nt external DNA docking site was placed at the central surface position enabling external labeling by the addition of a complementary 21-nt strand, that is labeled with AlexaFluor647 at the 3'-end (5'-GTGATGTAGGTGGTAGAGGAAT-3'). To screen the encapsulation at the corner of the TLO origami, the same external label was placed at a corner of the DNA origami. To screen the encapsulation of the 12HB, the external label was placed at a central position of the 12HB axis. For colocalizing purposes, a green Atto542 label was introduced into the TLO origami design, while a green Cy3 label was incorporated into the 12HB origami.

**Table S2** Modified staple strands of the TLO FLIM sensor. Sequences are denoted from 5'- to 3'-end. Internal dye modifications (surface and interface AlexaFluor647, Atto542 colocalization label) are labelled to the 3'-end of the corresponding staple strand (marked in red and green, respectively). The docking site staple strands for external labelling (external central and edge AlexaFluor647) exhibit a 21 nt long docking site, marked in red, on the 3'-end. The numbers for the 5'- end 3'-end of the staples represent the helix number in the corresponding caDNAno file. Number in brackets represent the starting and ending position of the staple in the corresponding helix.

| Name | Docking Site Length (nt) | Sequence (5` to 3`) | 5'-end | 3'-end |
|---|---|---|---|---|
| Surface | X | AGGCTATCGAGAATCGTAACAACCTTGACCGT-AlexaFluor647 | 13[136] | 16[136] |
| Interface | X | TGATGCAGGGAACAAATTAAGTAAACAAACATCAA-AlexaFluor647 | 16[167] | 17[172] |
| External central | 21 | AGGCTATCGAGAATCGTAACAACCTTGACCGT-TTCCTCTACCACCTACATCAC | 13[136] | 16[136] |
| External corner | 21 | TGAGTTTTATTTCGGATAAACACCGCCACC-TTCCTCTACCACCTACATCAC | 1[296] | 4[296] |
| Atto542 colocalization | X | TGCGGATGTAGCTCAATTAAGCAAGTACCAAA-Atto542 | 9[136] | 12[136] |

**Table S3** Modified staple strands of the 12HB FLIM sensor. Sequences are denoted from 5'- to 3'-end. The docking site staple strands for external labelling (external central and edge AlexaFluor647) exhibit a 21 nt long docking site, marked in red, on the 3'-end. Internal dye modifications (Cy3 colocalization label) are labelled to the 3'-end of the corresponding staple strand (marked in green). The numbers for the 5'- end 3'-end of the staples represent the helix number in the corresponding caDNAno file. Number in brackets represent the starting and ending position of the staple in the corresponding helix.

| Name | Docking Site Length (nt) | Sequence (5` to 3`) | 5'-end | 3'-end |
|---|---|---|---|---|
| External central | 21 | TCGTTCACCGCCTGGCCCT -TTCCTCTACCACCTACATCAC | 10[331] | 11[344] |
| Cy3 colocalization | X | GTTTGAGGGGACCTCATTTGCCG -Cy3 | 4[125] | 4[103] |

#### 1.3. DNA PAINT docking sites

To simultaneously probe the encapsulation and structural integrity of the TLO origami, DNA PAINT docking sites were incorporated into the TLO with an internal AlexaFluor647 label at the central surface position. By placing three DNA PAINT docking sites on each of the four corners of the TLO surface, the overall shape of the TLO origami can be investigated with nanometer resolution (see Figure S11). The in 4×3 DNA PAINT docking site modified staple strands are given in Table S4 and carry the 8 nt long docking site at their 3'-end.

**Table S4** Modified staple strands of the TLO DNA PAINT nanoruler. Sequences are denoted from 5'- to 3'-end. The DNA PAINT docking site staple strands exhibit a triple T linker and an 8 nt long docking site, marked in green, on the 3'-end. The numbers for the 5'- end 3'-end of the staples represent the helix number in the corresponding caDNAno file. Number in brackets represent the starting and ending position of the staple in the corresponding helix.

| Name | Docking Site Length (nt) | Sequence (5` to 3`) | 5'-end ´ | 3'-end] |
|---|---|---|---|---|
| TLO Spot 1_1 | 8 | GATAGTTGCGCCGACACTAAAACGCAAGCGCGACCCAAAT-TTT- AACATTCC | 1[32] | 4[40] |
| TLO Spot 1_2 | 8 | GCACCAACATGACAACTCGGTTTATCAGCTTG-TTT-AACATTCC | 2[55] | 0[40] |
| TLO Spot 1_3 | 8 | TGCAGATAACACCAGATATTCATTAAACAAAG-TTT-AACATTCC | 6[55] | 3[55] |
| TLO Spot 2_1 | 8 | TCAGAGCCGATTAGGATGATACAGCCAGAGCC-TTT-AACATTCC | 1[232] | 4[232] |
| TLO Spot 2_2 | 8 | AAGAGAAGACCACCCTACGATCTAAAGTTTTG-TTT-AACATTCC | 2[247] | 0[232] |
| TLO Spot 2_3 | 8 | CAGCAAAACGTTTGCCAACCACCAGAGTGTAC-TTT-AACATTCC | 6[247] | 3[247] |
| TLO Spot 3_1 | 8 | TGGTGCTGACTGGTGTCGGTGCCCTCCGCTCA-TTT-AACATTCC | 21[40] | 24[40] |
| TLO Spot 3_2 | 8 | CCTGGGGTCACGCTGGCCCTTATAAATCAAAA-TTT-AACATTCC | 25[40] | 27[55] |
| TLO Spot 3_3 | 8 | AAGCGGTCGCCTAATGAATTGTTACCTGCATC-TTT-AACATTCC | 26[55] | 23[55] |
| TLO Spot 4_1 | 8 | GAATTGAGGAGGTGAGCAGAGATAATCCAGAA-TTT-AACATTCC | 21[232] | 24[232] |
| TLO Spot 4_2 | 8 | TGTTTTTAGCGCTTAAATCGGAACCCTAAAGG-TTT-AACATTCC | 25[232] | 27[247] |
| TLO Spot 4_3 | 8 | CACCCGCCTAATCAGTCTGGTAATGAACCCTT-TTT-AACATTCC | 26[247] | 23[247] |

#### 1.4. AFM imaging with JPK Nanowizard

For probing correct folding of the origami structures and probing the encapsulation by silica and PLL-PEG, AFM images were taken. AFM scans in aqueous solution (AFM buffer = 40 mM Tris, 2 mM EDTA, 12.5 mM $Mg(OAc)_2 \cdot 4\ H_2O$) were realized on a NanoWizard® 3 ultra AFM (JPK Instruments AG). For sample immobilization, a freshly cleaved mica surface (Quality V1, Plano GmbH) was incubated with 10 mM solution of Poly-L-ornithine (0.01% 30000 – 70000 g/mol, Sigma Aldrich) for 3 minutes. The mica was washed three times with ultra-pure water to get rid of unbound Poly-L-ornithine and blow-dried with air. The dried mica surface was incubated with 1 nM sample solution for 3 minutes and washed with AFM buffer three times. Measurements were performed in AC mode on a scan area of 3 x 3 µm with a micro cantilever ($v_{res}$ = 110 kHZ, $k_{spring}$ = 9 N/m, Olympus Corp.).

Leveling, background correction and extraction of height histograms of obtained AFM images were realized with the software Gwyddion (version 2.60).[4]

#### 1.5. Surface-Immobilization of DNA nanostructures

High precision µm microscope cover glass (170 µm, 22×22 mm, No. 1.5H glass slides, Carl Roth GmbH, Germany) were initially ultrasonicated in a 1% Hellmanex solution. After thoroughly washing with ultra-pure water, the glass slides were irradiated for 30 min in a UV ozone cleaner (PSD-UV4, Novascan Technologies, USA). Cleaned glass slides and microscope slides were assembled into an inverted flow chamber as described previously.[5] The assembled chambers were rinsed with 1× PBS, and passivated with 50 µL of BSA-biotin (0.5 mg/mL in PBS, Sigma Aldrich, USA) for 15 minutes and washed with 50 µL 1× PBS. The passivated surfaces were incubated with 50 µL Neutravidin (0.25 mg/mL-in 1× PBS, Sigma Aldrich, USA) or 50 µL Streptavidin (0.5 mg/mL in 1xPBS, Sigma Aldrich, USA) for 15 minutes and washed with 50 µL 1× PBS. The sample solution with DNA origami featuring several staple strands with biotin modifications on the base was diluted to approximately 50 pM in 1× PBS buffer containing 500 mM NaCl and incubated in the chambers for ca. 5 minutes and stored in a 1xTAE containing 10 mM $MgCl_2$. Sufficient surface density was probed with a TIRF microscope.

#### 1.6. Coating with PLL-PEG or silica

The PLL-PEG block copolymer K10PEG (1K) was purchased from Alamanda polymers ($mPEG_{1K}$-b-$PLKC_{10}$) and dissolved in ultra-pure water at a concentration of 2 mM. [6, 7] Aliquots were stored at -20 °C and thawed and ultrasonicated for 10 min before usage. To coat immobilized DNA origami nanostructures, the 2 mM PLL-PEG solution was diluted in a 1xTAE buffer containing 10 mM $MgCl_2$ to a final concentration of 20 µM. 50 µL of the prepared solution were incubated in the sample chambers for 30 min. The coated sample chambers were washed with 50 µL of 1xTAE 10 mM $MgCl_2$. To decomplex the cationic PLL-PEG coating from the DNA origami, a 20 µM solution of anionic dextran sulfate (Sigma Aldrich, M = 20 000 g/mol)[6] in 1xTAE 10 mM $MgCl_2$ was incubated in the coated sample chambers for 30 min and washed with 50 µL of 1xTAE 10 mM $MgCl_2$.

For the silicification of immobilized DNA origami an adapted version of the protocol of Fan and co-workers was applied.[1, 8] Initially, a precursor solution was prepared by adding 5 mL of 1×TAE buffer (40 mM Tris, 2mM EDTA, 12.5 mM $MgAc_2$, pH=8.0) to a 10 mL glass bottle with a suitably-size magnet and then slowly adding 100 µL of TMAPS (50% (wt/wt) in methanol, TCI America). This solution was stirred vigorously for 20 min at room temperature. After that, 100 µL of TEOS (98%, Sigma Aldrich) were slowly added and the resulting solution was again stirred for 20 min at room temperature. 50 µL of the precursor solution was incubated in sample chambers for 24 h. The coated sample chambers were washed with 100 µL 80% ethanol and with 100 µL ultra-pure water. The samples were then stored in 1xTAE 10 mM $MgCl_2$.

For AFM imaging, mica slides with immobilized DNA origami were analogously incubated with either the 20 µM PLL-PEG solution or with the silica precursor solution.

#### 1.7. Photostabilization of fluorescent labels

Fluorescence lifetime imaging microscopy (FLIM) with AlexaFluor647 as imager fluorophore was carried out under photostabilizing conditions. For oxygen removal and triplet state quenching, a 1× TAE buffer with 10 mM $MgCl_2$, 12 mM 3,4-dihydroxybenzoic acid (PCA, Sigma Aldrich), 2 mM oxidized Trolox/Trolox-quinone mixture and 56 µM protocatechuate 3,4-dioxygenase (PCD, from pseudomonas sp., Sigma Aldrich) was prepared as described elsewhere.[9, 10] The flow chambers with surface immobilized origami sample were completely filled with photostabilizing buffer and sealed with a two-component glue to prevent oxygen solvation. The first measurements were carried out at least 15 minutes after introducing the oxygen removal system to allow the equilibration of the oxygen concentration in the sample solution.

##### 1.8. Fluorescence lifetime imaging

Fluorescence lifetime imaging microscopy (FLIM) was performed on a home-built confocal microscope based on an Olympus IX-71 inverted microscope as described previously.[11] AlexaFluor647 modifications labeled to surface-immobilized DNA origami were excited by a pulsed 640 nm excitation at a repetition rate of 40 MHz. The setup was controlled by a commercial software package (SymPhoTime64, PicoQuant GmbH).

For FLIM imaging with photostabilization, the laser was set to a power of 10 µW before the excitation dichroic mirror, FLIM imaging without photostabilization was performed at an excitation power of 2 µW. FLIM scans were acquired with a scan size of 200x200 px, a pixel size of 100 nm and a dwell time of 2 ms in monodirectional mode.

FCS studies were performed under photostabilizing conditions and with an excitation power set to 2 µW. After scanning an overview FLIM map as described above, individual Alexa647 labels were picked and time traces were acquired for up to 45 seconds per spot. Acquired time traces were autocorrelated using a commercial software package (SymPhoTime64, PicoQuant GmbH).

##### 1.9. Degradation studies of bare and coated DNA origami

To probe the stability of coated and bare DNA origami in degrading conditions, either a low-salt buffer or a DNAse solution were incubated on immobilized nanostructures.

Magnesium ion free conditions were realized by incubation a 1× TAE solution without magnesium on immobilized nanostructures. After 30 minutes of incubation, the samples were washed and stored in 1× TAE containing 10 mM $MgCl_2$.

Enzymatic degradation of immobilized DNA origami was tested by incubation in a DNAse solution (1:10 dilution of DNase I (1 U/µl) in 1x TAE containing 10 mM $MgCl_2$, Thermo Scientific). After 30 min of incubation, the samples were washed and stored in a 1× TAE containing 10 mM $MgCl_2$.

##### 1.10. FLIM analysis

Acquired FLIM scans (.ptu files) were exported from the acquisition software (SymPhoTime64, PicoQuant GmbH) and further analysed with a custom-written Python software.

First, individual spots representing single DNA origami nanostructures were picked by an intensity threshold. In every pick area, for each pixel the TCPSC decay was re-centered to 0 ns to subtract offset from varying excitation pulse positions in the TCPSC histograms. The average arrival time of each pixel was estimated by the median arrival time divided by ln2. To obtain spot-integrated fluorescence lifetimes for each pick area, the average photon arrival times of each pixel were weighted by their photon counts and averaged. In order to obtain absolute fluorescence lifetime values, the fluorescence lifetime decay of all photons from an individual pick were reconvoluted with an IRF decay curve, measured on the same day as the investigated sample.

#### 1.11. DNA PAINT imaging and analysis

DNA-PAINT measurements on the TLO with 4×3 8 nt DNA-PAINT docking site were carried out on a commercial Nanoimager S (ONI Ltd., UK). Red excitation at 640 nm was realized with a 1100 mW laser, green excitation at 532 nm with a 1000 mW laser, respectively. The microscope was set to TIRF illumination and a pixel size of 117 nm. In order to not corrupt the first acquired frames by photobleaching, the objective was first focused into the sample plane on a random section of the glass surface and the auto focus was activated. Subsequently the imaging lasers were shut off. Before starting measurements, the sample slide was moved to a new region of interest while still being kept in focus by the auto focus. The data acquisition was initialized by activating the lasers and taking frames of 100 ms over a user defined acquisition protocol.

All DNA-PAINT measurements were conducted at ca. 2.6 kW/cm$^2$ at 532 nm in TIRF illumination with an exposure time of 100 ms and 12,000 frames over 20 min. For colocalization, the TLO were previously imaged at ca. 100 W/cm$^2$ at 640 nm with an exposure time of 100 ms and 10 frames over 1 second. For imaging, a 1× PBS buffer containing 500 mM NaCl and an imager concentration of 5 nM was used. The 8 nt imager oligonucleotide with a Cy3B label on the 3'-end was purchased from Eurofins Genomics GmbH (Germany) and consisted of the sequence 5'-GGAATGTT-3'.

Acquired DNA-PAINT raw data were analyzed using the Picasso software package.[5] The obtained TIF-movies were first analyzed with the "localize" software. For fitting the centroid position information of single point spread functions (PSF) of individual imager strands, the MLE (maximum likelihood estimation) analysis was used with a minimal net gradient of 5000 and a box size of 5. The fitted localizations were further analyzed with the "render" software in Picasso. X-Y-drift of the localizations was corrected with the RCC drift correction. For further analysis, individual DNA origami nanostructures were picked with a pick diameter of 1.2 camera pixels. The corresponding pick region statistics such as binding kinetics were exported for further analysis. The picked DNA origami nanostructures were further aligned with the "average" module in Picasso (oversampling = 40, 10 iterations). For overview, aligned structures were rendered in a 2D grid.

#### 2. Supplementary Figures and Notes

**Table S5.** Central fluorescence lifetime values for internal and external AF647 labels on TLO or 12HB DNA. For every sample, the central fluorescence lifetime values and standard deviations of the Gaussian fit distributions are given for no coating, after coating with PLL-PEG, after uncoating the PLL-PEG coating with dextran sulfate, and after coating with silica.

| | | | Fitted fluorescence lifetime populations (ns) | | | |
|---|---|---|---|---|---|---|
| **Sample** | **Coating** | **Sample size** | $\tau_1$ | $\sigma_1$ | $\tau_2$ | $\sigma_2$ |
| Int. AF647 on TLO surface | uncoated | 752 | 1.08 | 0.05 | | |
| | PLL-PEG | 1420 | 1.33 | 0.07 | 1.57 | 0.18 |
| | Dextran sulfate | 511 | 1.08 | 0.05 | | |
| | $SiO_2$ | 529 | 1.29 | 0.09 | 1.61 | 0.27 |
| Int. AF647 at TLO interface | uncoated | 1082 | 1.22 | 0.06 | 1.37 | 0.19 |
| | PLL-PEG | 960 | 1.56 | 0.18 | 1.65 | 0.05 |
| | Dextran sulfate | 711 | 1.25 | 0.06 | 1.38 | 0.15 |
| | $SiO_2$ | 1659 | 1.56 | 0.11 | 1.76 | 0.23 |
| Ext. AF647 at TLO surface center | uncoated | 2202 | 1.32 | 0.07 | | |
| | PLL-PEG | 2004 | 1.55 | 0.08 | | |
| | Dextran sulfate | 870 | 1.32 | 0.08 | | |
| | $SiO_2$ | 1883 | 1.60 | 0.19 | | |
| Ext. AF647 at TLO surface corner | uncoated | 1744 | 1.41 | 0.09 | | |
| | PLL-PEG | 1575 | 1.58 | 0.09 | | |
| | $SiO_2$ | 810 | 1.55 | 0.28 | | |
| Ext. AF647 on 12HB | uncoated | 1499 | 1.13 | 0.05 | | |
| | PLL-PEG | 663 | 1.39 | 0.10 | 1.52 | 0.23 |
| | $SiO_2$ | 1261 | 1.37 | 0.15 | 1.76 | 0.20 |

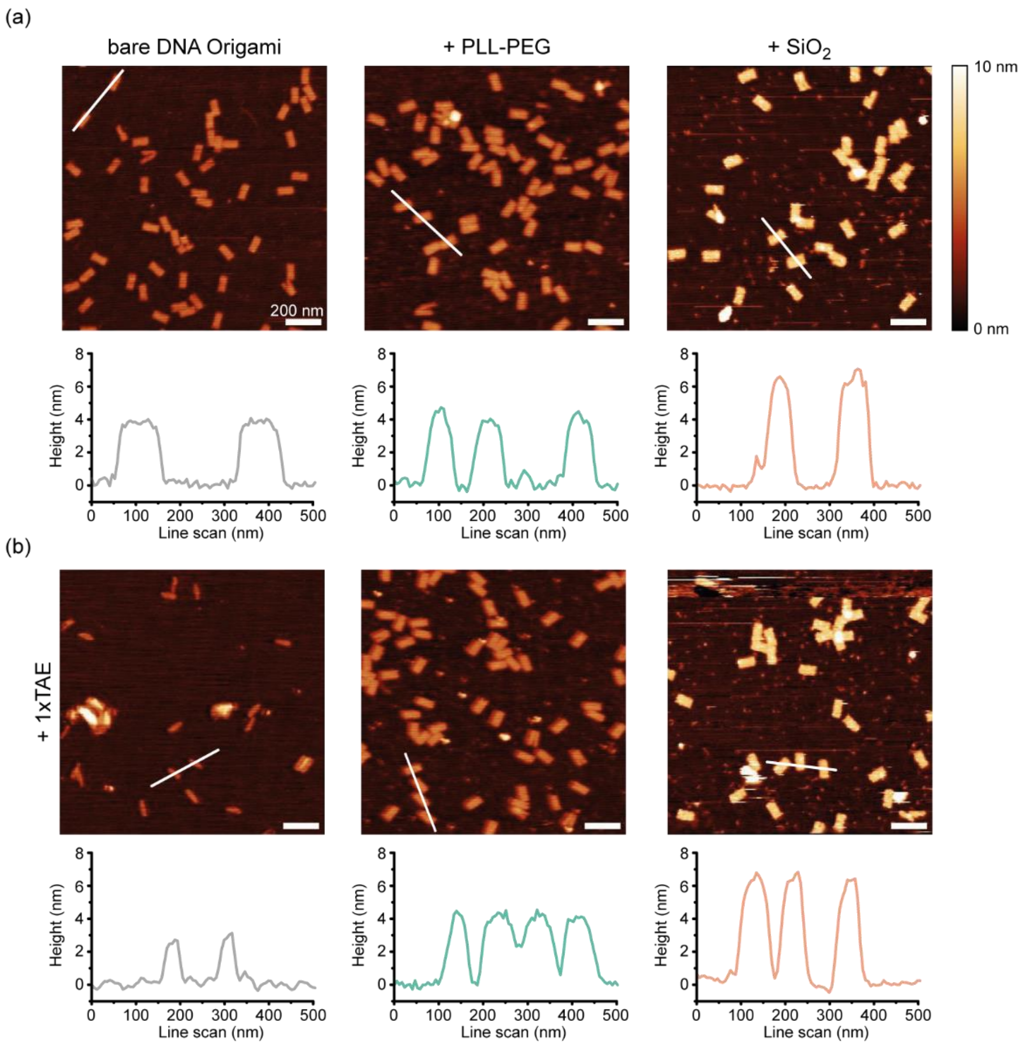

**Figure S2.** AFM characterization of TLO coated TLO with silica and PLL-PEG. A) AFM scans of bare TLO (left), after coating with PLL-PEG (middle) and after coating with silica (right). Below: Exemplary extracted line scan profiles (white lines in the AFM scans) B) AFM scans of bare TLO (left), PLL-PEG coated (middle) and silica coated (right) after incubation in 1× TAE 0 mM $MgCl_2$. Below: Exemplary extracted line scan profiles (white lines in the AFM scans). Scale bars represent 200 nm.

AFM characterization of PLL-PEG coated DNA origami has not been achieved so far, since structures coated in solution lose their charge and by that their affinity to mica surfaces prepared with either a metal cation (e.g., $Ni^{2+}$) or a cationic polymer such as poly-L-ornithine.

Exemplary AFM scans of immobilized TLO reveal the designed rectangular shape. While the encapsulation with silica leads to a measurable height increase of around 2 nm, PLL-PEG-coated TLO exhibit similar heights as bare TLO of around 4 nm. The soft nature of the PEG encapsulation layer is obviously not detectable by AFM, even though the gentle tapping mode was used for imaging. Incubation with a magnesium-free buffer (1× TAE 0 mM $MgCl_2$) however leads to collapse of the bare TLO, while the PLL-PEG-coated and silica-coated TLO nanostructures stay intact, indicating the successful coating of the DNA origami.

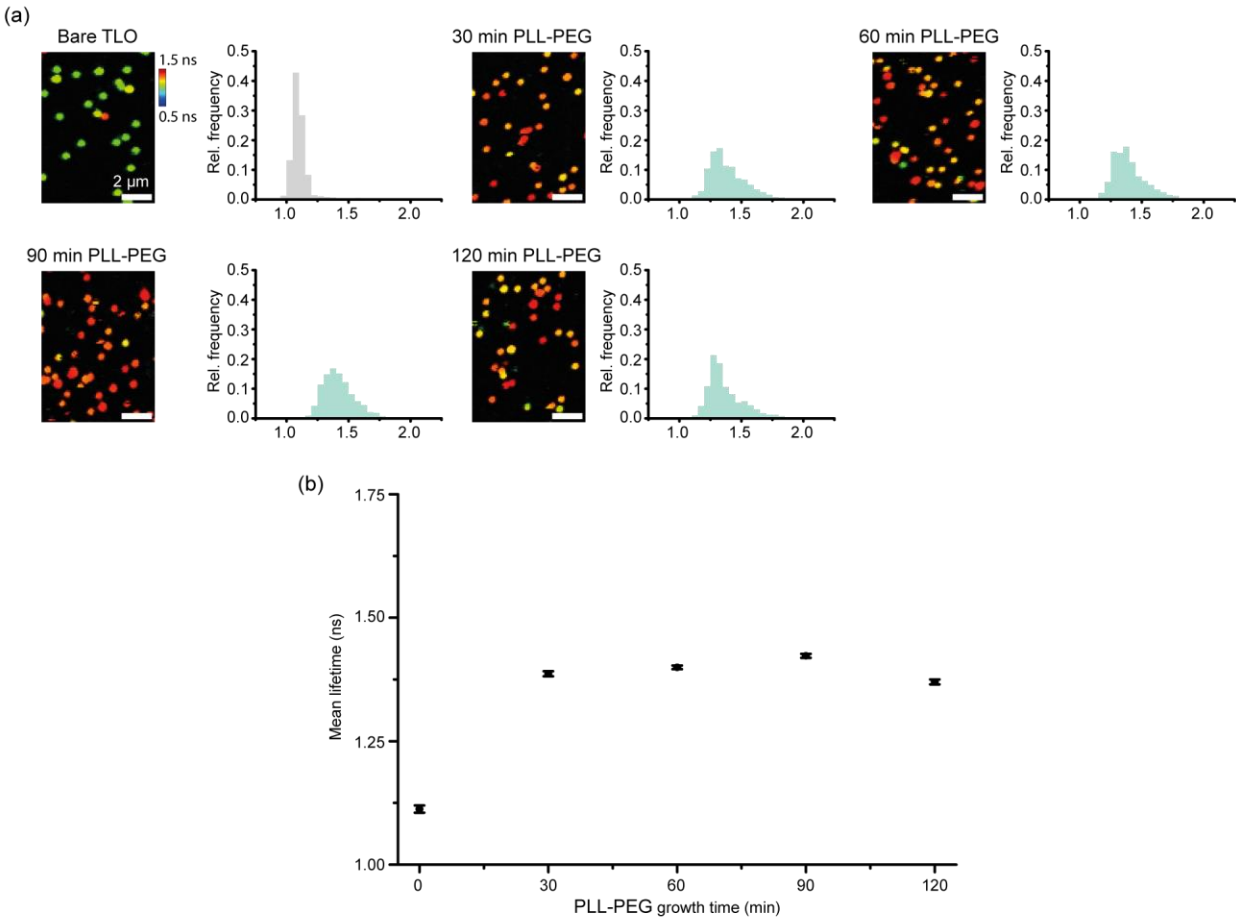

**Figure S3**. Incremental FLIM shift induced by PLL-PEG coating over time. a) FLIM scans of bare TLO and after the addition of PLL-PEG solution over varying times with corresponding spot-integrated lifetimes. b) Average lifetimes over PLL-PEG incubation time. Scale bar is 2 µm.

**Table S6.** Incremental mean spot-integrated fluorescence lifetime values and standard errors of the mean for PLL-PEG coating over time.

| Time (min) | Sample size | Mean $\tau_f$ (ns) | S.E.M. (ns) |
|---|---|---|---|
| 0 | 628 | 1.11 | 0.01 |
| 30 | 882 | 1.39 | 0.01 |
| 60 | 1423 | 1.40 | 0.00 |
| 90 | 1394 | 1.42 | 0.00 |
| 120 | 980 | 1.37 | 0.01 |

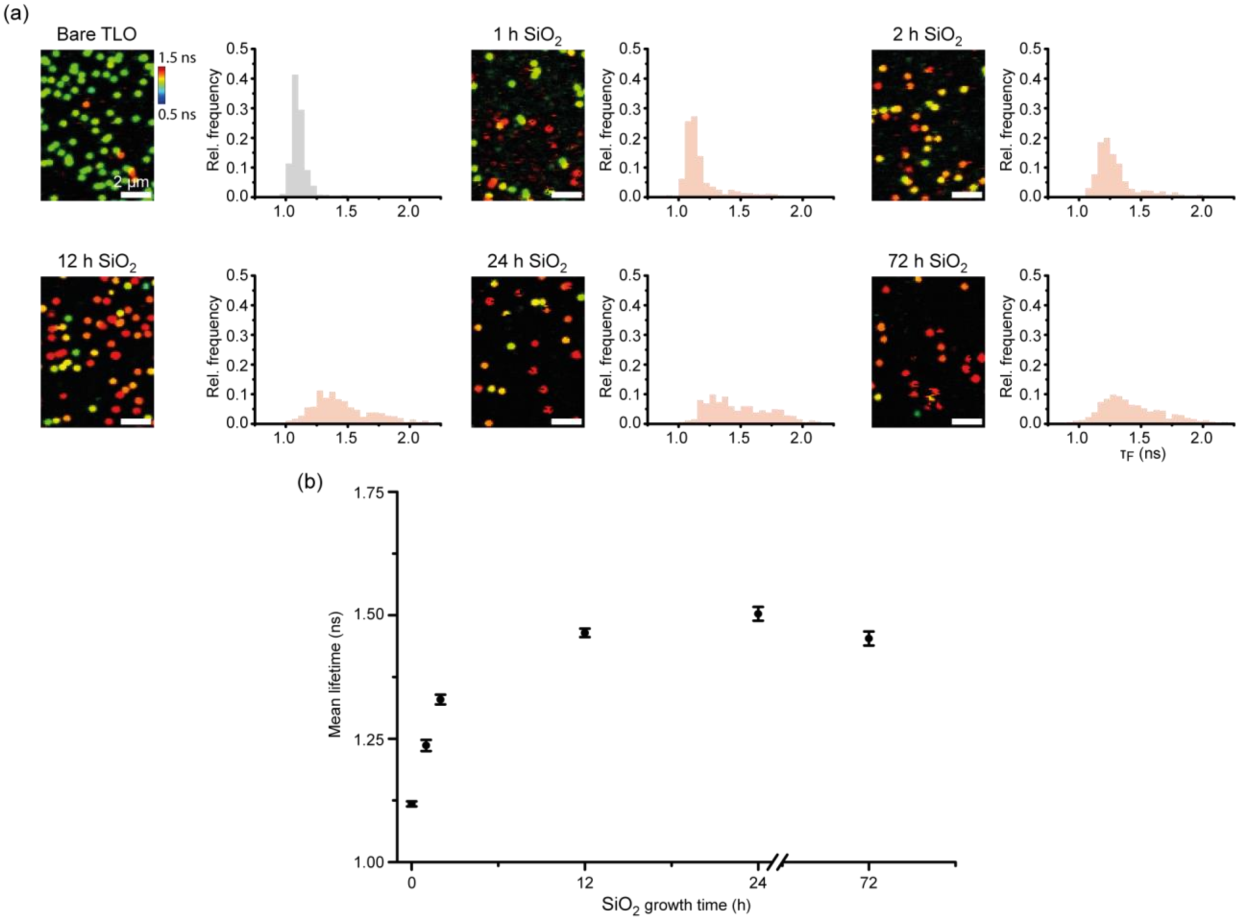

**Figure S4.** Incremental FLIM shift induced by silica coating over time. a) FLIM scans of bare TLO and after the addition of silica precursor solution over varying times with corresponding spot-integrated lifetimes. b) Average lifetimes over silica precursor incubation time. Scale bar is 2 µm.

**Table S7**. Incremental mean spot-integrated fluorescence lifetime values and standard errors of the mean for silica coating over time.

| Time (h) | Sample size | Mean $\tau_f$ (ns) | S.E.M. (ns) |
|---|---|---|---|
| 0 | 913 | 1.12 | 0.01 |
| 1 | 985 | 1.24 | 0.01 |
| 2 | 933 | 1.33 | 0.01 |
| 12 | 936 | 1.46 | 0.01 |
| 24 | 527 | 1.50 | 0.01 |
| 72 | 755 | 1.45 | 0.01 |

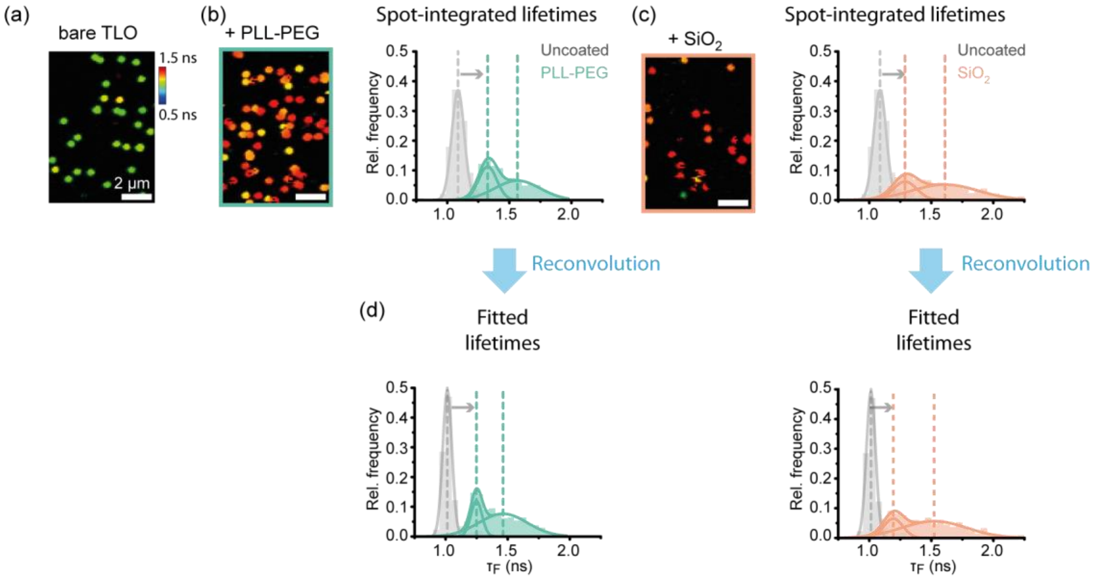

**Figure S5.** Reconvolution and fluorescence lifetime fitting compared to spot-integrated fluorescence lifetime distributions. A) Exemplary FLIM scans of bare TLO, PLL-PEG coated TLO and silica coated TLO and corresponding spot-integrated lifetime histograms. B) Lifetime histograms after spot-wise reconvolution and lifetime fitting. Scale bar is 2 µm.

Exemplary FLIM scans of bare, PLL-PEG coated and silica coated TLO as obtained from the acquisition software (SymPhoTime64, PicoQuant GmbH). After picking individual spots with a custom written Python code (see section 1.10), spot-integrated lifetimes can be extracted and visualized in a histogram, revealing a similar shift in the lifetime for both PLL-PEG and silica coating. To obtain absolute fluorescence lifetimes, the picked FLIM data were spot-wise re-convoluted with the measured IRF of the confocal setup and the lifetime decay fitted. After re-convolution and fitting, PLL-PEG and silica coating still show a similar shift in lifetime. While the absolute shift in the fluorescence lifetimes remained very comparable, the relative shift increased (fitted values in Table S8). The comparable significant shift in spot-wise integrated lifetimes highlights the potential fast readout of the cyanine based FLIM sensor for probing the successful encapsulation process within minutes and without complicated data analysis.

**Table S8**. Re-convolution and fluorescence lifetime fitting compared to spot-integrated fluorescence lifetime distributions. For every sample, the central fluorescence lifetime values and standard deviations of the Gaussian fit distributions are given for no coating, after coating with PLL-PEG, and after coating with silica.

| | | Spot-integrated fluorescence lifetime (ns) | | | | Re-convoluted fluorescence lifetime (ns) | | | |
|---|---|---|---|---|---|---|---|---|---|
| **Coating** | **Sample size** | $\tau_1$ | $\sigma_1$ | $\tau_2$ | $\sigma_2$ | $\tau_1$ | $\sigma_1$ | $\tau_2$ | $\sigma_2$ |
| uncoated | 752 | 1.08 | 0.05 | | | 1.01 | 0.04 | | |
| PLL-PEG | 1420 | 1.33 | 0.07 | 1.57 | 0.18 | 1.25 | 0.04 | 1.46 | 0.19 |
| $SiO_2$ | 529 | 1.29 | 0.09 | 1.61 | 0.27 | 1.19 | 0.08 | 1.52 | 0.26 |

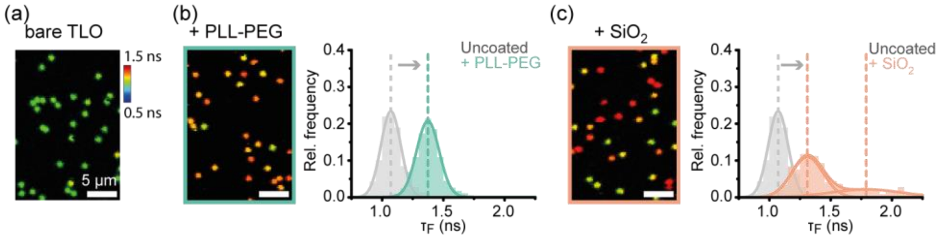

**Figure S6.** Fluorescence lifetime shift of AF647 induced by PLL-PEG or silica coating without photostabilization. a) Exemplary FLIM scans acquired without photostabilization on uncoated TLO (a), PLL-PEG coated TLO (b) and silica coated TLO (c) and corresponding spot-integrated fluorescence lifetime distributions. Scale bar is 2 µm.

Exemplary FLIM scans of bare, PLL-PEG coated and silica coated TLO as obtained from the acquisition software (SymPhoTime64, PicoQuant GmbH) and measured without photostabilization. Histograms of the spot-wise integrated lifetimes exhibit comparable shifts for the PLL-PEG and silica coating to when measured with photostabilization (fitted values in Table S9). Being able to probe the coating process even in aerobic conditions without further buffer preparation enables a fast and easy probing of the coating and degradation processes in real-time.

**Table S9**. Central fluorescence lifetime values and standard deviations of the Gaussian fit distributions measured without photostabilization before coating, after coating with PLL-PEG and after coating with silica.

| | | Fitted fluorescence lifetime populations (ns) | | | |
|---|---|---|---|---|---|
| **Coating** | **Sample size** | **$\tau_1$** | **$\sigma_1$** | **$\tau_2$** | **$\sigma_2$** |
| **uncoated** | 676 | 1.07 | 0.08 | | |
| **PLL-PEG** | 623 | 1.37 | 0.09 | | |
| **$SiO_2$** | 571 | 1.31 | 0.13 | 1.79 | 0.29 |

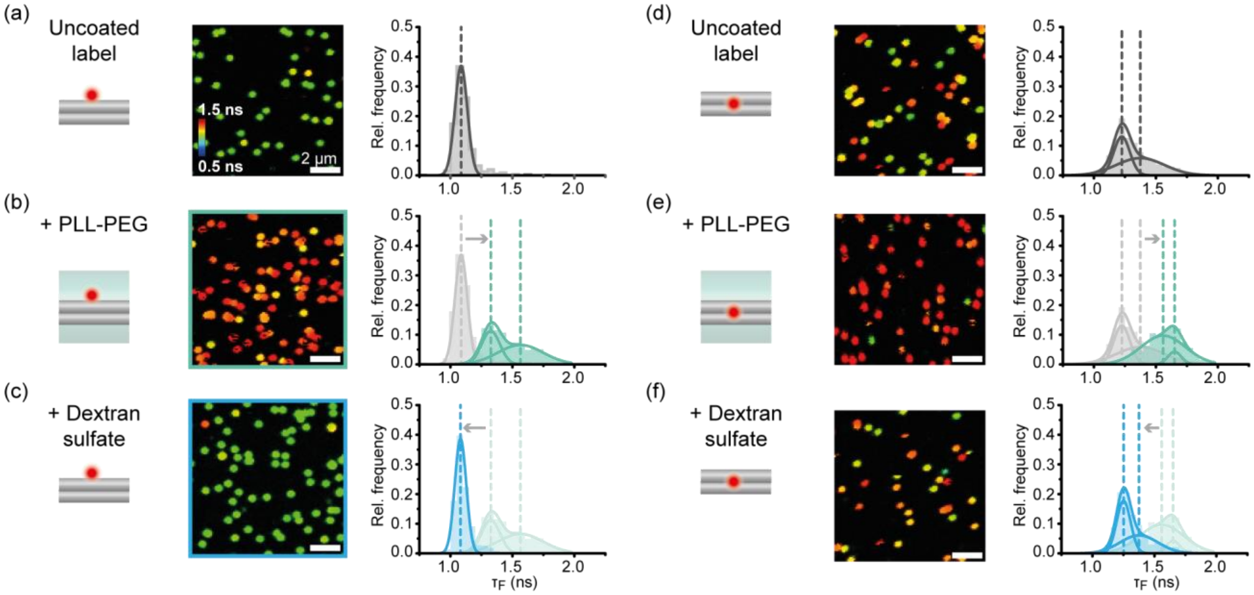

**Figure S7.** Probing the reversible coating and uncoating with PLL-PEG and dextran sulfate on TLO nanostructures with internal AF647 labels. a) FLIM scan and spot-integrated fluorescence lifetime distribution for uncoated TLO with internal AF647 label at the DNA surface. b) FLIM scan and spot-integrated fluorescence lifetime distribution for TLO with internal AF647 label at the DNA surface after PLL-PEG coating. c) FLIM scan and spot-integrated fluorescence lifetime distribution for previously PLL-PEG coated TLO with internal AF647 label at the DNA surface after de-complexing with dextran sulfate. d) to f) FLIM scans and spot-integrated fluorescence lifetime distributions for TLO with internal AF647 label at the DNA interface inside the DNA origami before coating, after coating with PLL-PEG and after the subsequent decomplexation with dextran sulfate. Scale bar is 2 µm.

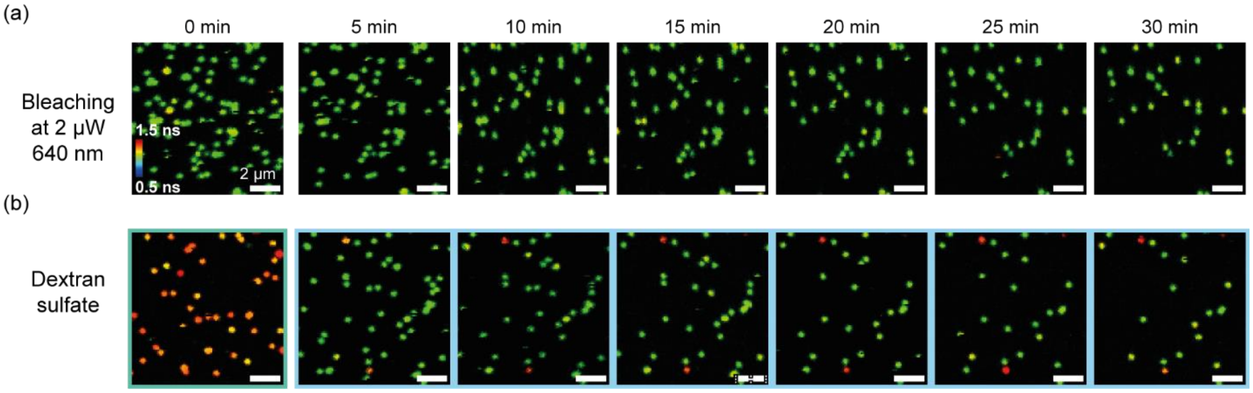

**Figure S8.** Real-time FLIM imaging of TLO with internal AF647 label at the DNA surface without photostabilization. a) Repetitive scanning of the same field of view will lead to gradual bleaching of AF647 labels on immobilized TLO nanostructures over time. b) Addition of dextran sulfate decomplexes the PLL-PEG coating within the first 5 min. Scale bar is 2 µm.

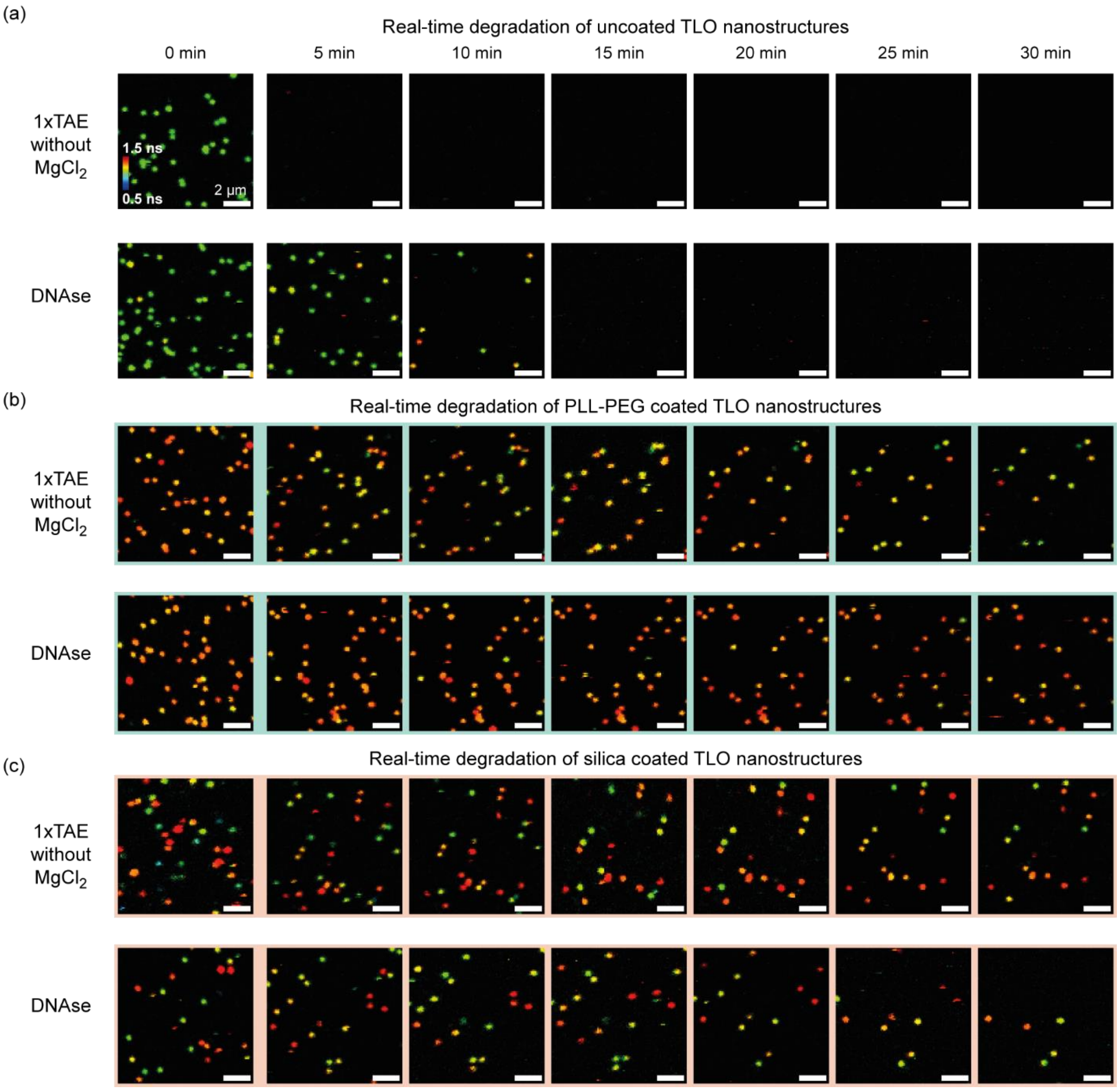

**Figure S9.** Real-time FLIM imaging of TLO with internal AF647 label at the DNA surface without photostabilization in degrading conditions. a) FLIM scans of uncoated TLO reveal a rapid degradation in magnesium-free conditions (1xTAE 0 mM $MgCl_2$) or under incubation in DNAse I. FLIM scans of PLL-PEG coated TLO (b) and silica coated TLO (c) over time reveal a highly improved stability in degrading conditions (1× TAE 0 mM $MgCl_2$, DNAse I). Scale bar is 2 µm.

**Table S10.** Fitted fluorescence lifetime populations for internal and external AF647 labels on TLO after degradation in magnesium ion free TAE buffer or in DNAse I solution. For every sample, the central fluorescence lifetime values and standard deviations of the Gaussian fit distributions are given after incubation over 30 min in the degrading condition.

| | | | | Fitted fluorescence lifetime populations (ns) | | | |
|---|---|---|---|---|---|---|---|
| **Sample** | **Coating** | **Degradation** | **Sample size** | $\tau_1$ | $\sigma_1$ | $\tau_2$ | $\sigma_2$ |
| Int. AF647 on TLO surface | PLL-PEG | 1xTAE w/o $Mg^{2+}$ | 756 | 1.12 | 0.06 | 1.67 | 0.17 |
| | | DNASe I | 1027 | 1.25 | 0.07 | 1.47 | 0.20 |
| | $SiO_2$ | 1xTAE w/o $Mg^{2+}$ | 664 | 1.06 | 0.05 | 1.55 | 0.21 |
| | | DNASe I | 886 | 1.29 | 0.15 | 1.58 | 0.24 |
| Int. AF647 at TLO interface | PLL-PEG | 1xTAE w/o $Mg^{2+}$ | 1059 | 1.64 | 0.13 | | |
| | | DNASe I | 1093 | 1.31 | 0.04 | 1.60 | 0.15 |
| | $SiO_2$ | 1xTAE w/o $Mg^{2+}$ | 1553 | 1.53 | 0.13 | 1.77 | 0.21 |
| | | DNASe I | 622 | 1.62 | 0.20 | | |
| Ext. AF647 at TLO surface center | PLL-PEG | 1xTAE w/o $Mg^{2+}$ | 908 | 1.53 | 0.12 | | |
| | | DNASe I | 1160 | 1.45 | 0.14 | | |
| | $SiO_2$ | 1xTAE w/o $Mg^{2+}$ | 1847 | 1.59 | 0.17 | | |
| | | DNASe I | 816 | 1.65 | 0.22 | | |

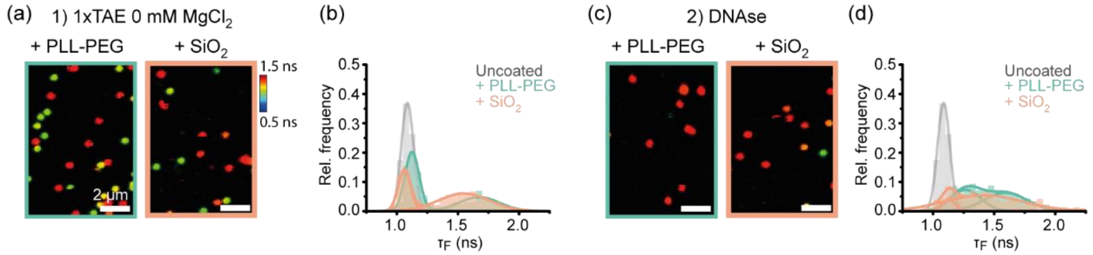

**Figure S10.** Two-step degradation study on coated TLO with internal AF647 label at the DNA surface. a) Exemplary FLIM scans of PLL-PEG and silica coated TLO after 30 min incubation in low-salt conditions (1xTAE without $MgCl_2$). b) Spot-integrated lifetime distributions reveal a partial degradation of the coating layers resulting in a subpopulation with lifetimes similar to an uncoated TLO. c) Exemplary FLIM scans of PLL-PEG and silica coated TLO after a second 30 min incubation in DNAse I solution. d) Spot-integrated lifetime distributions reveal a degradation mainly of the subpopulations with fluorescence lifetimes comparable to uncoated TLO. Scale bar is 2 µm.

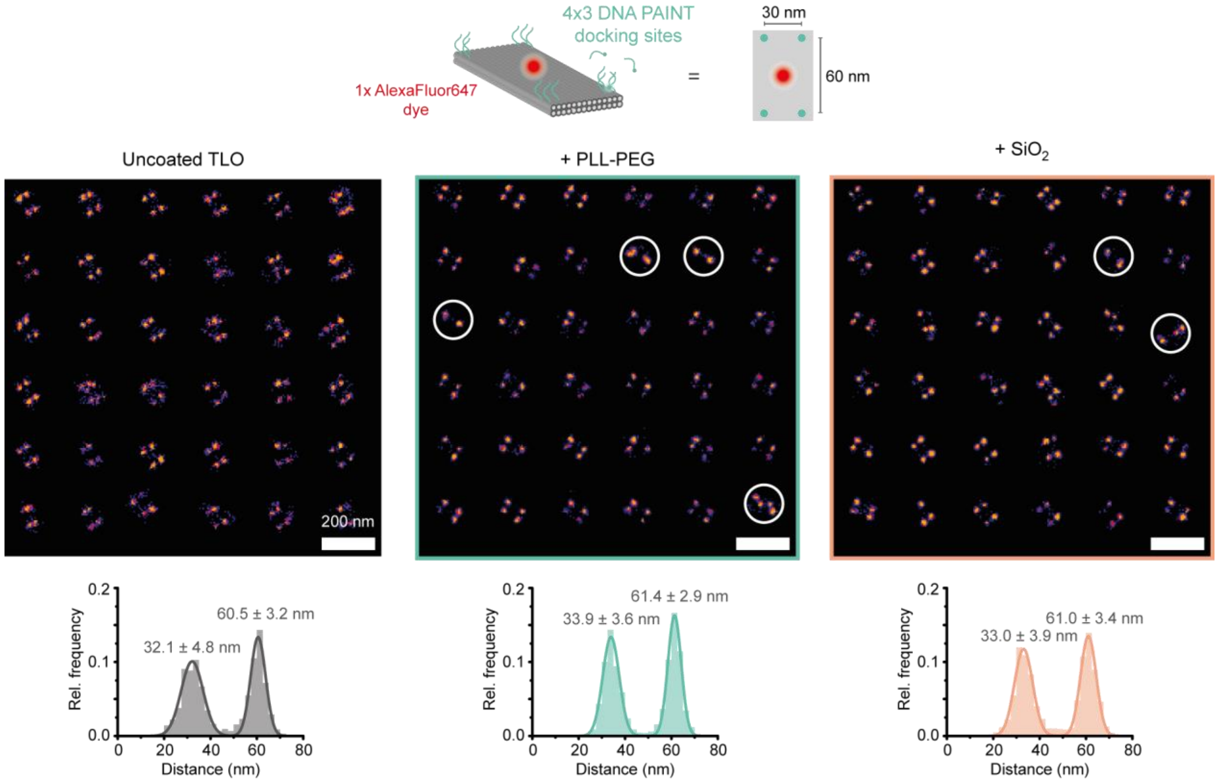

**Figure S11.** Distance analysis of 4x3 DNA PAINT TLO nanostructure. A nearest neighbour analysis revealed comparable short (ca. 30 nm) and long (ca. 60 nm) distances for uncoated TLO (grey), PLL-PEG coated TLO (green) and silica coated TLO (orange). Reconstructed DNA-PAINT images revealed a small fraction of deformed TLO nanostructures upon coating (highlighted with white circles). Scale bar is 200 nm.

**Table S11.** Fitted fluorescence lifetime populations for internal AF647 label on 4x3 DNA PAINT TLO. For every sample, the central fluorescence lifetime values and standard deviations of the Gaussian fit distributions are given before coating, after coating with PLL-PEG, and after coating with silica.

| | | Fitted fluorescence lifetime populations (ns) | | | |
|---|---|---|---|---|---|
| **Coating** | **Sample size** | $\tau_1$ | $\sigma_1$ | $\tau_2$ | $\sigma_2$ |
| uncoated | 837 | 1.06 | 0.04 | | |
| PLL-PEG | 704 | 1.28 | 0.05 | | |
| $SiO_2$ | 571 | 1.26 | 0.06 | 1.39 | 0.17 |

**Table S12.** Fitted dark-time populations for 4x3 DNA PAINT TLO. For every sample, the central dark-time values and standard deviations of the Gaussian fit distributions are given before coating, after coating with PLL-PEG, and after coating with silica.

| Coating | Sample size | $\tau_d$ (s) | $\sigma_d$ (s) |
|---|---|---|---|
| uncoated | 1003 | 6.9 | 2.0 |
| PLL-PEG | 1030 | 6.6 | 1.6 |
| $SiO_2$ | 723 | 6.0 | 1.8 |

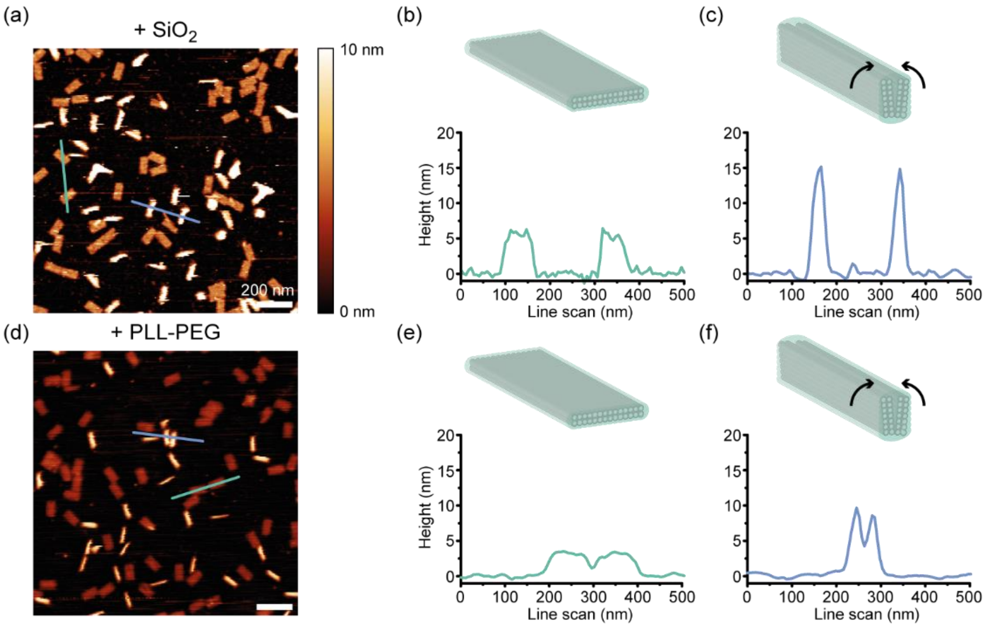

**Figure S12.** Exemplary AFM scans of partially rolling up TLO nanostructures. (a) AFM scan of silicified TLO with partially rolled-up nanostructures. b) Exemplary line scan (green line) of an intact silica-coated TLO nanostructure reveals heights of around 5 to 6 nm. c) Exemplary line scan (blue line) of a rolled-up, silica coated TLO reveals heights of up to 15 nm. d) AFM scan of PLL-PEG-coated TLO with partially rolled-up nanostructures. e) Exemplary line scan (blue line) of an intact PLL_PEG coated TLO nanostructure reveals heights of around 4 nm. f) Exemplary line scan (blue line) of a rolled-up, PLL-PEG coated TLO reveals heights of up to 10 nm. Scale bars represent 200 nm.

#### Supplementary Note 1: Mechanistic studies on the fluorescence lifetime shift: water quenching vs. restricted photoisomerization.

To better understand the mechanistic origin of the observed shift in the fluorescence lifetime upon coating, we went on to compare the observed fluorescence lifetime shifts of AF647 upon coating of the DNA origami with PLL-PEG or silica with expected fluorescence lifetimes where either quenching by water or by photoisomerization is eliminated. To determine the maximum contrast in the fluorescence lifetime induced by a complete suppressed quenching by water, we probed the fluorescence lifetime increase of an AF647 internally labeled to the upper surface of uncoated TLO origami when the $H_2O$ buffer is exchanged with a $D_2O$ analogue (Figure S13). We observed a shift from initially 1.08 ns to 1.51 ns, defining the expected range of fluorescence lifetime shift due to reduced water quenching.

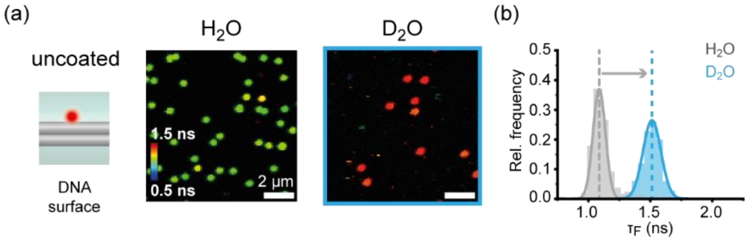

**Figure S13**. Fluorescence lifetime shift of AF647 label on TLO for suppressed water quenching. a) Exemplary FLIM scans of an internal AF647 label at the DNA surface of uncoated TLO, measured in a $H_2O$ buffer (left) and in a $D_2O$ buffer (right). b) Spot-integrated fluorescence lifetime distributions reveal a shift from 1.08 ns ($H_2O$) to 1.51 ns ($D_2O$) when quenching of the dye by water molecules is completely suppressed. Scale bars represent 2 µm.

Using fluorescence intensity autocorrelation analysis, fluctuations in the fluorescence signal of a fluorophore can be quantified by autocorrelating the single-molecule fluorescence trajectories. For AF647 diffusing in solution, a decay of the autocorrelation function due to photoisomerization in the range of µs has been observed. Here, an increase of steric restriction by, *e.g.,* increasing viscosity or by a chemical group in close proximity led to a decrease in the amplitude of the photoisomerization decay and a shift of the decay time to longer time scales.[12, 13] To investigate the change of the photoisomerization of AF647 upon coating of the DNA origami, we performed fluorescence intensity autocorrelation studies on immobilized uncoated and coated TLO, which were internally labeled with AF647 at the upper DNA surface (Figure S14). For uncoated nanostructures ($\tau_f$ ca. 1.05 ns), the sensor dye exhibited a tight distribution of the autocorrelation curves with a decay in the µs range attributed to photoisomerization as previously reported (Figure S14a).[12, 13]

For PLL-PEG-coated TLO, we performed autocorrelation analysis for the two obtained fluorescence lifetime populations around 1.33 and 1.57 ns individually (Figure S14b). While the autocorrelation for the shorter lifetime population revealed a tight distribution with an average decay curve quite similar to uncoated TLO, the higher lifetime population showed a quite heterogenous distribution of intensity autocorrelation curves with a significantly lower amplitude and slower decay time indicating a slowed-down photoisomerization. Additionally, a second decay process in the ms range occurred for the higher fluorescence lifetime population. We attributed this process to the formation of long-lived radical states of the AF647 dye, indicating a decreased accessibility of the dye for photostabilization additives in the imaging buffer.[9]

We performed the same intensity autocorrelation analysis on the two fluorescence lifetime populations of AF647 on silicified TLO around 1.29 and 1.61 ns (Figure S14c), however, for both fluorescence lifetime population heterogenous distribution of autocorrelation curves was obtained. In both cases, some curves exhibited a similar decay as uncoated TLO and others revealed a clearly lowered amplitude and slowed down decay time. The average autocorrelation of both populations revealed a lowered amplitude and slowed down decay time indicating a restricted photoisomerization similar to the average autocorrelation of the PLL-PEG coated sensor with a higher fluorescence lifetime around 1.57 ns.

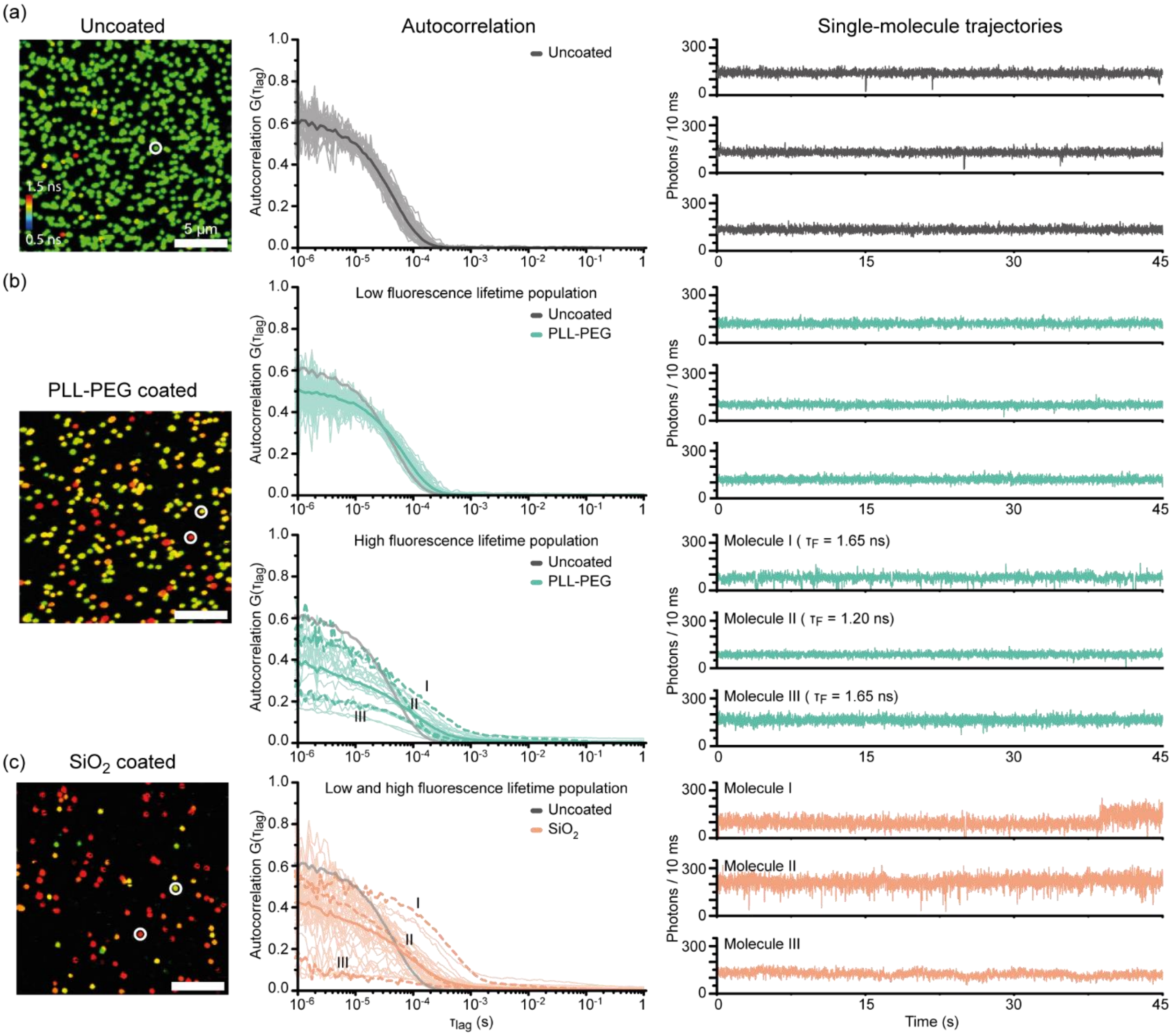

**Figure S14.** Fluorescence intensity autocorrelation on uncoated, PLL-PEG coated and silica coated TLO internally labeled with AF647 on the DNA surface. a) Exemplary FLIM scan of uncoated TLO (left), autocorrelation curves for picked nanostructures (dark grey curve represents average decay) (middle), and three exemplary single-molecule trajectories (right). b) Exemplary FLIM scan of PLL-PEG coated TLO (left). Autocorrelation curves for picked nanostructures with fluorescence lifetimes around 1.3 ns (top row) and around 1.6 ns (lower row) reveal different decay distributions and average decay curves (dark green curves) (middle), and three exemplary single-molecule trajectories for each population (right). c). Exemplary FLIM scan of silica coated TLO (left), autocorrelation curves for picked nanostructures (dark orange curve represents average decay) (middle), and three exemplary single-molecule trajectories (right). In some trajectories, a switching between two states was visible. Scale bars represent 5 µm.

The similar autocorrelation distributions for both silicified fluorescence lifetime populations could be explained by slow switching of the sensor dye between two environments with different grades of steric restriction, for example, by binding and unbinding to the DNA helix or coating agent. While we observed a slow switching in some of the single-molecule trajectories used for fluorescence intensity autocorrelation analysis (Figure S14c), we also noticed a more pronounced fluctuation in the fluorescence lifetime of silicified TLO when scanning the same field of view repeatedly over time than in PLL-PEG coated TLO (Figure S15). This could indicate a higher heterogeneity in the silica coating enabling a more dynamic switching behavior between different microenvironments. The fluorescence intensity autocorrelation analysis suggests, that the photoisomerization of the coated AF647 labels with a lower fluorescence lifetime around ca. 1.30 ns is only slightly affected and that the fluorescence lifetime increase upon coating stems dominantly from reduced water quenching. The higher fluorescence lifetime populations around 1.60 ns though seem to stem from additional restriction of the photoisomerization of the cyanine dye, possibly due to binding to DNA origami backbone or to the coating agent itself. Even though fluorescence intensity autocorrelation analysis revealed a quite heterogenous behavior AF647 upon coating and, in turn, complex photo-physics, the effects of reduced water quenching and restricted photoisomerization resulted in significant fluorescence lifetime shifts overall.

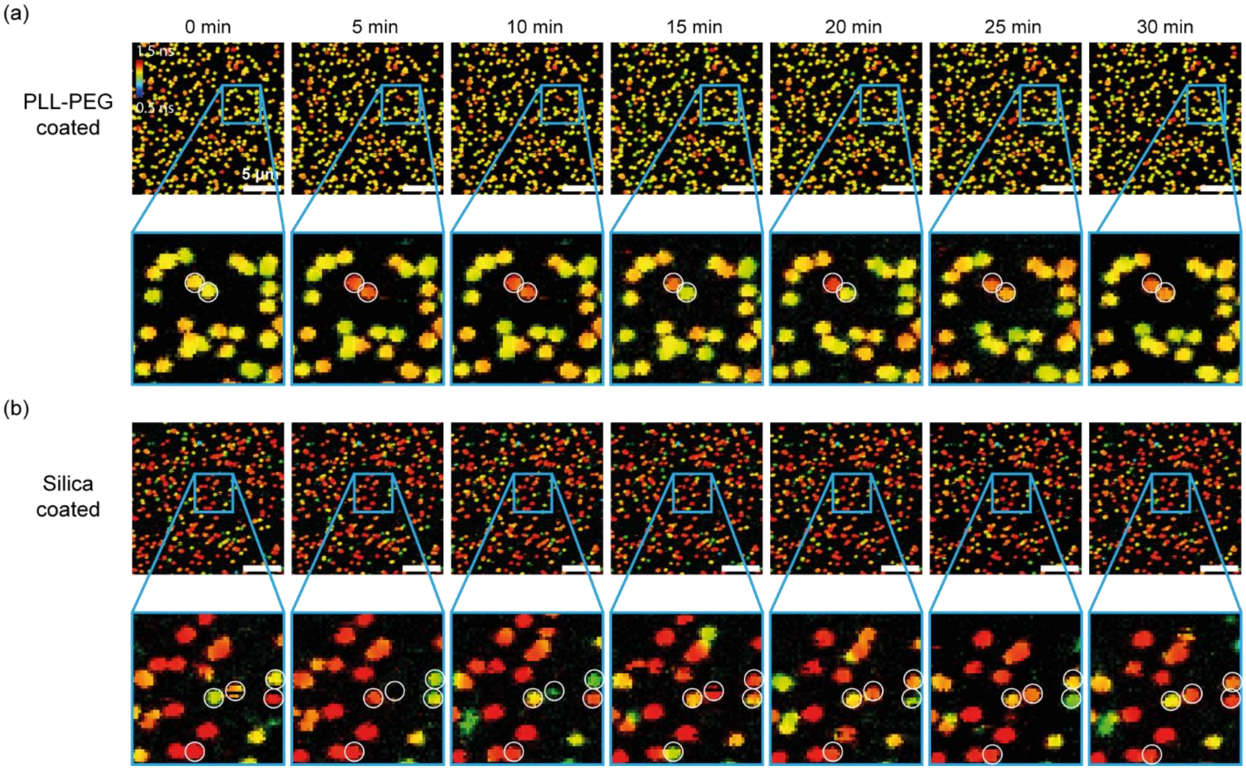

**Figure S15.** Slow FLIM fluctuations over time for PLL-PEG and silica-coated TLO with internal AF647 label on the DNA surface. a) Exemplary repetitive FLIM scans on PLL-PEG coated TLO exhibiting slight fluorescence lifetime fluctuations for a few nanostructures. b) Exemplary repetitive FLIM scans on silica coated TLO exhibiting pronounced fluorescence lifetime fluctuations for some nanostructures. Individual spots who exhibited pronounced fluctuations in the fluorescence lifetime are highlighted with white circles. Scale bars represent 5 µm.

### 3. Appendix

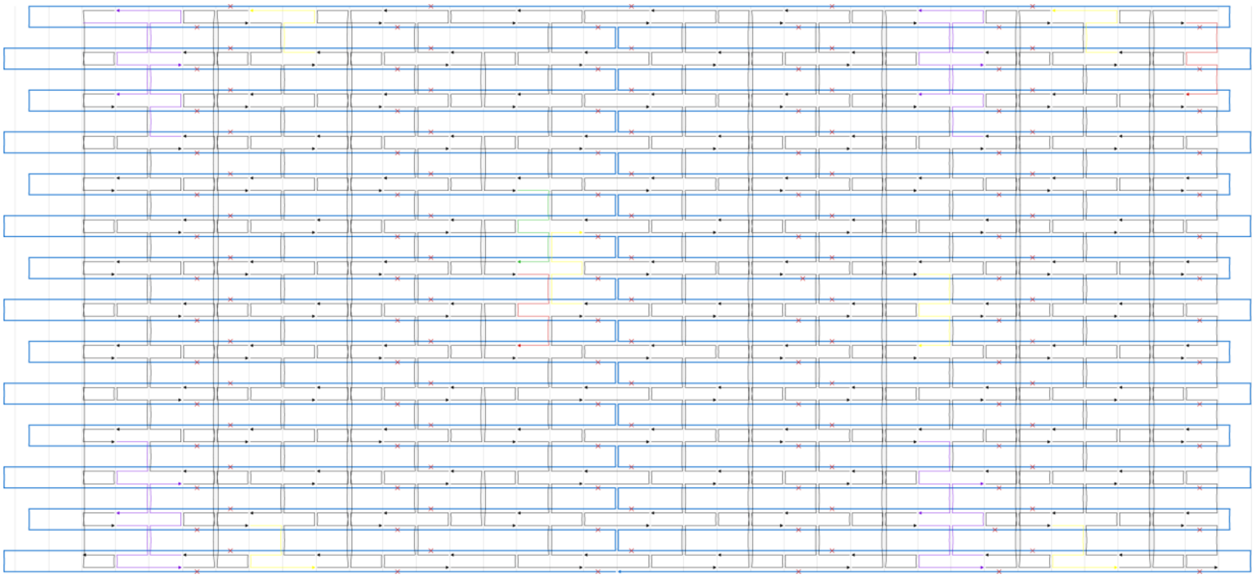

**Figure S16.** Cadnano design file of TLO design. In yellow biotinylated staples, in purple PAINT staples, in green Atto542 staple and in red AF647 staple (internally or externally labeled).

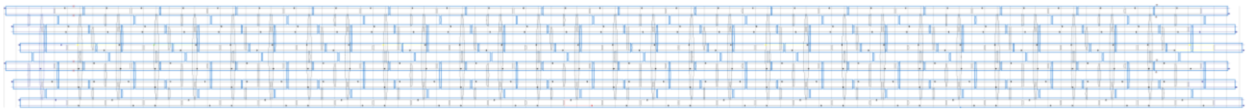

**Figure S17.** Cadnano design file of 12HB design. In yellow biotinylated staples, in green Cy3 labeled staple and in red AF647 staple (externally labeled).

**Table S13.** Staple strands of the TLO DNA origami. Sequences are denoted from 5'- to 3'-end. The numbers for the 5'- end 3'-end of the staples represent the helix number in the corresponding caDNAno file. Number in brackets represent the starting and ending position of the staple in the corresponding helix. Six biotin modified staple strands for surface immobilization are marked with biotin.

| Sequence (5` to 3`) | 5'-end ´ | 3'-end] |
|---|---|---|
| TTGACAAGCGAGAGGACCACATCCGGAAGC | 4[71] | 9[71] |
| TGCGGATGTAGCTCAATTAAGCAAGTACCAAA | 9[136] | 12[136] |
| CGAGAAAAGAATCATTAAGTAATTGAGAGAAT | 15[184] | 10[184] |
| CAGCAAAACGTTTGCCAACCACCAGAGTGTAC | 6[247] | 3[247] |
| AAACTCCAAGTTGATTCTACTAATTAAAAATT | 9[72] | 12[72] |
| TCGTCTTTGCTTTTGATTAGCGGGATAGCCCC | 0[231] | 5[231] |
| GCCGCCACTGGGAAGGTCTGCCAGAATTCGCG | 18[87] | 15[87] |
| GTACCGACAGCCAGTACGCAAGACGTAAATGC | 13[168] | 16[168] |
| TTTCATTCAGAGGCGTATAATCCATTATCA | 17[200] | 20[200] |
| TTGTGAATATTACAGATAGTAATGACCATA | 5[104] | 8[104] |
| AAACGCAATTGGCCTTGCGTCAGATCGAGAGG | 7[184] | 2[184] |
| AAACAGAAGATAGCTTCGTCGCTATGCGTTAT | 19[280] | 14[280] |
| TACCTGAGAACAAAATAACTATATAAAGAACG | 18[183] | 15[183] |
| CGTTTTCCGGAAACGCGGAATATAAGACTC | 5[200] | 8[200] |
| CGATAGCATGCCTTTAGATATTCAGGAAAGCG | 6[183] | 3[183] |
| GATGTGCTTTTCCCAGCATCGACACGGCCTTT | 17[136] | 20[136] |
| CCAATAGCCCTCATACACCATCCTGCGAAC | 15[56] | 10[56] |

| | | |
|---|---|---|
| CCTTGCTTACATCGGGAAATTATTCAATTCGA | 17[264] | 20[264] |
| CAATCAATTAAACACCGTATCATATTAATTAA | 12[295] | 17[295] |
| CTTCACCGCGTTGCGCGTAATCATGCGCGCCT | 26[87] | 23[87] |
| CAGTCTCTGATTTTGCGTTTAGTACCGCCACC | 3[184] | 1[199] |
| TTAAACAGTTCAGAAAGATAAGAGCTAACGGA | 8[135] | 6[120] |
| TCACCGACGGAACCAGTCAGAGCCAGTGCCTT | 6[279] | 3[279] |
| GACAGGAATGGTTGCTTTTTGGGGTCGAGG | 25[200] | 27[215] |
| AACGTGCTAGGAGGCCCCTACATTACATTGGC | 26[183] | 23[183] |
| TTTGCCAGGGCTGACCATTTCAACTTCCATTA | 7[88] | 2[88] |
| CTAAAGGAATTGCGAACTTTTGCGGGATCGTC | 0[135] | 1[135] |
| CAAAATAGAACCGGAACGAGTACTACGAAG | 7[56] | 2[56] |
| ATCAAGTTGCACCGTATGGCAACACGTAGAAA | 5[168] | 8[168] |
| AAGGTGGAAGCAAAGAACCAGATCAACTAA | 11[56] | 6[56] |
| CGACGGCTATTACGCTTGGTGTTCATCAAC | 18[119] | 15[119] |
| GAGAGGGCATGTCAACCAGCTTAGATGGGC | 13[104] | 16[104] |
| TCATCGAGTTCTGACCTGAGAATCATGGAAAC | 12[231] | 17[231] |
| GCACCAACATGACAACTCGGTTTATCAGCTTG | 2[55] | 0[40] |
| TTGTCACACATTGACATTTCGGTCGTTTTGCT | 7[216] | 2[216] |
| AAATACCCGGGTATTTATCAACTCCCAATC | 15[248] | 10[248] |
| CAGAGCACAGATATAAGCGCATGCTGAATA | 11[152] | 10[152] |
| TGGTGCTGACTGGTGTCGGTGCCCTCCGCTCA | 21[40] | 24[40] |
| GGCTTGAGTTAGGAATCTTTTGCAGACTATTA | 5[72] | 8[72] |
| GGGAGTTTTTTTCATCGACCTGACCAGGCG | 1[104] | 4[104] |
| ATGAATCGTGGGCGCCAAAGGGCGAAAAACCG | 25[136] | 27[151] |
| AAATCACCTTGAGCCAAAAAGGGCGTTACCAG | 5[264] | 8[264] |
| TCTCCGTAATCCAATATAAGAGAGTCTGGA | 15[152] | 14[152] |
| CAATCGTCACGCGTGGCGGGGAAGAGCGGG | 24[167] | 25[167] |
| TGATGCAGGGAACAAATTAAGTAAACAAAC | 16[167] | 17[167] |
| GGGACATTTAAAAGTTAATCAACAAGTTACAA | 23[216] | 18[216] |
| CCGGCGAATCACTTGCCGCAAATTCATCGCCA | 27[280] | 22[280] |
| TCAGACGAAGACACCATCACCAATAAGTCAGA | 4[199] | 9[199] |
| TACCGAGCAACAAGAGGCAACAGCTGATTGCC | 24[103] | 26[88] |
| AACTTAAAACCGTGCAGCGATCGGACCCCGGT | 19[88] | 14[88] |
| CTGAACCTGAAAAATCGATTATTTTTGACGCT | 21[168] | 24[168] |
| TACAACGTAATTGTAAACCATCGCCCACGC | 3[56] | 1[71] |
| CAATTCCAGGCAAAATTTTGCCCC | 24[39] | 26[32] |
| CAGAGCCCCAAAGACTTTGGGAAATAATAA | 4[263] | 9[263] |
| AGCTAAACTTCCTCGTGCCCACTACGTGAACC | 25[168] | 27[183] |
| TAAAGACAAAGGCCGTAATAATTTTTTCAC | 2[119] | 0[104] |
| AACATTATGACCCTGTTTTGAGAGAAATATGC | 12[135] | 10[120] |
| CAGTACCCCGCCACCTAGTAAATGAATTTT | 2[215] | 0[200] |
| ATAACCGAAAATACGTTATCATCGGAGTAATC | 1[72] | 4[72] |
| GTTGAGATATGGTTTATTCATCAACCTGATAA | 6[87] | 3[87] |
| TGATAATCGTTCTAGCCGCAAGGAAGTAGTAG | 14[87] | 11[87] |
| AGAATTAAAACAGGGAGAAGGCTTCAATAGCA | 9[168] | 12[168] |

| | | |
|---|---|---|
| CTGACCTTATTAAATTCTAAAAATAACGGA | 23[248] | 18[248] |
| TCCCGTATCATCATAGATGATGTGGGTAAC | 19[152] | 18[152] |
| AATCGCGTGAATTACTTTTTAATTTAGTTA | 18[215] | 15[215] |
| AGGGTAGCCAGCGAAAGAATAGAAAGGAACAA | 2[151] | 0[136] |
| CATATAACACAGGTCATTTACCCTAAAGAAGT | 10[87] | 7[87] |
| TTCGCCTGAATCAATAATTTATCAATGGTTTG | 18[247] | 15[247] |
| TTAATTGGTACGGTGATCATACTTGCGGGA | 9[104] | 12[104] |
| TCCCTTACGTCTGGTCGCCTCCGGTAGCTCTC | 22[55] | 19[55] |
| TTCGAGCTTTAGTTTGTGGGGCGCATGCAATG | 9[40] | 12[40] |
| GTCTCGTCCAGCGCATGCTCGTTAACTCAC | 20[71] | 25[71] |
| ATACCCAAGCGGGAGGTTTTGTTTCAGCTAAT | 8[231] | 13[231] |
| TGCCGTAACCGCCAGCCCAGAATCTATTAACA | 27[216] | 22[216] |
| ACCCTCAGAACGGCTAGGCGCAGACTTTGAAA | 1[136] | 4[136] |
| AATAGATACTGATAGGGCTATTTTGATTAG | 21[296] | 24[296] |
| AAGCGGTCGCCTAATGAATTGTTACCTGCATC | 26[55] | 23[55] |
| CAAGATTCCGAGGAATAAGCCCATTAGAGC | 11[248] | 6[248] |
| TCACCAGAAACCTGTTTGAGGAAGAATGCG | 26[119] | 23[119] |
| GAGCCCCCGGCCTTGGAGGCCAAGCAGAAG | 27[248] | 22[248] |
| AAAGTAAGCCTGAATCCTAATTTGCCCATCCT | 8[295] | 13[295] |
| AGCAAATTAAAGCTAGCCTGAGAATATAAA | 12[167] | 13[167] |
| CAAATAAGAATTGAGTACGCAATATATGGTTT | 10[247] | 7[247] |
| CAGGCAAACGAAACGTTGAAGGGACCAGAGCA | 17[40] | 20[40] |
| AGTCCTGATTACCAGTGATAAATAACGCTGAG | 13[264] | 16[264] |
| GACTGGATAGCGTCCAAAAACGAAGTCATTTT | 7[120] | 9[135] |
| GTTGATATCTCAGGAGTAAACAACTTTCAACA | 2[183] | 0[168] |
| TGGTAATTTAGCGTACATTTTCAGGGATAG | 3[248] | 1[263] |
| GCAACTGTGGGAACGGCAGAAACAGTTTTTTC | 17[72] | 20[72] |
| CTCAGAACAGGGAGGGAGGTGAATTAGCTATC | 4[295] | 9[295] |
| CTTATTAGTCACCAGTAAAATTCAATAACGGA | 5[232] | 8[232] |
| CATCGTAGCTTTTTCAGTAATTTATTTAACAA | 12[199] | 17[199] |
| TCTTCGCCAGTGCCACATTTGCGATGCTGA | 17[104] | 20[104] |
| GATGGCAAGGTTATATTAATTACAGGCAGAGG | 19[184] | 14[184] |
| ATTAATTTCTGGCCAAGCGGTCAGCTGAGAAG | 20[231] | 25[231] |
| AACTAAACTCCTTTTACGAGAAAATGTTTA | 10[119] | 7[119] |
| AGATTCACAACAAAGAAAACCCTCATTTCAAT | 23[184] | 18[184] |
| ACAACATTTACCTTATGTACAGCTCCATGT | 6[119] | 3[119] |
| GAGTAGATTCAAAGCGCGGATTGCATAAAAAC | 10[55] | 7[55] |
| TCTGGCCTTATTTCAATGATAAATTTTCATTC | 15[88] | 10[88] |
| AGACGATCGCTGGCAAGCAGCACACCGGAA | 23[56] | 18[56] |
| TGTTTTTAGCGCTTAAATCGGAACCCTAAAGG | 25[232] | 27[247] |
| GCCAGGGTGCAAGGCGACGGCGGACGTCGGAT | 18[151] | 15[151] |
| GCATCGTATTTCTGCTAGCTTTCAGGTGCCAT | 16[103] | 21[103] |
| AATGGGATAGGTCACGCAGCTGGCTAATCGTA | 16[135] | 14[120] |
| AGTGATGAAGGGTAAATACGGCTGTTGTAAAA | 20[135] | 18[120] |
| ATTAAATGTGAGCGAGATGAACGGGAAAGGGG | 15[120] | 17[135] |

| | | |
|---|---|---|
| ATAAATACATAAAGGATCAGTATTGGGAAG | 7[152] | 6[152] |
| GCACTCAAGCGGGGTCCGCACAGGTAAAAAAA | 22[151] | 19[151] |
| CATACATGCCAGACGTCTCAGAACCGCCACCC | 3[216] | 1[231] |
| GGTCTGAGTACTTCTGGAATACCAGTTGAAAG | 16[231] | 21[231] |
| CTTTCGAGGATTATACAAAGAGGCCTTGCCCT | 0[39] | 5[39] |
| ATTCAACCAGAAAAGCTCAAAAATTTTGAGGG | 13[72] | 16[72] |
| CATTAACAATCAGGTCGGATTAGAATTCATCA | 11[88] | 6[88] |
| ACAATCGGGCGCCATTGGCCTCAGTTTTTTAA | 18[55] | 15[55] |
| CAATATTAAGCACTAATGCGCCGCTACAGGGC | 24[231] | 26[216] |
| AATAAGAAAATCGGCTAATAATATCCAGTTAC | 15[280] | 10[280] |
| GTTGGGCGGTTGTGTATCACGACGGAGGTGTC | 19[120] | 21[135] |
| CACCCGCCTAATCAGTCTGGTAATGAACCCTT | 26[247] | 23[247] |
| GATTAAGATTTTCATTACCATTAGGAGAAAGG | 8[39] | 13[39] |
| GATAGTTGCGCCGACACTAAAACGCAAGCGCGACCCAAAT | 1[32] | 4[40] |
| TACCTTTTCTGTAAATAGATTAAGAGGCGTTA | 18[279] | 15[279] |
| TGGGTAAACGGCAGCACGCGGTCCGCGGATCA | 22[87] | 19[87] |
| TCTATCAGGGCGATGTAGAATCGAGGCGGT | 27[152] | 26[152] |
| CAAGCCCAGTATTAAGACGGGGTCACCACCCT | 1[264] | 4[264] |
| GACAATATTATTAGACATAGATTAAGTAACAG | 23[280] | 18[280] |
| GGGTAATAATAGCAGCGTTTTATTTATTTT | 9[200] | 12[200] |
| TTTTCCCCGTCAGATATTGCGTTTTGAGGA | 17[296] | 20[296] |
| ACGACGAGCCAACATAATATATCCTCCGGC | 13[200] | 16[200] |
| TGCAGATAACACCAGATATTCATTAAACAAAG | 6[55] | 3[55] |
| GTGCACTCCCGGCAAACCGTCGGTGAGGTGGA | 23[88] | 18[88] |
| AGTACATAATTGCTTTAATAATGGCGAACGTT | 17[232] | 20[232] |
| ATCACCCATGGAAATAGATTAAAGAGAGCCAG | 27[184] | 22[184] |
| CCGCCTGAGTTGGCATGAGTAACTGATTGT | 22[215] | 19[215] |
| GAGCAAGACAGCCATATTTTGCACTTATCATT | 9[264] | 12[264] |
| CAGCAAATCAAATATCAACCACCATATCAGAT | 22[183] | 19[183] |
| GTGAGCCTCCTCACAGCGTGCCAGGCGGTATG | 24[135] | 22[120] |
| GCGGGCCGTTTTCACGGCGCGGTTCTGCATTA | 23[120] | 25[135] |
| GACGAGAACATAACGCAGACGACGATCAAAAA | 5[40] | 8[40] |
| TCACCGTAAAGTATAGGCCAGAATCAAACAAA | 1[168] | 4[168] |
| CATTTTCGAAAAGGTAACCGCGCCATCCGGTA | 14[183] | 11[183] |
| CACTCCAGCTCCGTGGACAGCGCCCGTCAGCG | 16[39] | 21[39] |
| GAATTGAGGAGGTGAGCAGAGATAATCCAGAA | 21[232] | 24[232] |
| GGCTCATTACGTTAATATACTGCGCAAATGCT | 5[136] | 8[136] |
| AGTGTAGCGTCCATCACTGAGTAGGTGGCACA | 26[279] | 23[279] |
| AACATAAACTGAACACTTAGCAAATATAAAAG | 10[183] | 7[183] |
| CATAAGGTCATTAAACCCGGAATCGGAACG | 3[152] | 2[152] |
| TAACAACCAATAAACAGCCGTTGCGAACCT | 14[215] | 11[215] |
| GAGTAACAGCCTGTAGCCATGTACCGTAACAC | 3[280] | 1[295] |
| AATTTACCCTGTTTAGGAATCATCCTTGAA | 13[296] | 16[296] |
| ATACATATTCATTGACTTAATTTAGACGGG | 8[167] | 9[167] |
| ACAAATTCACAAGAAAGTCTTTCCCCAGCTAC | 14[279] | 11[279] |

| | | |
|---|---|---|
| CTGCCAGCTGAAATGTAAAGCAGCGCTTTC | 23[152] | 22[152] |
| AATCAAAATCCAATAATCTGGAAGTAATGCCG | 8[103] | 13[103] |
| CATCCTCAAGCGGTGCGTTCAGCAGTGTAAAG | 20[39] | 25[39] |
| TTCTAAGAGCAGTATGCCTGAACAGAAACCAT | 11[184] | 6[184] |
| AAATGAATGAGCGCTGCATGATAGTTTATT | 10[215] | 7[215] |
| GCCGCCAGATCAATAGAGCACCATAGAGATAA | 4[231] | 9[231] |
| AATGCCAAGGTTTCTTGTGTCACTGGTCATAG | 21[72] | 24[72] |
| ACGGAAAACAGTATCCGCCATTACAGGAAG | 19[56] | 14[56] |
| ACCGATTGCGCCACCCAGCCACCAATTCTGAA | 7[280] | 2[280] |
| CAGTTTGGTCGAATTCTCACTGCCGCTGGTAA | 27[88] | 22[88] |
| CTCAGAAAGGCGGATCGTTCCAGAGGCAGG | 1[200] | 4[200] |
| ATTGTATAAGTCAAATTATTTTAAGAGCTGAA | 14[55] | 11[55] |
| AAGAGAAGACCACCCTACGATCTAAAGTTTTG | 2[247] | 0[232] |
| TTTAGAAGAACGCCACCCAAAACAGGCTGC | 12[71] | 17[71] |
| TTGCCGTTTGTGGTGCGCCCTGCGCGCTTTCC | 20[103] | 25[103] |
| TTAGGTTGTTCATCAAAATTATTCAATCAATA | 16[199] | 21[199] |
| ATAAAACAGAAGGTTACCTTTGCCAAGGGTTA | 22[247] | 19[247] |
| AATTTTATCAGATAGCAATAGCAATATCACCG | 11[280] | 6[280] |
| CCTGAGTATTGTTAAATTTAAATTCCGGAAAC | 12[39] | 17[39] |
| TACTTAGCCGGAACGACAGAGGCTTAAGAACT | 3[120] | 5[135] |
| CATAGGCTAGGGGGTAGTAGAAAGGAGTACCT | 4[103] | 9[103] |
| GTTGAAAAGAAATCCGGAGGAAGTTTTAATCA | 0[103] | 5[103] |
| AAGGAAAAGTTGCTATTATTTAAATAGATA | 8[263] | 13[263] |
| TCAGAGCCGATTAGGATGATACAGCCAGAGCC | 1[232] | 4[232] |
| AAGAGTCCCATATCAAGAAACATATCTTTA | 16[263] | 21[263] |
| CCTGGGGTCACGCTGGCCCTTATAAATCAAAA | 25[40] | 27[55] |
| CAACTCGGAAAGCGTAACCACCCCGAGTAA | 20[263] | 25[263] |
| AACATAGCATAAAGAAGAATATACGAGCCGTC | 16[295] | 21[295] |
| CTCATAGAAGTTTTAAGGCTGACATAATCA | 0[263] | 5[263] |
| TTAAAAATAACAACTATTTACAAATGCACGTA | 22[279] | 19[279] |
| GACGACGAAGAGACGATAACCTACCGCAAG | 16[71] | 21[71] |
| ACCAGCGGCCACCAGATCTTTTGACTCCTC | 7[248] | 2[248] |
| CGTGGACTCCAACGTCAGGGTGGTTTTTCTTT | 27[120] | 26[120] |
| TAAATCCGAACCGAACAGGACGGCGACAGA | 4[167] | 5[167] |
| CAAAGAATTAGCAAAACATGTTTTATCTACAA | 11[120] | 13[135] |
| CCAAGAAGACCGTGTATAAAGCGTGAATAA | 12[263] | 17[263] |
| TAATGCTGGCTTAGAGATCCCCCTGAATCGTC | 10[151] | 7[151] |
| TAGTCAGCATCAATTCCCAATTAATATGAT | 8[71] | 13[71] |
| CAGTACAAACTACAACGTGCCCGTAACCTATTCCGGAACC | 0[303] | 5[295] |
| AAAATAAAAACAATGACGAACAAAGACATTCA | 10[279] | 7[279] |
| GTTTCAGCGGAGTGAGACAGCATAGGTGTA | 0[167] | 1[167] |
| AGTCGGGTGAGACGGTCCACTATTAAAGAA | 25[104] | 27[119] |
| GAATAGCCTGTGTGAAGTGAGCCATAAACA | 27[56] | 22[56] |
| GGAGCACTACCGAACGAAGAATACAAGAACTC | 21[264] | 24[264] |
| AAACTAGTAGCTATTAATACTTAGGCAAGG | 14[119] | 11[119] |

| | | |
|---|---|---|
| TCTGGTCCAACAGTGCGACCAGACAGGAAA | 21[200] | 24[200] |
| GGAATTAAAAAAAGCATTGCAGTCACCTTG | 20[167] | 21[167] |
| TTTAGAAGTTTTGAATCCCTAAAAAACCGTTG | 20[295] | 25[295] |
| ATTGTGTCTCTCCAAAGTCGCTGAGGCTTGCA | 3[88] | 1[103] |
| TAGCAATGGAGCGGGGAAAGGA | 25[296] | 27[303] |
| CCGGAGACAGCAAATATCAGCTCAGAAGATCG | 13[40] | 16[40] |
| AACAGTAGCGCCTGTTAAACCAAGCCTTAAAT | 14[247] | 11[247] |
| ATTTCATCAACAAGCAAACATGTTAACGTCAA | 15[216] | 10[216] |
| TTTTGCGGCAGTCACACCACGCTGGGATTTTA | 20[199] | 25[199] |
| GAAGCCTTTCCTGTAGTCATATGTTGCGGGCC | 12[103] | 17[103] |
| AAACTATCGATTTAGTGCGCGTAACCACCA | 24[263] | 26[248] |
| AACGCTCAAATCAAGTTTTGACGAGCACGTAT | 24[199] | 26[184] |
| TTGCGTATGCCAACGCCCTGTTCTGGGGGTTT | 26[151] | 23[151] |
| ATCAAGAACAAAAGAATTCCTGATGAAGGAGC | 17[168] | 20[168] |
| CTTATTACACGCGAGGCCTTTACACTGTCCAG | 8[199] | 13[199] |
| GCAAGGCATCGGCATGGAGGTTGTAAGCGT | 6[215] | 3[215] |
| AGCCGGGAACCAGCTGTTAAACCGCCAGCA | 22[119] | 19[119] |
| TTGGATTAAGACTACCCTTTTTTAGCCATATT | 19[216] | 14[216] |
| CAACGTAACGTTTACCCAAAAGGACGTTTTAA | 4[39] | 9[39] |
| TAATAACACGTGGCGACGCTAGGGCGCTGGCA | 24[295] | 26[280] |
| GCGTACTACGGTACGCATTGCATAATAAAA | 26[215] | 23[215] |
| GAGCCTTGAGATTTGAATGCCAGTAAATTG | 0[71] | 5[71] |
| CCCGACTTAAGAACTGAATATCAGTACCATTA | 11[216] | 6[216] |
| CCCACGCTCACTGTTTGCGGCCTCCCCGGG | 21[104] | 24[104] |
| CCCACAAGAAACGATTTTTTGAAGTACCGCAC | 9[232] | 12[232] |
| GAGGACAGATGAACGGTGCGATTTTTGAGGAC | 4[135] | 2[120] |
| CAGCATCATCCGCCGGGTCATACCTCGCGTCC | 21[136] | 24[136] |
| AAAAATCTATACCAGTCTGACCAACGGTCAAT | 6[151] | 3[151] |
| CTGTATGGGAATTTACAAGTGCCGCTGTAGCG | 0[199] | 5[199] |
| CTGTTTCCCGAGATATGAGAGAGTTGCAGC | 24[71] | 26[56] |
| AGGCTATCGAGAATCGTAACAACCTTGACCGT | 13[136] | 16[136] |
| GAACCTAAATAGTGATATGTGACAACGCTC | 19[248] | 14[248] |
| TGAGTTTTATTTCGGATAAACACCGCCACC | 1[296] | 4[296] |
| TTACCGATCCAGAGCTTACCAAGTAGAAAC | 9[296] | 12[296] |
| GCCTCCCTTCATTAAAAGGTAATTTTAAGA | 5[296] | 8[296] |
| GCAGAACGGGCTTAATTAAATTTAAAATCATA-**Biotin** | 13[232] | 16[232] |
| AACGGGTATATATTCGAAAAAGGCTCCAAAAG-**Biotin** | 2[87] | 0[72] |
| ATTAATTGCCTGGCCCGGGTTGAGTGTTGTTC-**Biotin** | 25[72] | 27[87] |
| ACATGAAAATAGGAACCATTCCACAGACAGCC-**Biotin** | 2[279] | 0[264] |
| GCAAACAAAGGTCATTAATCGGTTTAAAGCCT-**Biotin** | 14[151] | 11[151] |
| AAGAGTCTGGTCACGCAGCTTGACGGGGAAAG-**Biotin** | 25[264] | 27[279] |

**Table S14.** Staple strands of the 12HB DNA origami. Sequences are denoted from 5'- to 3'-end. The numbers for the 5'- end 3'-end of the staples represent the helix number in the corresponding caDNAno file. Number in brackets represent the starting and ending position of the staple in the corresponding helix. Six biotin modified staple strands for surface immobilization are marked with biotin.

| Sequence (5` to 3`) | 5'-end ´ | 3'-end] |
|---|---|---|
| AAAGGGCGCTGGCAAGTATTGGC | 11[681] | 10[668] |
| GCGCCTGAATGCCAACGGCCCAGCCTCCCGCGTGCCTGTTCTTCTTTTT | 7[42] | 8[25] |
| TTGACGGGGAAAGCTTCACCAGAAATGGCATCACT | 11[651] | 6[658] |
| CATTCAACCCAAAATGTAGAACCCTCATGAATTAGTACAACC | 9[147] | 5[160] |
| TCAGAGGTGTGTCGGCCAGAATGAGTGCACTCTGTGGT | 4[60] | 7[62] |
| GGCATAAGCGTCTTCGAGGAAACGCA | 8[466] | 9[482] |
| TACATAAATTCTGGGCACTAACAACT | 8[634] | 9[650] |
| CAATCCAAAATACTGAACAGTAG | 3[457] | 10[458] |
| CATAGTTAATTTGTAAATGTCGC | 3[541] | 10[542] |
| GAACAAGAGTCCACCAATTTTTTAGTTGTCGTAGG | 11[483] | 6[490] |
| TTGAAGCCCTTTTTAAGAAAAGT | 7[441] | 7[463] |
| AAGCACAGAGCCTAATTATTGTTAGCGATTAAGACTCCTT | 7[464] | 8[448] |
| GATGTTTTTCTTTTCACCA | 10[289] | 11[302] |
| GGTCACGCCAGCACAGGAGTTAG | 3[373] | 10[374] |
| TGAACAGCTTGATACCGATAGTT | 8[363] | 8[341] |
| AAAATTCCATTCAGGCTTTTGCAAAA | 8[256] | 9[272] |
| TCCCATCCTAATGAGAATAACAT | 0[496] | 0[474] |
| ATCAGCGGGGTCAGCTTTCAGAG | 3[56] | 3[78] |
| TTCGCTATTCGCAAGACAAAGTTAATTTCATCTTC | 5[539] | 4[546] |
| TTGAGAATATCTTTCCTTATCACTCATCGAGAACA | 5[497] | 4[504] |
| GGGCGTGAAATATTAGCGCCATTCGC | 8[130] | 9[146] |
| GGCGCCCCGCCGAATCCTGAGAAGTGAGGCCGATTAAAGG | 3[667] | 0[665] |
| TTTTTTGTTTAATAAAGTAATTC | 3[476] | 3[498] |
| AAATCAGCCAGTAATAACACTATTTTTGAAGCCTTAAATC | 7[506] | 8[490] |
| AGCACTAAATCGGATCGTATTTAGACTTATATCTG | 11[609] | 6[616] |
| GGTGCCGTCGAGAGGGTTGATAT | 8[405] | 8[383] |
| GTCAGAATCAGGCAGGATTCGCG | 3[205] | 10[206] |
| TTTTTTATAACGTGCTTTCCTCTTTATAACAGTACTAT | 2[698] | 3[678] |
| AGACGGGAGAATTGACGGAAATT | 0[454] | 0[432] |
| TAAGCCAGAGAGCCAGAAGGAAACTCGATAGCCGAACAAA | 4[480] | 7[482] |
| CGCCTGACGGTAGAAAGATTCTAATGCAGATACAT | 5[245] | 4[252] |
| CAGTCTTGATTTTAAGAACTCAACGTTGCGTAT | 0[263] | 11[272] |
| CATAGAATTTGCGGTTTGAAAGAGGA | 8[298] | 9[314] |
| GCGCAGCGACCAGCGATTATATATCATCGCCTGAT | 5[287] | 4[294] |
| TTTTTAAAAACGCTCATGGAAATA | 8[698] | 8[679] |
| AATCAGTTAAAACGTGGGAGAAA | 3[121] | 10[122] |
| AGACAACCTGAACAGTATTCGAC | 3[625] | 10[626] |
| TTTGCAACCAGCTTACGGCGGTGGTGAGGTTTCAGTTGAGGATCCTTTTT | 3[25] | 10[29] |
| TGCAACACTATCATAACCCTCGT | 7[231] | 7[253] |
| AACGAACCTCCCGACTTGCGGGA | 8[531] | 8[509] |

| | | |
|---|---|---|
| CCGAACGGTGTACAGACCAGGCG | 8[321] | 8[299] |
| ATTCAAGGGGAAGGTAAATGTGGCAAATAAATC | 0[431] | 11[440] |
| GTCACCAGTACAAGGTTGAGGCA | 3[350] | 3[372] |
| TAAATCGGTTGGTGCACATCAAAAATAA | 6[153] | 2[140] |
| AGACGGCGAACGTGGCGAG | 10[667] | 11[680] |
| CCCTTCATATAAAAGAACGTAGAGCCTTAAAGGTGAATTA | 11[429] | 0[413] |
| AACTTTAATCATGGGTAGCAACG | 3[266] | 3[288] |
| ACCATCACCCAAATAAACAGTTCATTTGATTCGCC | 11[567] | 6[574] |
| TGCCTAATGAGTGAGAAAAGCTCATATGTAGCTGA | 11[147] | 6[154] |
| TTTTTTGGTAATGGGTAACCATCCCACTTTTT | 1[21] | 2[25] |
| GGAGCAGCCACCACCCTTCGCATAACGACAATGACAACAA | 7[338] | 8[322] |
| AAAAGTGTCAGCAACAATTGCAGGCGCT | 6[69] | 2[56] |
| GGTTTGCGCATTTTAACGCGAGGCGT | 8[508] | 9[524] |
| AAAAGAATAGCCCGATACATACGCAGTAAGCTATC | 11[441] | 6[448] |
| TTTCACGAGAATGACCATTTTCATTTGGTCAATAACCTGT | 7[212] | 8[196] |
| TCGGTCATACCGGGGGTTTCTGC | 8[69] | 8[47] |
| CCTCCGAAATCGGCAAAAT | 10[415] | 11[428] |
| TTCCATTGACCCAAAGAGGCTTTGAGGA | 2[307] | 3[307] |
| ACGCGTCGGCTGTAAGACGACGACAATA | 2[517] | 3[517] |
| GTCCGTCCTGCAAGATCGTCGGATTCTCTTCGCATTGGACGA | 9[105] | 5[118] |
| GTCAGTCGTTTAACGAGATGGCAATTCA | 6[615] | 2[602] |
| GAGCTTAAGAGGTCCCAATTCTGCAATTCCATATAACAGT | 4[228] | 7[230] |
| GCAGCACTTTGCTCTGAGCCGGGTCACTGTTGCCCTGCGGCTTTTT | 10[48] | 0[21] |
| TACCTGGTTTGCCCCAGCA | 10[373] | 11[386] |
| AATGCTGTAGCTGAGAAAGGCCG | 4[209] | 4[187] |
| CTATATTAAAGAACGTGGA | 10[499] | 11[512] |
| CGGTAGTACTCAATCCGCTGCTGGTCATGGTC | 0[53] | 11[62] |
| CTTGAAAACACCCTAACGGCATA | 3[247] | 10[248] |
| AAGTAAGAGCCGCCAGTACCAGGCGG | 8[382] | 9[398] |
| AAAAGATAGGGTTGAGTGT | 10[457] | 11[470] |
| TTCGCCATAAACTCTGGAGGTGTCCAGC | 2[55] | 3[55] |
| AGGGCGAAAAACCGATTTAACGTAGGGCAAATACC | 11[525] | 6[532] |
| CCCACATGTGAGTGAATAACTGATGCTTTTAACCTCCGGC | 11[555] | 0[539] |
| TTTTTAGGAGCGGGCGCTAGGAAGGGAAGAAAGCGAATTTTT | 10[702] | 11[702] |
| TGCCATACATAAAGATTAACTGAACACCAACAGCCGGAATAG | 9[441] | 5[454] |
| TTTTTCCGGTGCAGCACCGATCCCTTACACTTGCC | 5[29] | 4[52] |
| ACAGCTGATTGCCCGTCGCTGCGCCCACACGTTGA | 11[315] | 6[322] |
| ATTAAAATAAGTGCGACGATTGGCCTTG | 2[391] | 3[391] |
| AAAACGAAAGAGGCTCATTATAC | 0[286] | 0[264] |
| TGTCCAAGTACCAGAAACCCCAG | 3[499] | 10[500] |
| TTACCAATAAGGCTTGCAGTGCGGAAGTTTAGACTGGATA | 7[254] | 8[238] |
| TTAGTGTGAATCCCTCTAATAAAACGAAAGAACGATGAATTA | 9[231] | 5[244] |
| ATCAGAGCCTTTAACGGGGTCTTAATGCCCCCTGC | 5[371] | 4[378] |
| TTACCTCTTAGCAAATTTCAACCGATTG | 6[447] | 2[434] |

| | | |
|---|---|---|
| AAAACGGAATACCCAAAAGAACT | 8[489] | 8[467] |
| GTCCACGCGCCACCTCACCGTTGAAACA | 11[364] | 6[364] |
| TTTTTATCCAGCGCAGTGTCACTGC | 7[21] | 7[41] |
| GATGAATAAATCCTGTAGGTGAGGCGGTAGCGTAAGTCCTCA | 9[609] | 5[622] |
| GCTAAATCGGTTTGACTATTATA | 3[182] | 3[204] |
| CAGCTTTGAATACCAAGTTACAA | 7[567] | 7[589] |
| GGTTGCTTTGACGAGCACGTTTTT | 3[679] | 3[698] |
| CATGCCAGTGAGCGCTAATATCCAATAATAAGAGC | 5[455] | 4[462] |
| TATGCATTACAGAGGATGGTTTAATTTC | 2[265] | 3[265] |
| ACTGCCCGCTTTCCTGAAAAGCTATATTTTAAATA | 11[189] | 6[196] |
| TGATTTAGAAAACTCAAGAGTCAATAGT | 6[573] | 2[560] |
| TGGGCGCCAGGGTGATTCATTAGAGTAACCTGCTC | 11[273] | 6[280] |
| TGCAACTCAAAAGGCCGTACCAAAAACA | 6[195] | 2[182] |
| AAATAGGTAATTTACAAATAAGAAACGA | 2[475] | 3[475] |
| TGTTCCAACGCTAACGAACAAGTCAGCAGGGAAGCGCATT | 11[471] | 0[455] |
| GTGCCTGCTTTAAACAGGGAGAGAGTTTCAAAGCGAACCA | 11[219] | 0[203] |
| GTTTGATGGTGGTTCAGAACCCCGCCTCACAGAAT | 11[399] | 6[406] |
| TCACCGTCACCGGCGCAGTCTCT | 0[412] | 0[390] |
| AGACGTCGTCACCCTCAGATCTTGACGCTGGCTGACCTTC | 7[296] | 8[280] |
| TTTAGCAAACGCCACAATATAACTATATTCCCTTATAAATGG | 9[525] | 5[538] |
| AGCGTATCATTCCACAGACCCGCCACAGTTGCAGCAAGCG | 0[347] | 11[363] |
| GTATGTGAAATTGTTATCC | 10[79] | 11[92] |
| CCGAACTTTAATAAAAGCAAAGCGGATT | 2[223] | 3[223] |
| GTGAGTTAAAGGCCGCTGACACTCATGAAGGCACCAACCT | 11[303] | 0[287] |
| GCGCCCGCACCCTCTCGAGGTGAATT | 8[340] | 9[356] |
| ACAGTTTTTCAGATTTCAATTACCGTCGCAGAGGCGAATT | 4[606] | 7[608] |
| TTTAGAACGCGAATTACTAGAAAACTATAAACACCGGAAT | 4[564] | 7[566] |
| TGACCTAAATTTTTAAACCAAGT | 4[545] | 4[523] |
| TAAAGAGGCAAAATATTTTATAA | 3[163] | 10[164] |
| GTTTACCGCGCCCAATAGCAAGC | 7[483] | 7[505] |
| TACCGGGATAGCAATGAATATAT | 3[331] | 10[332] |
| AAATTGTGTCGAGAATACCACAT | 4[293] | 4[271] |
| AAATGCGTTATACAAATTCTTAC | 8[573] | 8[551] |
| CAGATATAGGCTTGAACAGACGTTAGTAAAGCCCAAAAATTT | 9[315] | 5[328] |
| TAAGATCTGTAAATCGTTGTTAATTGTAAAGCCAACGCTC | 7[548] | 8[532] |
| CATTCTATCAGGGCGATGG | 10[541] | 11[554] |
| CTCCAATTTAGGCAGAGACAATCAATCAAGAAAAATAATA | 11[513] | 0[497] |
| GAGACAAAGATTATCAGGTCATTGACGAGAGATCTACAAA | 4[186] | 7[188] |
| AGGGACAAAATCTTCCAGCGCCAAAGAC | 2[433] | 3[433] |
| AAAATTTTTTAAAATGAGCAAAAGAA | 8[592] | 9[608] |
| CATCGGGAGAAATTCAAATATAT | 4[587] | 4[565] |
| ATCATTTACATAAAAGTATCAAAATTATAAGAAACTTCAATA | 9[567] | 5[580] |
| GCTACGACAGCAACTAAAAACCG | 3[289] | 10[290] |
| TTAGGTTGGGTTATAGATAAGTC | 0[538] | 0[516] |

| | | |
|---|---|---|
| TATTGCCTTTAGCGTCAGACTGT | 7[399] | 7[421] |
| TTTTTCCGGGTACCGAGCTCGAATTCGTAATCTGGTCA | 11[29] | 10[49] |
| CTAAAGACTTTTAGGAACCCATG | 3[308] | 3[330] |
| GTGGAACGACGGGCTCTCAACTT | 3[79] | 10[80] |
| TCAGGTGAAATTTCTACGGAAACAATCG | 6[111] | 2[98] |
| AAGACGCTGAGACCAGAAGGAGC | 3[560] | 3[582] |
| AGCAGTCGGGAAACCTGTC | 10[205] | 11[218] |
| AACAACATGTTCATCCTTGAAAA | 3[518] | 3[540] |
| ATAATGAATCCTGAGATTACGAGCATGTGACAAAAACTTATT | 9[483] | 5[496] |
| GAGGTAACGTTATTAATTTTAAAACAAATAATGGAAGGGT | 11[597] | 0[581] |
| ACCGCATTCCAACGGTATTCTAAGCGAGATATAGAAGGCT | 4[522] | 7[524] |
| CAGCATCAACCGCACGGCGGGCCGTT | 8[46] | 9[62] |
| GCTCAAGTTGGGTAACGGGCGGAAAAATTTGTGAGAGATA | 11[93] | 0[77] |
| GGAATCGGAACATTGCACGTTAA | 3[583] | 10[584] |
| ATAAGAAGCCACCCAAACTTGAGCCATTATCAATACATCAGT | 9[399] | 5[412] |
| GGCGACACCACCCTCAGGTTGTACTGTACCGTTCCAGTAA | 11[387] | 0[371] |
| CATGTCAGAGATTTGATGTGAATTACCT | 6[279] | 2[266] |
| AATAGCTGTCACACGCAACGGTACGCCAGCGCTTAATGTAGTA | 9[651] | 5[664] |
| GCAGCACCGTAAGTGCCCGTATA | 4[419] | 4[397] |
| ATGAATCCCAGTCACGATCGAACGTGCCGGCCAGAGCACA | 7[86] | 8[70] |
| TATGTGATAAATAAGGCGTTAAA | 7[525] | 7[547] |
| TTAATGAATCGGCCATTCATTCCAATACGCATAGT | 11[231] | 6[238] |
| ATTCTTTTCATAATCAAAATCAC | 8[447] | 8[425] |
| AATCGTTGAGTAACATTGGAATTACCTAATTACATTTAAC | 7[590] | 8[574] |
| ATTTTGCCAGAGGGGGTAATAGT | 8[279] | 8[257] |
| AGCGCCACCACGGAATACGCCTCAGACCAGAGCCACCACC | 7[422] | 8[406] |
| AAAAAAGGCAGCCTTTACAATCTTACCAGTTTG | 0[473] | 11[482] |
| TAATCGTAGCATTACCTGAGAGTCTG | 8[172] | 9[188] |
| CAAGTGCTGAGTAAGAAAATAAATCCTC | 6[405] | 2[392] |
| GGCTAAAGTACGGTGTCTGGAAG | 7[189] | 7[211] |
| CCTACATACGTAGCGGCCAGCCATTGCAACAGGTTTTT | 8[678] | 9[698] |
| CTATTTCGGAACGAGTGAGAATA | 4[377] | 4[355] |
| TCAACATCAGTTAAATAGCGAGAGTGAGACGACGATAAAA | 4[270] | 7[272] |
| AATAACGCGCGGGGAGAGG | 10[247] | 11[260] |
| AAGAGATTCATTTTGTTTAAGAGGAAGC | 6[237] | 2[224] |
| CAAATGGTTCAGAAGAACGAGTAGAT | 8[214] | 9[230] |
| AAAAGGGCGACAATTATTTATCC | 3[434] | 3[456] |
| ATAGCTGTTTCCTGGAACGTCCATAACGCCGTAAA | 11[63] | 6[70] |
| TGTAGGGGATTTAGTAACACTGAGTTTC | 2[349] | 3[349] |
| AAAAATCTACGTGCGTTTTAATT | 0[244] | 0[222] |
| AGAGTTTATACCAGTAGCACCTGAAACCATCGATA | 5[413] | 4[420] |
| GTGTATTAAGAGGCTGAGACTCC | 7[357] | 7[379] |
| GAAGTCAACCCAAATGGCAAAAGAATACTCGGAACAGAATCC | 9[273] | 5[286] |
| CGGTTAACAAAGCTGCTGTAACAACAAGGACGTTGGGAAG | 11[261] | 0[245] |

| | | |
|---|---|---|
| ACTACCTTTAAACGGGTAACAGGGAGACGGGCA | 0[305] | 11[314] |
| AATCCAAAAAAAAGGCTCCAAAA | 7[315] | 7[337] |
| GAGAGCCTCAGAACCGCATTTTCTGTAACGATCTAAAGTT | 11[345] | 0[329] |
| AAATCCCCGAAACAATTCATGAGGAAGT | 6[321] | 2[308] |
| TACCTAATATCAAAATCATTCAATATTACGTGA | 0[557] | 11[566] |
| GTATACAGGTAATGTGTAGGTAGTCAAATCACCAT | 5[161] | 4[168] |
| AACGTTGTAGAAACAGCGGATAGTTGGGCGGTTGT | 5[77] | 4[84] |
| GTTTATGTCACATGGGAATCCAC | 3[415] | 10[416] |
| ATATTCACAAACAAATTCATATG | 3[392] | 3[414] |
| GACCGGAAGCAATTGCGGGAGAA | 0[202] | 0[180] |
| TCAAGCAGAACCACCACTCACTCAGGTAGCCCGGAATAGG | 7[380] | 8[364] |
| AGCCTCCCCAGGGTCCGGCAAACGCG | 8[88] | 9[104] |
| TTCATTTTCTGCTAAACAACTGAACAACTAAAGGA | 5[329] | 4[336] |
| TCGTTCACCGCCTGGCCCT | 10[331] | 11[344] |
| CGGAAGCACGCAAACTTATTAGCGTT | 8[424] | 9[440] |
| GAGCAAGGTGGCATTTACTCCAACAGGTTCTTTACGTCAACA | 9[189] | 5[202] |
| ATTGCGAATAATGTACAACGGAG | 4[335] | 4[313] |
| CTTTTTTTCGTCTCGTCGCTGGC | 8[111] | 8[89] |
| GACCGTCGAACGGGGAAGCTAATGCAGA | 6[531] | 2[518] |
| GCGTCATACATGCCCTCATAGTT | 0[370] | 0[348] |
| GAAAGTTCAACAATCAGCTTGCTTAGCTTTAATTGTATCG | 4[354] | 7[356] |
| TGTAAATCATGCTCCTTTTGATAATTGCTGAATAT | 5[203] | 4[210] |
| TTCACCTAGCGTGGCGGGTGAAGGGATACCAGTGCATAAAAA | 9[63] | 5[76] |
| ATTTGCCAAGCGGAACTGACCAACGAGTCAATCATAAGGG | 4[312] | 7[314] |
| TAGAACCTACCAGTCTGAGAGAC | 0[580] | 0[558] |
| GGGTTACCTGCAGCCAGCGGTGTTTTT | 4[51] | 4[29] |
| GAATTATCCAATAACGATAGCTTAGATT | 2[559] | 3[559] |
| TTGTCGTCTTTCTACGTAATGCC | 0[328] | 0[306] |
| ACTACTTAGCCGGAACGAGGCGC | 7[273] | 7[295] |
| TTTTTGTCCATCACGCAAATTCCGAGTAAAAGAGTCTTTTTT | 4[702] | 5[702] |
| TTTTTCGGGAGCTAAACAGGTTGTTAGAATCAGAGTTTTT | 0[694] | 1[694] |
| AATCATAATAACCCGGCGTCAAAAATGA | 6[489] | 2[476] |
| AGCAAGCCGTTTAAGAATTGAGT | 4[503] | 4[481] |
| AACAGAGTGCCTGGGGTTTTGCTCACAGAAGGATTAGGAT | 4[396] | 7[398] |
| CCAGCCAAACTTCTGATTGCCGTTTTGGGTAAAGTTAAAC | 4[102] | 7[104] |
| TGAAATTGTTTCAGGGAACTACAACGCC | 6[363] | 2[350] |
| GCCCGCACAGGCGGCCTTTAGTG | 7[63] | 7[85] |
| CAGTAAGAACCTTGAGCCTGTTTAGT | 8[550] | 9[566] |
| ACCAAATTACCAGGTCATAGCCCCGAGTTTTCATCGGCAT | 4[438] | 7[440] |
| TCTTATACTCAGAAAGGCTTTTGATGATATTGACACGCTATT | 9[357] | 5[370] |
| GCCTTATACCCTGTAATACCAATTCTTGCGCTC | 0[179] | 11[188] |
| TTTTTGCGTCCGTGCCTGCATCAGACGTTTTT | 9[25] | 6[21] |
| TTATGGCCTGAGCACCTCAGAGCATAAA | 2[181] | 3[181] |
| CGAGCACAGACTTCAAATACCTCAAAAGCTGCA | 0[221] | 11[230] |

| | | |
|---|---|---|
| GCATCAAAAAGAAGTAAATTGGG | 3[224] | 3[246] |
| TAAGTAGAAGAACTCAAACTATCG | 7[651] | 7[673] |
| ATTTGGCAAATCAACAGTTGAAA | 7[609] | 7[631] |
| GTTGAAACAAACATCAAGAAAAC | 8[615] | 8[593] |
| GAATTGTAGCCAGAATGGATCAGAGCAAATCCT | 0[389] | 11[398] |
| GCTTGACCATTAGATACATTTCG | 8[237] | 8[215] |
| CTGAAAACCTGTTTATCAAACATGTAACGTCAA | 0[515] | 11[524] |
| GACTTTCTCCGTGGCGCGGTTG | 0[76] | 0[54] |
| ACACAACATACGAGGGATGTGGCTATTAATCGGCC | 11[105] | 6[112] |
| TTTTTAACAATATTACCGTCGCTGGTAATATCCAGTTTTT | 6[694] | 7[694] |
| TGCCTGAACAGCAAATGAATGCGCGAACT | 6[657] | 2[644] |
| CAAATATCAAACCAGATGAATAT | 4[629] | 4[607] |
| CAATATGATATTGATGGGCGCAT | 4[167] | 4[145] |
| TTCTGGAATAATCCTGATTTTGCCCGGCCGTAA | 0[599] | 11[608] |
| TTAACAAGAGAATCGATGAACGG | 8[195] | 8[173] |
| GGGCCGGAAGCATAAAGTG | 10[121] | 11[134] |
| GTTTGAGGGGACCTCATTTGCCG | 4[125] | 4[103] |
| GTATTAGAGCCGTCAATAGATAA | 8[657] | 8[635] |
| GCTAATGCCGGAGAGGGTAGCTA | 7[147] | 7[169] |
| TACTTCTTTGATAAAAATCTAAA | 4[671] | 4[649] |
| GAAAGATCGCACTCCAGCCAGCT | 7[105] | 7[127] |
| TCAGGCTGCGCAACTGTTGGGAA | 8[153] | 8[131] |
| ATACCCTTCGTGCCACGCTGAACCTTGCTGAACCT | 5[623] | 4[630] |
| CATAATATTCCGTAATGGGATCCGTGCATCTGCCA | 5[119] | 4[126] |
| TTTTTATCCAATAAATCTCTACCCCGGTAAAACTAGCATG | 7[170] | 8[154] |
| CCGATAATAAAAGGGACTTAACACCGCGAACCACCAGCAG | 11[639] | 0[623] |
| CATCAGCGTCTGGCCTTCCACAGGAACCTGGGG | 0[137] | 11[146] |
| GGAATAACAGAGATAGACATACAAACTTGAGGATTTAGAA | 7[632] | 8[616] |
| CCGGAAGACGTACAGCGCCGCGATTACAATTCC | 0[95] | 11[104] |
| TTCGCGGATTGATTGCTCATTTTTTAAC | 2[139] | 3[139] |
| TAAAGGATTGTATAAGCGCACAAACGACATTAAATGTGAG | 11[135] | 0[119] |
| GATAAAAATTTTTAGCCAGCTTT | 0[160] | 0[138] |
| GATAGTGCAACATGATATTTTTGAATGG | 2[643] | 3[643] |
| GGATAACCTCACAATTTTTGTTA | 3[98] | 3[120] |
| TCAATAATAAAGTGTATCATCATATTCC | 2[601] | 3[601] |
| CAATAGGAACGCAAATTAAGCAA | 3[140] | 3[162] |
| GCGAAAGACGCAAAGCCGCCACGGGAAC | 2[97] | 3[97] |
| TTCCGAATTGTAAACGTGTCGCCAGCATCGGTGCGGGCCT | 7[128] | 8[112] |
| ACATCATTTAAATTGCGTAGAAACAGTACCTTTTA | 5[581] | 4[588] |
| AAGATAAAACAGTTGGATTATAC | 0[622] | 0[600] |
| AACACCCTAAAGGGAGCCC | 10[625] | 11[638] |
| GCATCGAGCCAGATATCTTTAGGACCTGAGGAAGGTTATC | 4[648] | 7[650] |
| CGTAAAGGTCACGAAACCAGGCAATAGCACCGCTTCTGGT | 4[144] | 7[146] |
| CGAGTAACAACCGTTTACCAGTC | 0[118] | 0[96] |

| | | |
|---|---|---|
| GCCTTACGCTGCGCGTAAAATTATTTTTTGACGCTCAATC | 7[674] | 8[658] |
| CCGAACCCCCTAAAACATCGACCAGTTTAGAGC | 0[641] | 11[650] |
| TGCGTACTAATAGTAGTTGAAATGCATATTTCAACGCAAG | 11[177] | 0[161] |
| GATTTTAGACAGGCATTAAAAATA | 0[664] | 0[642] |
| TGATTATCAGATATACGTGGCAC | 3[602] | 3[624] |
| TGGCAAGTTTTTTGGGGTC | 10[583] | 11[596] |
| TCAGCTAACTCACATTAAT | 10[163] | 11[176] |
| CTATTAGTCTTTCGCCGCTACAG | 3[644] | 3[666] |
| AACGCCAAAAGGCGGATGGCTTA | 4[251] | 4[229] |
| AAGAAACAATGACCGGAAACGTC | 4[461] | 4[439] |
| GTACATCGACATCGTTAACGGCA | 4[83] | 4[61] |
| ATACCACCATCAGTGAGGCCAAACCGTTGTAGCAA | 5[665] | 4[672] |
| AACGCCAAAAGGCGGATGGCTTA**-Biotin** | 4[251] | 4[229] |
| AAGAAACAATGACCGGAAACGTC**-Biotin** | 4[461] | 4[439] |
| GTACATCGACATCGTTAACGGCA**-Biotin** | 4[83] | 4[61] |
| ATACCACCATCAGTGAGGCCAAACCGTTGTAGCAA**-Biotin** | 5[665] | 4[672] |

#### References

(1) Wassermann, L. M.; Scheckenbach, M.; Baptist, A. V.; Glembockyte, V.; Heuer-Jungemann, A. Full Site-Specific Addressability in DNA Origami-Templated Silica Nanostructures. *Advanced Materials* **2023**, *35* (23), 2212024. DOI: https://doi.org/10.1002/adma.202212024.

(2) Douglas, S. M.; Marblestone, A. H.; Teerapittayanon, S.; Vazquez, A.; Church, G. M.; Shih, W. M. Rapid prototyping of 3D DNA-origami shapes with caDNAno. *Nucleic Acids Res* **2009**, *37* (15), 5001-5006. DOI: 10.1093/nar/gkp436 From NLM.

(3) Nickels, P. C.; Wünsch, B.; Holzmeister, P.; Bae, W.; Kneer, L. M.; Grohmann, D.; Tinnefeld, P.; Liedl, T. Molecular force spectroscopy with a DNA origami–based nanoscopic force clamp. *Science* **2016**, *354* (6310), 305-307. DOI: 10.1126/science.aah5974.

(4) Nečas, D.; Klapetek, P. Gwyddion: an open-source software for SPM data analysis. *Open Physics* **2012**, *10* (1), 181-188. DOI: doi:10.2478/s11534-011-0096-2 (acccessed 2024-07-23).

(5) Schnitzbauer, J.; Strauss, M. T.; Schlichthaerle, T.; Schueder, F.; Jungmann, R. Super-resolution microscopy with DNA-PAINT. *Nature Protocols* **2017**, *12* (6), 1198-1228. DOI: 10.1038/nprot.2017.024.

(6) Agarwal, N. P.; Matthies, M.; Gür, F. N.; Osada, K.; Schmidt, T. L. Block Copolymer Micellization as a Protection Strategy for DNA Origami. *Angewandte Chemie International Edition* **2017**, *56* (20), 5460-5464. DOI: https://doi.org/10.1002/anie.201608873.

(7) Ponnuswamy, N.; Bastings, M. M. C.; Nathwani, B.; Ryu, J. H.; Chou, L. Y. T.; Vinther, M.; Li, W. A.; Anastassacos, F. M.; Mooney, D. J.; Shih, W. M. Oligolysine-based coating protects DNA nanostructures from low-salt denaturation and nuclease degradation. *Nature Communications* **2017**, *8* (1), 15654. DOI: 10.1038/ncomms15654.

(8) Jing, X.; Zhang, F.; Pan, M.; Dai, X.; Li, J.; Wang, L.; Liu, X.; Yan, H.; Fan, C. Solidifying framework nucleic acids with silica. *Nature Protocols* **2019**, *14* (8), 2416-2436. DOI: 10.1038/s41596-019-0184-0.

(9) Cordes, T.; Vogelsang, J.; Tinnefeld, P. On the mechanism of Trolox as antiblinking and antibleaching reagent. *J Am Chem Soc* **2009**, *131* (14), 5018-5019. DOI: 10.1021/ja809117z From NLM.

(10) Zähringer, J.; Cole, F.; Bohlen, J.; Steiner, F.; Kamińska, I.; Tinnefeld, P. Combining pMINFLUX, graphene energy transfer and DNA-PAINT for nanometer precise 3D super-resolution microscopy. *Light: Science & Applications* **2023**, *12* (1), 70. DOI: 10.1038/s41377-023-01111-8.

(11) Schröder, T.; Scheible, M. B.; Steiner, F.; Vogelsang, J.; Tinnefeld, P. Interchromophoric Interactions Determine the Maximum Brightness Density in DNA Origami Structures. *Nano Letters* **2019**, *19* (2), 1275-1281. DOI: 10.1021/acs.nanolett.8b04845.

(12) Widengren, J.; Schwille, P. Characterization of Photoinduced Isomerization and Back-Isomerization of the Cyanine Dye Cy5 by Fluorescence Correlation Spectroscopy. *The Journal of Physical Chemistry A* **2000**, *104* (27), 6416-6428. DOI: 10.1021/jp000059s.

(13) Kitamura, A.; Tornmalm, J.; Demirbay, B.; Piguet, J.; Kinjo, M.; Widengren, J. Trans-cis isomerization kinetics of cyanine dyes reports on the folding states of exogeneous RNA G-quadruplexes in live cells. *Nucleic Acids Research* **2023**, *51* (5), e27-e27. DOI: 10.1093/nar/gkac1255 (acccessed 10/8/2024).
